# FuzzyClusTeR: a web server for analysis of tandem and diffuse DNA repeat clusters with application to telomeric-like repeats

**DOI:** 10.64898/2026.03.19.712643

**Authors:** Anna Y. Aksenova, Anna S. Zhuk, Artem G. Lada, Alexei V. Sergeev, Kirill V. Volkov, Arsen Batagov

**Affiliations:** Laboratory of Amyloid Biology, St. Petersburg State University, 199034 St. Petersburg, Russia; Matheomics, Skolkovo Innovation Center, Moscow, 143026 Russia; Institute of Applied Computer Science, ITMO University, 197101 St. Petersburg, Russia; Department of Microbiology and Molecular Genetics, University of California, Davis, CA 95616, USA; Research Park, St. Petersburg State University, 199034 St. Petersburg, Russia; Center for Molecular and Cell Technologies, Research Park, St. Petersburg State University, 199034 St. Petersburg, Russia; Mesh Bio Pte. Ltd, 10 Anson Road, #22-02, Singapore

## Abstract

DNA repeats constitute a large fraction of eukaryotic genomes and play important roles in genome stability and evolution. While tandem repeats such as microsatellites have been extensively studied, the genomic organization and potential functions of dispersed or loosely organized repeat patterns remain poorly understood. Here we present *FuzzyClusTeR*, a web server for the identification, visualization and enrichment analysis of DNA repeat clusters in genomic sequences. Using parameterized metrics, *FuzzyClusTeR* detects both classical tandem repeats and regions where related motifs occur in proximity without forming perfect tandem arrays, which we term *diffuse (or fuzzy) repeat clusters*. The server supports analysis of user-defined sequences as well as genome-scale datasets, including the T2T-CHM13 and GRCh38 human genome assemblies, and provides interactive visualization and statistical tools for assessing the genomic distribution of repetitive motifs and corresponding clusters. As a demonstration, we analyzed telomeric-like repeats in the T2T-CHM13v2.0 genome and identified families of diffuse clusters enriched in these motifs. Comparison with simulated sequences suggests that these clusters represent non-random genomic patterns with potential evolutionary and functional significance. *FuzzyClusTeR* enables systematic exploration of repeat clustering across genomic regions or entire genomes. It is available at https://utils.researchpark.ru/bio/fuzzycluster

**GRAPHICAL ABSTRACT:** 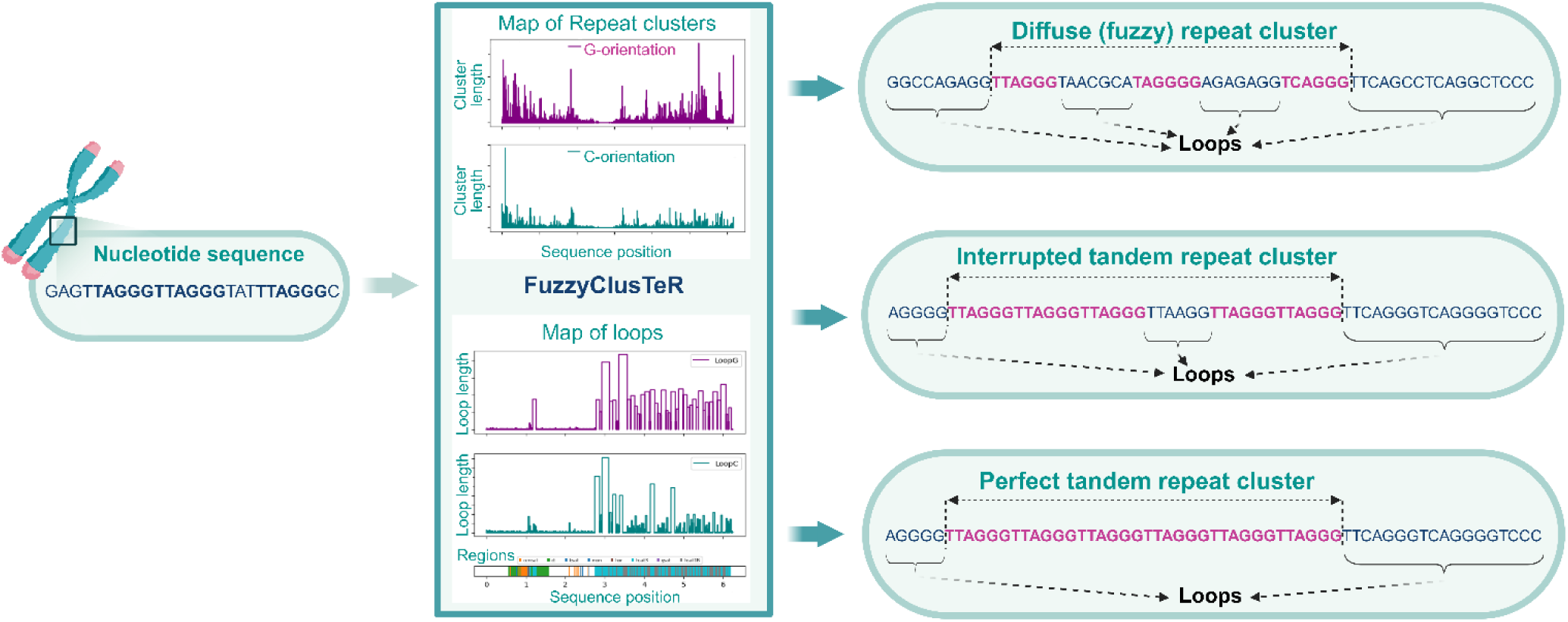

## INTRODUCTION

Ends of linear chromosomes in most eukaryotic species are composed of specific DNA repeats known as telomeric repeats. Together with protein complexes and RNAs, they form special protective structures called telomeres that fulfill several important functions: preventing chromosome ends from being misrecognized as double-strand breaks (DSBs) (1–4) and precluding progressive DNA loss during replication (5–7). The replication and repair machinery is strictly regulated at telomeres to avoid inappropriate activation of DNA damage response (DDR) and to prevent end-to-end chromosome fusions. Unsurprisingly, telomere structure and length are important factors regulating cell proliferation potential (8). Telomere length can be maintained by telomerase or via telomerase-independent mechanisms, such as Alternative Telomere Lengthening (ALT) (7, 9–14).

Telomeric repeats in all eukaryotic kingdoms share similarity and appear to originate from the same ancestral grounding telomere repeat. In most eukaryotes they are represented by short (5–8 bp) species-specific simple tandem repeats composed of TA/T-rich and G-rich parts that are able to form G-quadruplexes (15–17). The most widespread telomeric motif is the hexanucleotide TTAGGG repeat motif which is characteristic for all Metazoans and is also found in some invertebrates, protozoa and fungi (15, 18, 19). Many lineages of insects and non-insect arthropods carry pentanucleotide TTAGG telomeric repeats (19, 20). The typical plant telomeric motif is TTTAGGG and was originally described in *Arabidopsis thaliana* (21, 22). However, plants demonstrate significant variability in telomere composition (23). Telomeric repeats described in protozoan species include such examples as TTGGGG (in *Tetrahymena* and *Glaucoma*) or TTTTGGGG (*Oxytricha* and *Euplotes*) (16).

Variations in telomeric motif sequence have been described in a single organism. A notable example of such diversity is *Homo sapiens*, where besides the canonical TTAGGG, different variant repeats at telomeres have been reported. TTGGGG and TGAGGG along with the TTAGGG motif, have been described at human telomeres by Allshire *et al.* who used hybridization with specific oligonucleotide probes (24). TGAGGG and TCAGGG repeats were observed by Telomere Variant Repeat PCR (25). Other studies described CTAGGG repeats associated with а high rate of telomere instability in the male germline (26, 27). More variant types have been observed by NGS and FISH in two studies by Conomos *et al.* (28) and Lee *et al.* (29). They included TCAGGG, TTCGGG, GTAGGG, TGAGGG, TTGGGG, TAAGGG, ATAGGG, CTAGGG, TTTGGG, TTAAGGG and other variants with the more common ones being TCAGGG and TGAGGG. Interestingly, the percentage of different variant repeats depended on the cell-type (e.g. ALT *vs.* Telomerase-activated) (28–30). Grigorev *et al.* described some additional variants at telomeres, including TTAGGGG, TTAGG, TTGGG, TGGG and others (31). Collectively, these studies highlight the problem of telomere heterogeneity and its functional implications. Some of these telomeric variant repeats can perform specialized functions. For instance, variant TCAGGG repeats intersperse canonical telomeric repeats at ALT telomeres and serve as binding sites for orphan nuclear receptors of the NR2C/F classes which promote spatial proximity of telomeres and telomere-telomere recombination (28, 32, 33). The appearance of these repeats was assigned to DNA Pol eta which can alleviate the replication stress at telomeres (34–36). CTAGGG repeats compared to TTAGGG canonical variants bind better to POT1, which is a component of telomere-protecting Shelterin complex (26).

Telomeric repeats are not found exclusively at the ends of chromosomes. They are present in multiple internal sites of chromosomes in many species (17). Such sequences are called Interstitial Telomeric Sequences (ITSs) and are found in most vertebrates, including humans, and in some plants (21, 37–50). Commonly, ITSs are divided into two primary classes: heterochromatic ITSs (het-ITSs) represent large blocks of telomeric repeats that are present mainly in centromeric or pericentromeric regions, and short ITSs (s-ITSs) representing stretches of a limited number of TTAGGG hexamers. The human genome lacks large blocks of het-ITSs. The relatively few large ITSs identified in the human genome seem to originate from terminal fusion of ancestor chromosomes that gave rise to modern human chromosomes (40, 49, 51, 52). Human s-ITS were described in several studies and mostly are composed of 2 to 25 copies of perfect TTAGGG repeats, such sequences are found in all human chromosomes (40, 42, 43, 47, 49, 53–57).

The functions of such sequences in the genome remain enigmatic. Multiple studies suggest that they can bind different Shelterin components, such as RAP1, TRF1, TRF2 and TIN2 (52, 54, 58–61). ITSs have been implicated in genome structure organization and interstitial telomeric loop formation stabilized by TRF2 binding and TRF2-interacting Lamin A/C (55, 62). This conclusion stems from the observations that T-loops can be formed at interstitial telomeric sequences (interstitial telomeric loops, ITL) and that they are stabilized by TRF2 binding and TRF2-interacting lamin A/C. ITSs are unstable and can directly provoke different genome instability events including chromosome rearrangements and mutagenesis (63–65). Moreover, in ALT tumor cells such sequences can be inserted into different chromosomal positions through a targeted telomere insertion (TTI) mechanism mediated by the NR2C/F orphan nuclear receptors (33). Sieverling *et al.* analyzed whole-genome sequencing data from over 2500 matched tumor-control samples spanning 36 different tumor types and highlighted the pertinence of ITSs in tumorigenesis, revealing frequent telomere sequence insertions in tumors with ATRX or DAXX mutations (66).

The gapless telomere-to-telomere assembly of the human genome provides an unprecedented opportunity to analyze the entire genome and to evaluate the non-coding regions which were masked in the previous genome versions (67–69). Our analysis of the T2T-CHM13v2.0 human genome revealed numerous clusters of telomeric-like repeats in addition to canonical TTAGGG ITSs. These clusters are widespread across human chromosomes and represent different combinations of telomeric-like repeat variants separated by intervals of variable length. We refer to such regions as *diffuse (or fuzzy) repeat clusters*. To systematically characterize these patterns, we applied parameterized scoring metrics based on cluster length, repeat count, and the genomic distribution of repeat intervals. In parallel, we developed *FuzzyClusTeR,* a web-based tool that enables identification, visualization, and annotation of repeat clusters across selected genomic regions or entire genomes. Importantly, FuzzyClusTeR can be used to analyze clusters formed by a wide range of repetitive DNA motifs, including micro- and minisatellites. The server supports detection of both classical tandem repeats and diffuse repeat clusters under user-defined parameters and can facilitate the identification and annotation of such repeat clusters in genomes from diverse species. This approach enables systematic investigation of repeat clustering patterns and their potential roles in genome organization.

## MATERIAL AND METHODS

### Genome sequences and search patterns

We used the T2T-CHM13v2.0 human genome assembly in our analysis (67). This reference genome assembly is virtually gapless and represents repetitive and/or heterochromatic regions with a high degree of fidelity.

As a query we used two different patterns as determined by regex expressions:

(1) Canonical telomeric hexamer: TTAGGG (and its reverse complement CCCTAA),
(2) FuzzyTel: T{1, 2}A{0, 1}G{3, 5}|T{1, 2}\D{1}A{1}G{3, 5}|T{2}A{1}G{2} (and its reverse complement C{3, 5}T{0, 1}A{1, 2}|C{3, 5}T{1}\D{1}A{1, 2}|C{2}T{1}A{2})

Here, the pattern N{n, m} defines letter N repeated from n to m times. For example, G{3, 5} stands for either GGG, GGGG, or GGGGG. Additionally, the “|” character defines the logical operator “OR”, and “\D” defines any letter (non-digit). Standard nucleic acid code is typically used in the pattern matching; U is treated as equivalent to T. In addition, sequences containing degenerate IUPAC patterns, such as NRYBDKMHVSW can be submitted for the analysis – such regions are considered for loop-length determination but are not used for pattern matching and will be marked as red lines on the plots.

FuzzyTel pattern allows us to search for the most frequently occurring variants of telomeric repeats that can be obtained from the canonical TTAGGG repeat mostly by single nucleotide substitution or insertion/deletion according to the reported variants (see Figure 1). It was complemented by the TGGG motif and thus comprises telomeric repeat variants reported in (31, 66). The complete list of all the motifs included in the FuzzyTel pattern is provided in Table S1. The number of hits found for the canonical and FuzzyTel patterns, as well as the median loop lengths, are provided in Tables S2 and S3.

**Figure 1.**
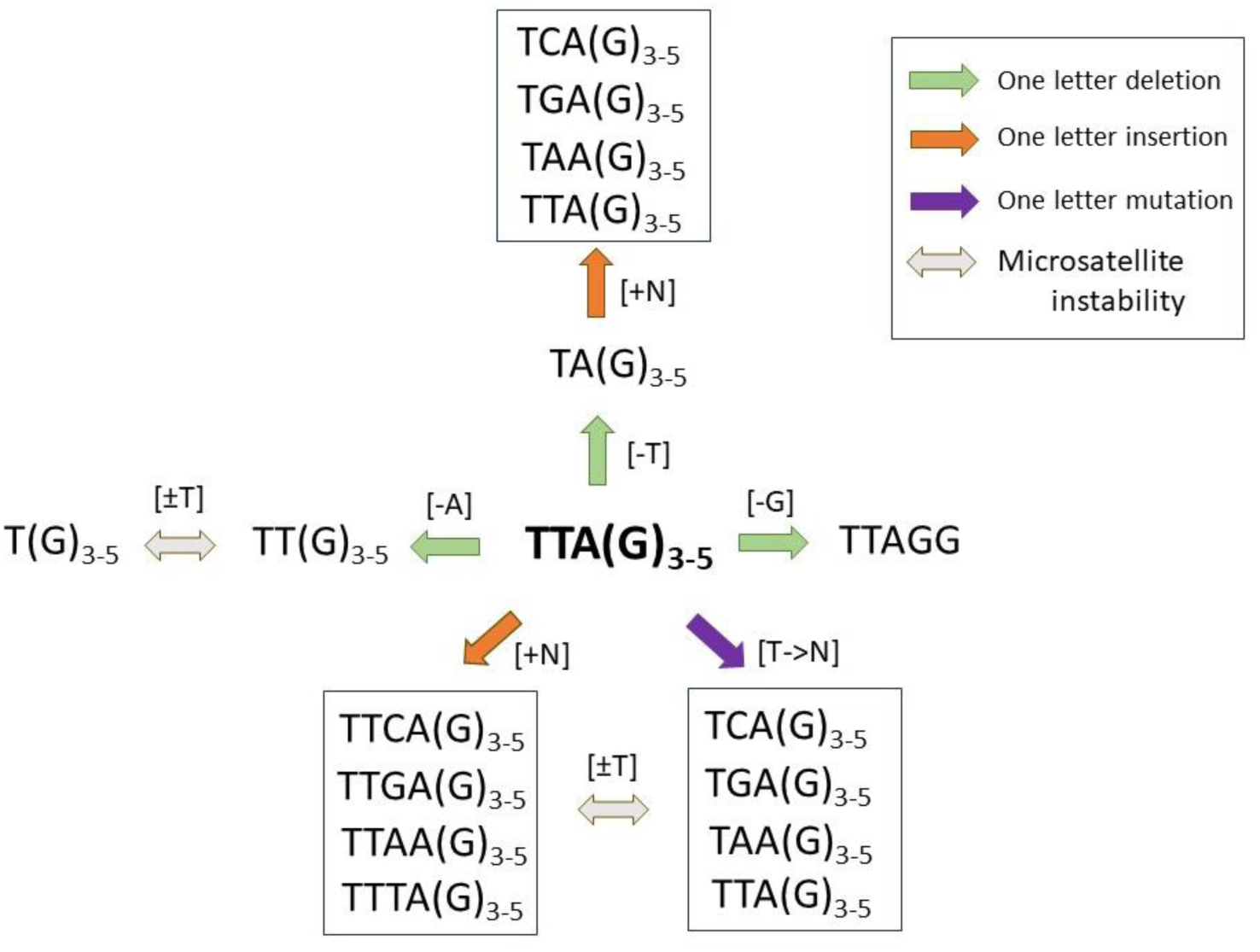
Pathways showing relationship between telomeric-like repeats and possible mutation changes which can convert some variants of the repeats into the others.

The artificial genome was generated using Python NumPy random.choice function (https://numpy.org/doc/2.1/reference/random/generated/numpy.random.choice.html) and MT19937, which implements the Mersenne Twister algorithm for pseudorandom number generation (70). Frequencies of A, T, G and C letters and chromosome sizes were calculated for each chromosome of the T2T-CHM13v2.0 human genome and passed as the p and the size arguments to np.random.choice function. Chromosomes were assembled as SeqRecord objects in the final form using the Bio.SeqRecord module of BioPython (https://biopython.org/). The artificial genome does not contain telomeres and serves for the purpose of demonstrating a reference random distribution of the canonical telomeric and telomeric-like repeats in a uniformly random nucleotide sequence of genomic length and the nucleotide frequency of the natural chromosomes.

### Algorithm for clustering and filtration of repeats

All studied patterns were searched using regular expressions and their locations in the T2T-CHM13v2.0 human genome were catalogued. We then computed the intervening intervals (referred to as ’loops’) between the identified locations of two sequentially mapped motifs. In cases where a loop length did not exceed a threshold, it was extended to merge with the succeeding motif. By default, the threshold for the loop length in our algorithm is taken as the median of the loop length distribution in the analyzed sequence. To identify and catalogue clusters of telomeric and telomeric-like repeats, we used а robust upper cutoff of the median, i.e. the median plus the standard deviation. For the FuzzyTel pattern such a cutoff was established as 90 nt and for the canonical telomeric repeats as 3660 nt, based on the human genome statistics. This methodology allowed us to derive clusters of conjoined intervals and individual telomeric-like repeats (those not incorporated into clusters). A schematic illustration of a diffuse cluster is presented in Figure S1. In all cases the cluster contains at least two repeats determined by the pattern.

The obtained dataset was then characterized and filtered using a parameter which is designated as the Cluster Score (CS). This parameter signifies the density of telomeric-like repeats within a cluster of specified length and is obtained by the following expression:

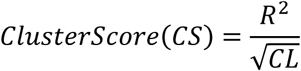

where R – is the number of repeats in a cluster,

CL - is the cluster length (nt).

The behavior of the algorithm for cluster identification is contingent on the CS and the loop length (both can be specified in FuzzyClusTeR). Besides that, we introduced a parameter designated as the Score Significance Ratio (SSR), reflecting the ratio between the Theoretical Cluster Score (TCS) and the observed CS for a given number of repeats (R). SSR is a measure of cluster significance.

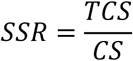

where the TCS is determined as:

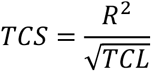

Thus,

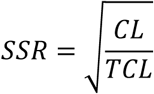

Theoretical Cluster Length (TCL) is determined based on the information about the median (or user-determined) loop length and the observed number repeats in a cluster:

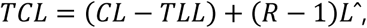

where R – is a number of repeats in a cluster i,

CL - the cluster length,

*L̂* – is a median (or user-defined) loop length,

TLL – total loop length, is the sum of all loops in a cluster of given length

The quadratic dependence on R can be interpreted as “amplified repeat density”. Unlike the linear R/CL ratio, it emphasizes the contribution of repeat number: as R increases, the score grows faster than linearly, reflecting the idea that clusters with a large number of repeats receive an enhanced contribution. Clusters with many repeats are more biologically relevant than those with only a few. Using √CL in the denominator, rather than CL, makes the metric more sensitive to the repeat density and prevents long but densely packed clusters from being underestimated, since length grows faster than the number of repeats. The resulting expression can serve as an equivalent of signal-to-noise ratio, since squaring inflates the contribution of monitored events (signal), while √CL serves as a normalizing scale. Overall, this formulation enables the comparison of clusters of different sizes without introducing a strong bias toward either short or long clusters. The combined implementation of CS and SSR allows enrichment of repeat clusters from the analyzed sequences by virtue of user-provided thresholds for these parameters. This allows selection and visualization of any repeat clusters for their subsequent study without the need for a preliminary hypothesis about their distribution.

### Simulation of theoretical distributions and their fitting

We used scipy.stats.gamma and scipy.stats.pareto (https://docs.scipy.org/doc/scipy/reference/stats.html) to simulate the theoretical Gamma and Pareto probability density functions (pdfs). Fitting of the CS data to the theoretical models was performed with scipy.optimize.curve_fit (https://docs.scipy.org/doc/scipy/reference/generated/scipy.optimize.curve_fit.html). The best results with respect to the normalized RMSE and QQ-plot were obtained for the Gamma pdf (shape≈0.7, scale≈0.2) for the CS distribution obtained from the pseudo genome for the canonical telomeric repeats. Similar results were obtained for the natural genome with the exception of the high CS values observed at the tail – this part of the distribution was fitted best with the Pareto (shape≈1.5, scale≈1.0) model. In the case of the clusters formed by the FuzzyTel pattern we obtained the best fit with the Gamma pdf (shape≈0.5, scale≈0.6) for the CS of the pseudo genome and with the Pareto pdf (shape≈1.7, scale≈1.0) for the natural T2T-CHM13v2.0 human genome. The QQ-plots built with the help of scipy.stats.probplot indicated the distribution type, as well as its parameters, shape and scale. RMSE was calculated for all the cases and normalized RMSE (NRMSE) was defined as RMSE adjusted by the cluster count. Note that in most cases a mixed Gamma-Pareto distribution model could fit the empirical distribution.

## RESULTS

### Analysis of repetitive pattern distribution using FuzzyClusTeR

Interspersed DNA repeats can be found at multiple locations in the human genome, but their significance and systematic occurrence have not been studied in details. In this study, we present a web-based tool, FuzzyClusTeR, and an algorithm for analysis and visualization of nucleotide sequence repeats that can be organized in tandem or *diffuse (or fuzzy) clusters*. We define *diffuse repeat clusters* as genomic regions in which related sequence motifs occur within a limited genomic neighborhood, but do not form continuous tandem arrays. Such clusters may contain varying number of matched patterns that occur in proximity but are separated by variable spacers and therefore do not form canonical tandem repeats. FuzzyClusTeR is freely available at https://utils.researchpark.ru/bio/fuzzycluster and can currently be used to find and investigate clusters formed by micro- or minisatellite repeat units.

The distribution of certain patterns or repetitive sequences in the genome can be gauged by examining the intervals (further referred to as ’loops’) between these repeats. The information about the median loop length across the analyzed sequence (such as human genome or individual chromosomes) is calculated automatically by the FuzzyClusTeR program and is used in our algorithm as a threshold for joining the intervals and subsequent identification of repeat cluster boundaries. The loop length can also be specified arbitrarily by the user and may take any value reflecting the expected distribution of repeats. Thus, for instance, in case of tandem repeats, it should be set to zero. Using the specified loop length threshold, our algorithm merges any predefined repeat with preceding loops of the threshold length or shorter into a single cluster.

FuzzyClusTeR supports the analysis of FASTA and multi-FASTA sequence files, as well as nucleic acid sequences that can be entered manually. Additionally, human genomes assemblies GRCh38 and T2T-CHM13v2.0 are preloaded and can be used for the analysis. FuzzyClusTeR performs separate analysis for the two sequence orientations, forward and reverse complement. Currently, there are two pre-determined patterns that can be chosen by the user: canonical human telomeric and telomeric-like FuzzyTel. Other repetitive patterns can be entered by the user manually as a single motif or multiple motifs (complex pattern). In the case of a complex pattern, the median loop length will be influenced by the most frequently occurring motif, so we recommend estimating a single pattern occurrence first to avoid a significant bias. The tool shows the position of loops between detected pattern repeats across the analyzed sequence (“Map of Loops” figure), analyzes the loop length distribution, including the median loop, the interquartile range (IQR), and the maximal loop length (“Loop length distribution” figure) and provides the information about found clusters (“Map of repeat clusters” figure and in table format). All the found individual motifs and their indexes can be downloaded in the form of csv files. The start and the end position of the identified repeat clusters along with their characteristics such as cluster length, the number of repeats in the cluster, the CS and the SSR scores, and the total length of all loops per cluster are also provided in the csv files for the user’s further evaluation and analysis. Additionally, there is a print option to display all pattern occurrences and repeat clusters in the sequence as text, which can be activated for short sequences.

Refinement of the repeat cluster search is achieved using the CS and the SSR cluster scores, as described in Material and Methods. The CS parameter reflects the density of repeats in an interval of given length (cluster). This parameter can be provided by the user and is used as a cut-off by the program to show clusters with the score equal to or higher than indicated. SSR parameter is another metric introduced here. It estimates the significance of аn observed cluster relative to its statistical expectation. SSR is calculated as the square root of the ratio between the observed cluster length (CL) and its theoretically expected estimate (TCL). Here, TCL is defined as the expected length of the cluster under the assumption that the motifs inside it are equally spaced. In this formula, the spacing (L^) is either a genome- and motif-specific constant (median loop length), or is user-defined. As follows, from the formula, SSR is scaled between 0 and 1. The lower the SSR, the more distant is the cluster from the TCL and, consequently, the more significant it is. Like the CS, this parameter can be provided by the user for selecting clusters above the specified significance threshold. The user can also set the minimum number of repeats to be considered in a cluster. FuzzyClusTeR also supports the analysis of perfect tandem repeats. When this option is enabled, the algorithm joins only repeats separated by zero-length intervals into singular clusters and report only these clusters. If а degenerate pattern is entered, these tandem repeats can represent any of the motifs that are described by the pattern.

### Distribution of telomeric-like repeats and their clusters in the human genome

We performed genome-wide analysis of the intervals between telomeric-like repeats in the T2T-CHM13v2.0 version of the human genome using (i) the canonical telomeric pattern, and (ii) the FuzzyTel telomeric-like pattern, which can be derived from the perfect telomeric hexamer by limited number of changes (see Material and Methods and Figure 1). The map of the loops generated by these patterns are presented in Figures S2 and S3. Remarkably, some genomic regions contained very long sequence stretches lacking any telomeric-like repeats. Such regions were observed in all chromosomes, and they specifically overlapped with some centromeric repeats (e.g. High Order Repeats, HOR regions) in most of the chromosomes. Interestingly, while in some chromosomes this behavior was observed in both G-rich and C-rich orientations, many others demonstrated remarkable asymmetry (Figures S2 and S3).

The canonical telomeric repeats are spaced at a median interval of approximately 3330-3350 nucleotides, whereas this interval reduced to about 80 nucleotides for the FuzzyTel pattern (see Figure S4, Table S3 and Supplementary files 1 and 2). The loop length distribution showed that most intervals across all the patterns are short, and the frequency of loops longer than 200 nucleotides was steadily declined (Figure 2). To estimate to which extent this distribution can be imitated by a pseudorandom process, we generated an artificial genome (hereafter also termed pseudo genome) as a uniformly random nucleotide sequence, and estimated loop distribution in it. As shown in Figure 2, short and very long loops are more characteristic of the natural genome, compared to the artificial one. This distribution accounts for the high natural abundance of the telomeric and telomeric-like repeats at telomeres and sub-telomeres, where they are either spaced at short or zero-length intervals and long regions, such as certain centromeric regions depleted of these sequences.

**Figure 2.**
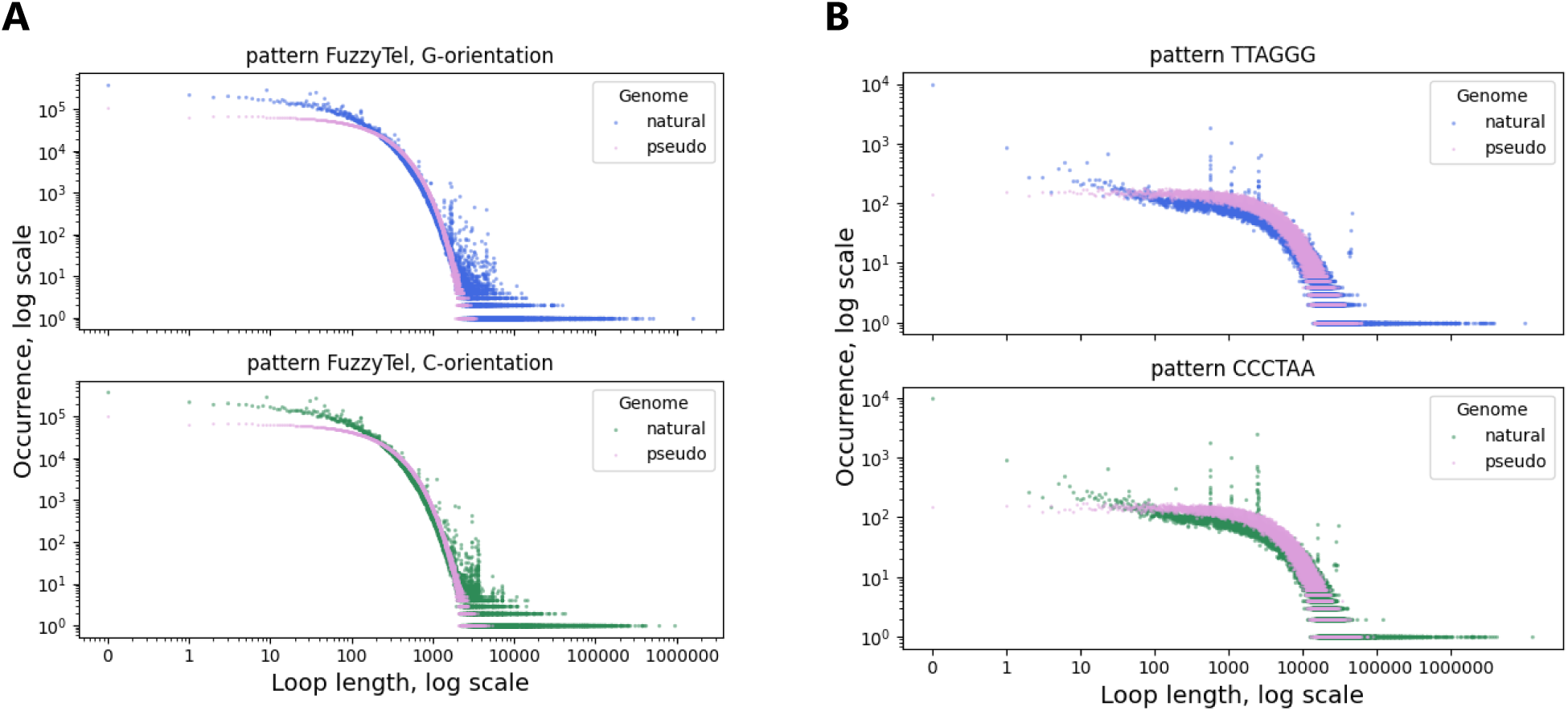
Loop length distribution in the genome for (A) Loops generated by the FuzzyTel pattern, (B) Loops generated by the canonical telomeric pattern. G-rich and C-rich orientations are shown separately. The T2T-CHM13v2.0 human genome (natural) and artificial genome (pseudo) were analyzed. The data is plotted in the double decimal logarithmic scale.

As described in the Material and Methods, we used 90 nucleotides as a threshold loop length for the purpose of joining intervals in a cluster for the FuzzyTel pattern, then ran the algorithm and identified the FuzzyTel clusters, i.e. genomic regions enriched in such motifs. For the canonical telomeric pattern the loop length threshold was set to 3660. The CS characteristic shows the repeat density in a cluster of a certain length and varies from a fraction of one to tens of thousands, depending on the cluster length and number of repeats. Interestingly, pseudo random distribution produced only a few clusters with CS more than 10 for the FuzzyTel pattern and more than 1.2 for the canonical telomeric pattern (see Figure 3). At the same time, many clusters with higher CS values were observed in the human T2T-CHM13v2.0 genome. This argues in favor of a mechanism that allowed the generation and propagation of such clusters during evolution of the human genome.

**Figure 3.**
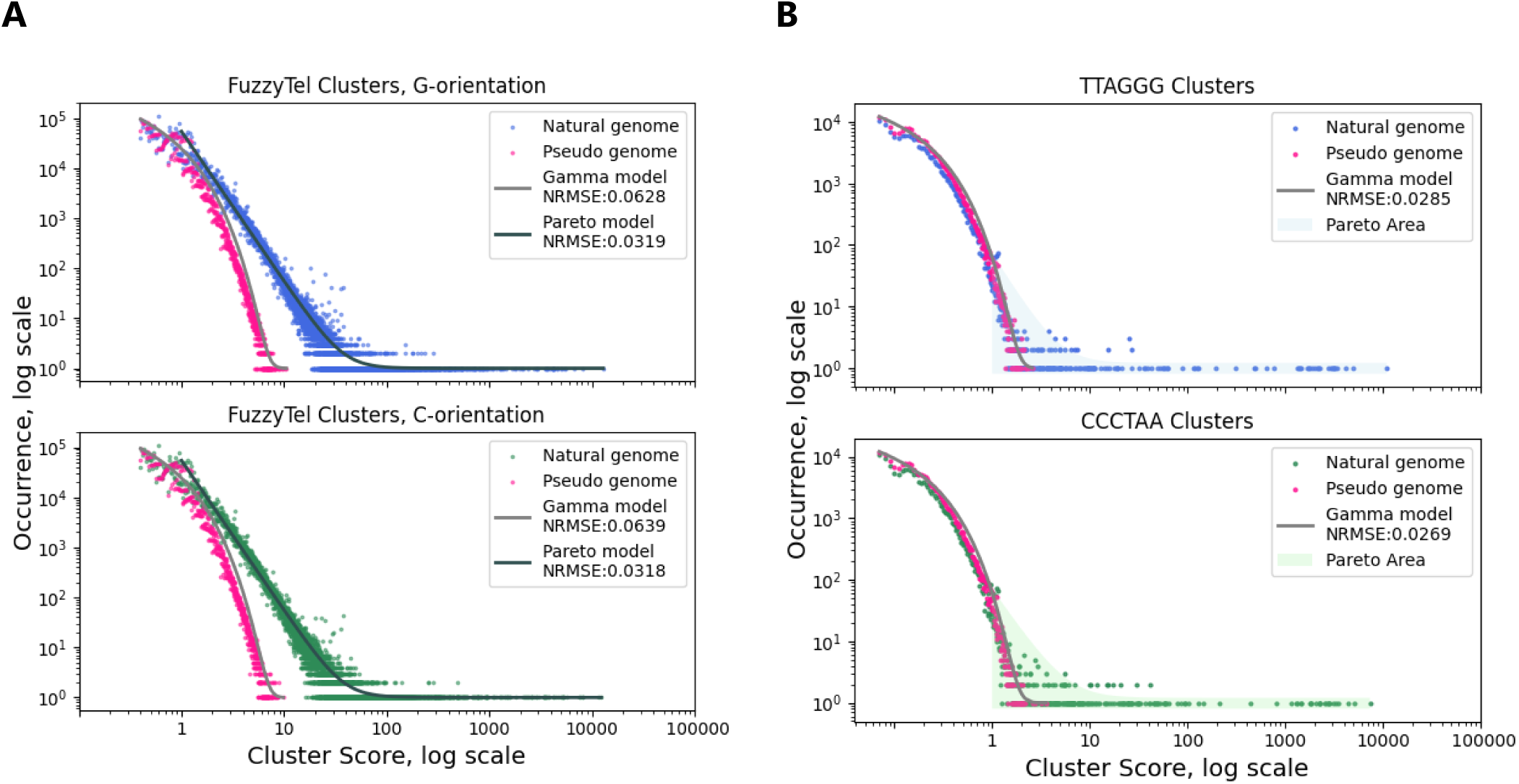
Cluster Score (CS) distribution in the genome for (A) FuzzyTel pattern and (B) Canonical telomeric pattern. G-rich and C-rich orientations are shown separately. The T2T-CHM13v2.0 human genome (natural) and artificial genome (pseudo) were analyzed. The data was fitted with Gamma and Pareto distributions and is plotted in the double decimal logarithmic scale.

More specifically, the observed CS frequency followed a Gamma-like distribution in the case of the pseudo genome, which is consistent with the hypothesis of their random occurrences. At the same time, for the CS obtained from the human T2T-CHM13v2.0 genome, we observed a Pareto-like heavy tailed distribution for both the canonical telomeric and FuzzyTel patterns (see Figure 3, Figure S5). As expected, the Pareto tail included telomeres, which naturally represent the longest clusters of telomeric repeats in the genome and also included many clusters with non-telomeric localization. To additionally assess the significance of observed clusters, we calculated the SSR ratio for the canonical telomeric and FuzzyTel patterns. Decreasing SSR values correspond to shorter loops within the clusters and increased repeat density relative to their theoretical expectations (TCS). Figures S6 and S7 demonstrate this relationship. In these plots clusters, represented as points, demonstrate the tendency to progressively deviate from the theoretically expected threshold line as their computed SSR values decrease (see Figures S6 and S7).

To visualize the most significant clusters, we plotted CS versus SSR for all the clusters found for the FuzzyTel and the canonical telomeric patterns in the T2T-CHM13v2.0 genome and performed the same analysis for the artificial genome. As shown in Figure 4, while some clusters can be reproduced by а pseudo-random process, many lie in the plot far outside the area of pseudo-random clusters. For the vast majority of the FuzzyTel clusters, SSR is decreased with the growth of CS. For the canonical telomeric repeats, we observed two types of clusters: clusters where CS was negatively correlated with SSR and clusters where these metrics behaved independently. Thus, the distribution of the canonical telomeric motif clusters was bimodal. In general, both the number of repeats per cluster and the cluster length (to a lesser degree), correlated positively with CS (Figures S8-S11).

**Figure 4.**
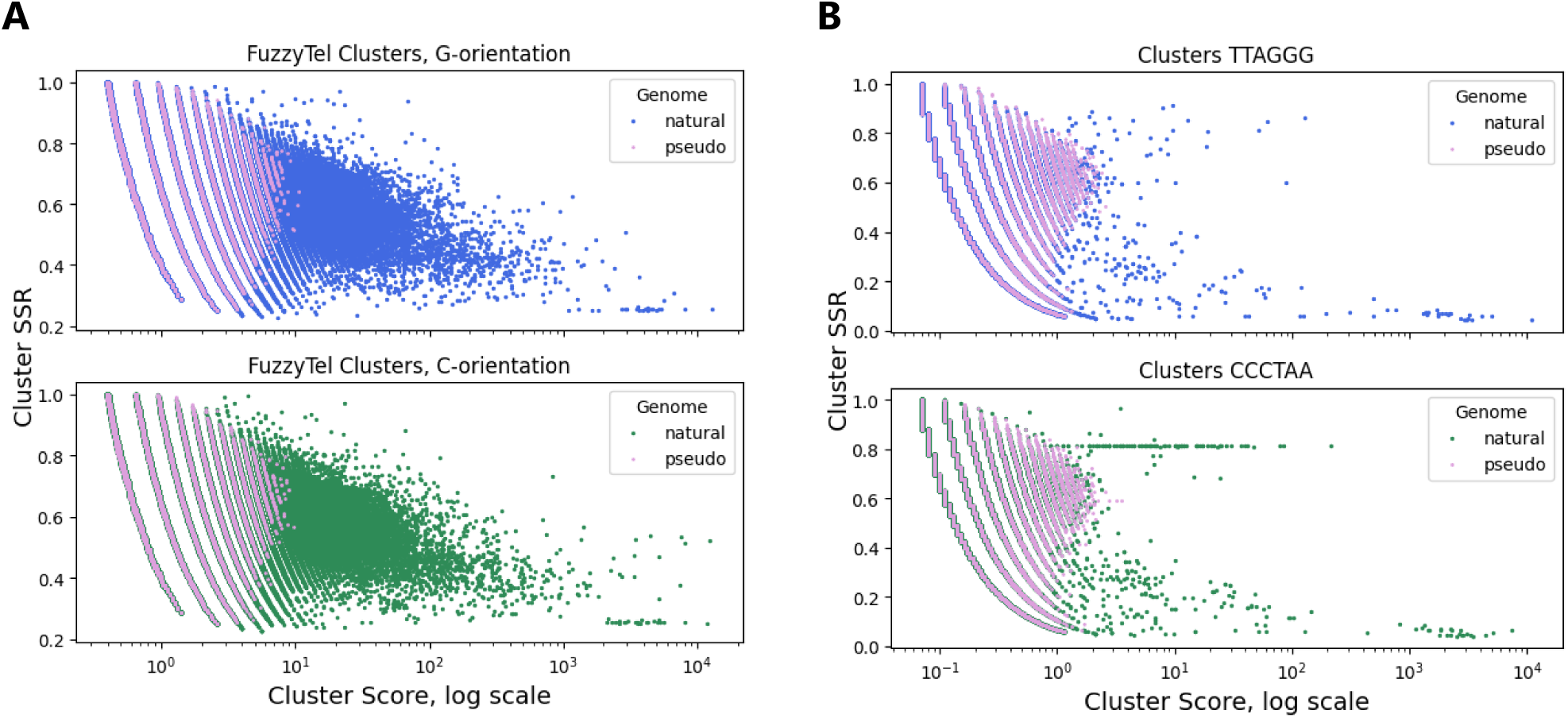
CS (Cluster Score) vs. SSR calculated for the T2T-CHM13v2.0 human genome (natural) and artificial genome (pseudo). (A) FuzzyTel pattern, (B) Canonical telomeric pattern.

To examine the clusters, which strongly deviate from the random model, we performed cluster enrichment at SSR ≤ 0.5 and CS ≥ 1.2 for the canonical telomeric pattern (Figure 5 and Supplementary files 3-5) and SSR ≤ 0.5 and CS ≥ 10 for the FuzzyTel pattern (Figure 6 and Supplementary files 6-8). As shown in Figure 4, very few clusters are detected in these regions in the pseudo genome. Thus, the selected parameters enrich for significant clusters, including telomeric regions where a high density of repeats is expected. In total, we selected 242 G-rich and 275 C-rich clusters matching the canonical telomeric pattern in telomeric and non-telomeric regions, with the total number of repeat units per cluster ranging from 3 to 937 (Supplementary Files 4–5). For the FuzzyTel pattern, we detected 3494 G-rich and 3359 C-rich clusters, with repeat unit counts ranging from 8 to 1015 (Supplementary Files 7–8).

**Figure 5.**
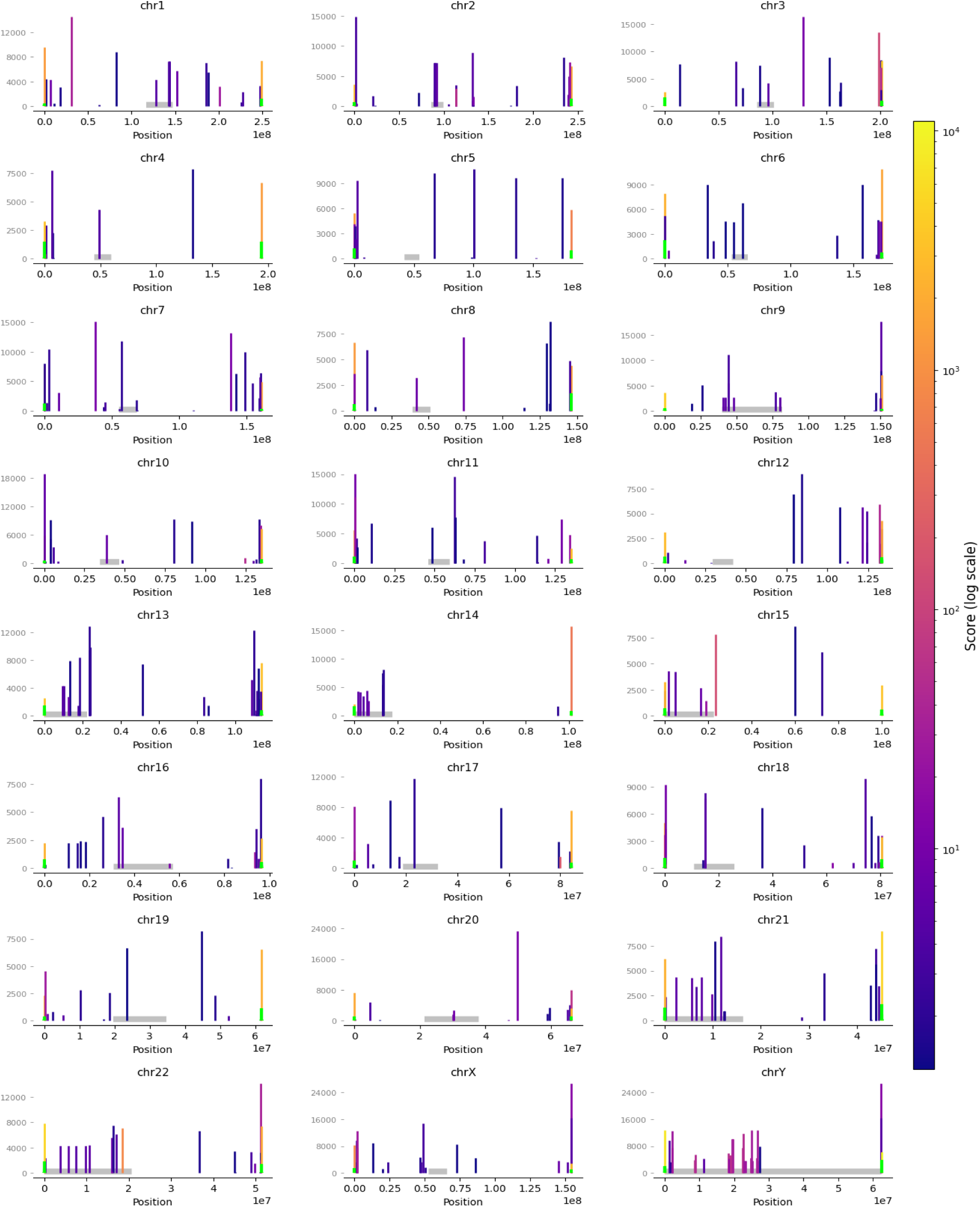
Clusters formed by the canonical telomeric pattern (TTAGGG or CCCTAA) across human chromosomes. Cluster score thresholds of CS ≥1.2 and SSR ≤ 0.5 were used for enrichment. Cluster length is represented along the vertical axes and the color reflects the log-scaled CS score; centromeric areas are grey, clusters with tandem canonical telomeric repeats (1 nt loops are allowed at most) are lime green.

**Figure 6.**
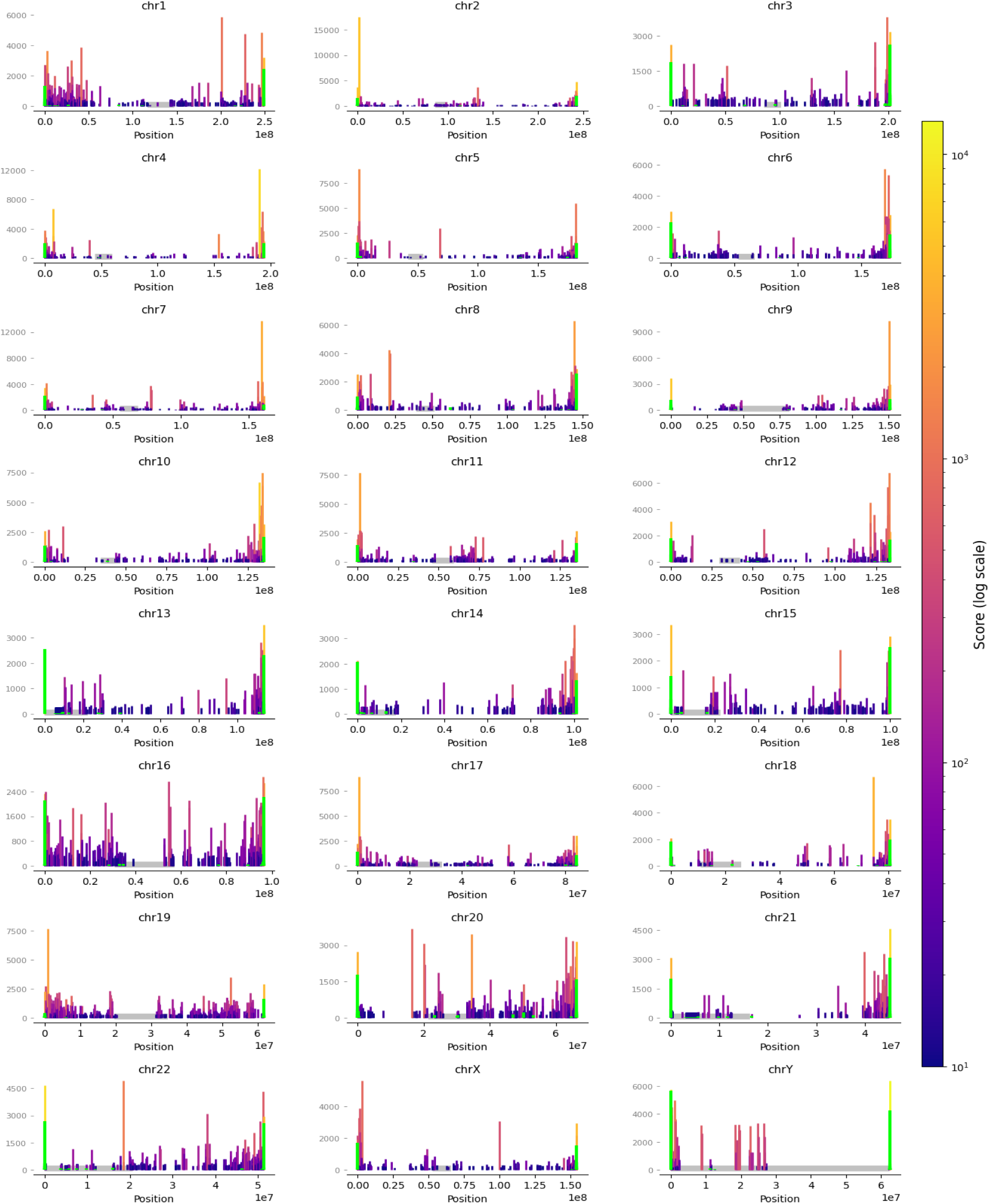
Clusters formed by the FuzzyTel pattern (G- and C-orientations) across human chromosomes. Cluster score thresholds of CS ≥10 and SSR ≤ 0.5 were used for enrichment. Cluster length is represented along the vertical axes and the color reflects the log-scaled CS score; centromeric areas are grey, clusters with tandem FuzzyTel repeats (1 nt loops are allowed at most) are lime green.

The identified clusters with non-telomeric localization can be divided into several groups: (1) diffuse clusters, in which motifs do not form uninterrupted tandem arrays - these clusters display considerable diversity in their organization and repeat density and often appear highly irregular; (2) clusters exhibiting regular loop spacing, correlating with the presence of large tandem repeat structures, such as minisatellites; (3) clusters with tandem or nearly tandem organization. Examples of some of these clusters are shown in Figures S12 and S13. The functional significance of diffuse clusters containing telomeric motifs remains unclear, and their potential role in genome organization will require further investigation.

Another interesting group consists of the so-called sparse clusters, in contrast to the dense clusters described above. In these regions, telomeric motifs are separated by large spacers and form long arrays. We identified sparse clusters containing canonical telomeric repeats with SSR > 0.5 and CS ≥ 1.2, some of which exhibit regular loop spacing indicating presence of larger DNA repeats (Figures S8, S9, S14 and S15). Notably, both sparse and dense clusters were detected in centromeric regions, most frequently overlapping with centromeric transition zones (Figures S16 and S17).

The human genome does not contain very long ITSs compared to other species; however, even short ITSs may have functional roles, for example by serving as binding sites for the Shelterin complex. We analyzed the distribution of tandem and nearly tandem canonical telomeric repeats (loop length ≤ 1) across all chromosomes of the CHM13v2.0 genome and observed an asymmetry in the occurrence of longer clusters between G-rich and C-rich orientations. In the standard reference genome representation, clusters in the C-rich orientation were more frequently detected (Figure S18).

## DISCUSSION

Multiple tools have been developed for identifying motifs and short tandem repeats within genomes. Motif Scraper was designed to identify specific motifs within FASTA files (71). Written in Python, this tool can be easily installed via the pip package manager and does not require substantial computing resources. Motif Scraper functions like a FASTA sequence pattern search tool and offers a user-friendly way to view and analyze data by providing results in a CSV format with coordinates, the start and end points of motifs, details about the DNA strands, and motif locations. Another tool, PERF, facilitates finding repeats both *de novo* and based on pre-defined motifs (72). PERF too was written in Python and enables rapid identification of short tandem repeats. To discover motifs using PERF, the list of repetitive elements must be structured in a specific format, rather than provided as a simple list of motifs. The *de novo* approach generates a list of repeats by considering specific parameters: the minimum and maximum lengths of the repeating motif, the cutoff for the minimum length of the repeat and the minimal or the maximal sequence length required for analysis. The output of PERF is displayed as a tab-delimited BED file. PERF is not intended for precise repeat searches, because it reports overlapping repeats, repeats in the middle of the motifs and partial repeats. Another approach for identifying *de novo* tandem repeats is provided by the Dot2dot algorithm (73). A variety of options can be adjusted in the Dot2dot for detecting repeats, such as minimal and maximal motif length, number of gaps allowed in the motifs, maximal insert size allowed between the motifs in the repeats. Dot2dot is implemented in C and is capable of generating output in either BED or DOT formats. Several other algorithms are available for the detection and *de novo* discovery of tandem repeats, including Tandem Repeats Finder (TRF) with a web version (74), Kmer-SSR (75), MREPS (76), MISA-web (77), and the Generic Repeat Finder (GRF) (76) for identifying terminal direct and inverted repeats. The number of repeats within genomes identified by different tools can vary due to the differences in the algorithm design, parameters used, and varying sensitivity to different types of repeats. Most of the existing algorithms are intended to detect repetitive patterns or find occurrences of specific motifs in the sequence and do not estimate the distribution of the studied patterns.

FuzzyClusTeR relies on a regular expression (regex) based string search and, instead of identifying *de novo* repeats, is designed for identification of pre-determined patterns (simple or complex), analysis of motif distribution along the provided linear character string and enrichment of regions, where the examined pattern occurs more frequently. FuzzyClusTeR is focused on analysis of distances between motif occurrences in the sequence, is adaptive to the user tasks and is capable of finding (i) all tandem repeats and various clusters of tandem repeats within the sequence, (ii) diffuse repeat clusters which may have different numbers of repeats separated by loops of varying lengths, (iii) clusters formed by larger DNA repeats, which contain matched patterns.

All clusters are scored based on the number of repeats and the length of the clusters, whose significance is estimated by the specific scoring metrics, CS and SSR. This parametrized approach allows the user to carry out enrichment analysis and visualization of virtually any repeat clusters for their subsequent study. Additionally, since FuzzyClusTeR is able to capture generic pattern distribution, this tool also facilitates studying sequence regularity.

Many studies have focused on the prevalence and genomic distribution of tandem repeats. For example, a recent comparative analysis of repeat lengths across genomes reported a widespread occurrence of long repetitive tracts and their stable distribution across mammals (78). In contrast, FuzzyClusTeR addresses a different aspect of genome organization by examining repeat clustering patterns and identifying genomic regions enriched in specific motifs. In addition to classical tandem arrays, genomes may contain dispersed or loosely organized repeat patterns, in which related motifs occur in proximity without forming continuous arrays. We refer to such regions as diffuse or fuzzy repeat clusters, representing a distinct form of genomic organization. Analysis of these patterns provides new opportunities for exploring repeat dynamics and genome structure. Applying this approach to the canonical telomeric and variant telomeric-like FuzzyTel patterns in the T2T-CHM13v2.0 human genome assembly allowed us to identify clusters enriched in these motifs and to characterize their genomic abundance.

We found that both short and long loops (or intervals) between telomeric motifs are more characteristic of а natural genome, compared to the pseudo genome. For instance, we observed that all types of telomeric-like repeats are exceptionally rare within certain centromeric regions. However, all chromosomes contain clusters of these repeats of varying lengths and significance. Our findings reveal that diffuse clusters of telomeric-like repeats are distributed along chromosomal arms, where they are interspersed and co-localized with genes, other repeat classes, and biologically significant sequences. The observed distribution of these clusters suggests their potential origin in various tandem repeats that expand through recombination-based mechanisms or DNA polymerase slippage during replication cycles. In some cases, telomeric-like motifs represent small cores in arrays of larger divergent satellite repeats that accumulated many changes, but retained intact telomeric units. This observation may suggest that longer repetitive sequences, while expanding and diverging during evolution, could keep telomeric motifs intact due to the selective pressure, while other sequences might be lost or mutated. In other cases, no larger repetitive units could be identified: such clusters enriched with telomeric-like motifs, had highly irregular organization patterns which may have resulted from telomeric-motif insertions or be remnants of highly mutated and evolved ITSs.

Telomeric repeat arrays can form secondary structures, such as G-quadruplexes, and promote the formation of R-loops, features that have been suggested to influence chromatin organization and gene regulation. Insertion of telomeric repeats into transcriptionally active regions may promote the formation or stabilization of R-loops, which in turn can contribute to various forms of genomic instability (79). A recent study of 205 human ITSs in U2OS, HeLa and HEK293 cell lines showed that these regions can be transcribed and contribute to TERRA pool (80). Because R-loop targeting is being explored as a potential therapeutic strategy in several malignancies (81), systematic characterization of telomeric repeat clusters - including both tandem and diffuse clusters - may provide additional insights into genomic regions prone to R-loop formation. In this context, genome sequencing of cancer patients could help identify somatic variants or population polymorphisms associated with such clusters. The T2T genome assembly represents a relatively recent resource, and, as additional datasets, including those of gene expression and protein interactions, become available, further analyses will help clarify the potential functional roles of the non-canonical ITSs identified in this study.

Future studies could build on these insights in several ways: (i) incorporating advanced algorithms for detailed characterization of repeat sequences and diffuse clusters, enabling deeper genomic analysis; (ii) investigating statistical models of repeat and cluster length distributions to uncover the biological mechanisms underlying their formation and expansion; (iii) applying the analysis presented here to a variety of biological species may lead to new insights into the mechanisms of evolution and its connection with genome stability and gene regulation.

In conclusion, we have developed a new web-based bioinformatics tool FuzzyClusTeR that allows the study of repeat distribution in nucleotide sequences and region enrichment analysis. This tool offers a user-friendly interface and helps to identify all occurrences of a specific nucleotide motif, to evaluate spacing intervals, to select the most significant diffuse or tandem repeat clusters, and to explore organization of larger repeats and repeat-containing sequences. By enabling systematic identification and visualization of both tandem and diffuse repeat clusters, FuzzyClusTeR provides a flexible framework for studying repeat organization across various genomes. We have applied FuzzyClusTeR to the study of telomeric-like repeats in the human genome. Our results suggest that telomeric-like repetitive motifs may frequently form genomic clusters, which lack strict tandem arrangement. Such diffuse clustering patterns may represent an additional layer of genome organization that has remained largely unexplored. The discovery of such clusters in the T2T-CHM13 genome illustrates the potential of this approach to reveal previously unrecognized patterns of repeat distribution. We anticipate that FuzzyClusTeR will facilitate further investigations into the evolutionary origins, functional roles and genomic dynamics of repetitive motifs. FuzzyClusTeR can be applied to investigation of structural features of genomic sequences. Its output can be used in a broad range of evolutionary and functional genomic studies. It enables systematic analysis of how the statistical properties of selected nucleotide patterns vary across sequences and organisms. Organisms with distinct repetitive sequence architectures can be directly compared using the proposed sequence-filtering strategies, which establish a consistent analytical basis within FuzzyClusTeR. Filtering based on clustering metrics supports unbiased assessment of associations between nucleotide patterns, structural elements and gene functions. Furthermore, fitting the resulting sets of genomic regions to a range of statistical distributions facilitates the evaluation of alternative models and potential drivers of sequence evolution.

## DATA AVAILABILITY

FuzzyClusTeR web server is freely available at https://utils.researchpark.ru/bio/fuzzycluster

## ACKNOWLEDGEMENTS

The research was concluded with the use of RRC “Center for Molecular and Cell Technologies, ” Scientific Park, St. Petersburg State University (St. Petersburg. Russia). We thank Mikhail Korman for critical reading of the manuscript and Mikhail Langovoy for discussion.

## AUTHOR CONTRIBUTIONS

Anna Y. Aksenova: Conceptualization, Formal analysis, Visualization, Methodology, Validation, Writing—original draft, review & editing, Algorithm creation, Software development. Anna S. Zhuk: Formal analysis, Visualization, Writing—review & editing. Artem G. Lada: Formal analysis, Methodology. Writing— review & editing. Alexei V. Sergeev: Methodology, Software development. Kirill V. Volkov: Formal analysis, Methodology, Writing—review & editing. Arsen Batagov: Formal analysis, Methodology, Validation, Writing—original draft, review & editing, Software development.

## SUPPLEMENTARY FIGURES AND TABLES

### Supplementary Tables

**Table S1.**
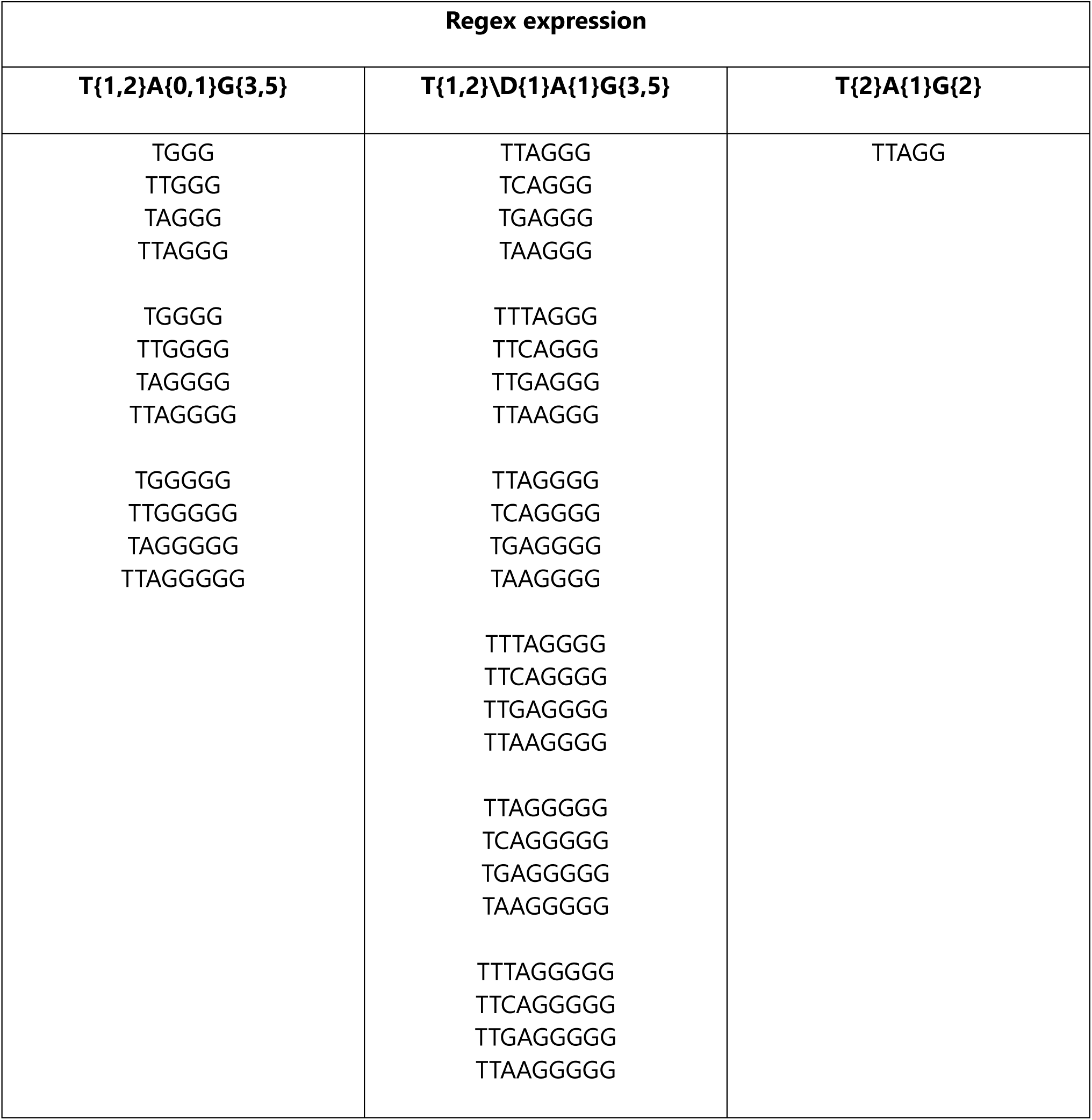
Specific nucleotide sequences determined by the FuzzyTel regex expressions (shown for G-rich orientation only).

**Table S2.** Counts of telomeric-like repeats by chromosome in the human T2T-CHM13v2.0 genome as determined by different patterns.

|  | Canonical pattern |  | FuzzyTel pattern |  |
| --- | --- | --- | --- | --- |
|  | Orientation |  | Orientation |  |
|  | TTAGGG | CCCTAA | G-rich | C-rich |
| **Chr1** | 44203 | 43871 | 1747173 | 1748604 |
| **Chr2** | 46191 | 46483 | 1677351 | 1680378 |
| **Chr3** | 38465 | 37964 | 1354091 | 1371488 |
| **Chr4** | 36173 | 35361 | 1211308 | 1181556 |
| **Chr5** | 34824 | 34193 | 1210256 | 1207979 |
| **Chr6** | 32836 | 32593 | 1144278 | 1146116 |
| **Chr7** | 29136 | 29510 | 1109338 | 1115714 |
| **Chr8** | 28579 | 27941 | 1001303 | 1002686 |
| **Chr9** | 23723 | 24109 | 912878 | 914484 |
| **Chr10** | 25236 | 25519 | 979862 | 977446 |
| **Chr11** | 25864 | 25618 | 994334 | 991520 |
| **Chr12** | 24899 | 25246 | 941685 | 945071 |
| **Chr13** | 19309 | 18959 | 680174 | 713817 |
| **Chr14** | 18157 | 17891 | 708709 | 696660 |
| **Chr15** | 16801 | 16671 | 678971 | 680437 |
| **Chr16** | 15467 | 15104 | 705189 | 706412 |
| **Chr17** | 14694 | 14516 | 701968 | 707373 |
| **Chr18** | 14812 | 15133 | 521479 | 527237 |
| **Chr19** | 9403 | 9483 | 570928 | 556204 |
| **Chr20** | 13000 | 11651 | 527813 | 523139 |
| **Chr21** | 7868 | 7790 | 299630 | 305598 |
| **Chr22** | 7931 | 7915 | 416114 | 424261 |
| **ChrX** | 29453 | 29211 | 1043407 | 1033766 |
| **ChrY** | 6580 | 12280 | 285947 | 244765 |
| **Total** | 563604 | 565012 | 21424186 | 21402711 |

**Table S3.** Loop length medians between telomeric repeats determined by different patterns in the T2T-CHM13v2.0 genome.

|  | **Canonical** |  | **FuzzyTel** |  |
| --- | --- | --- | --- | --- |
|  | **Orientation** |  | **Orientation** |  |
|  | **TTAGGG** | **CCCTAA** | **G-rich** | **C-rich** |
| **Chr1** | 3375.0 | 3418.0 | 76.0 | 76.0 |
| **Chr2** | 3376.0 | 3368.0 | 82.0 | 82.0 |
| **Chr3** | 3314.0 | 3355.0 | 84.0 | 84.0 |
| **Chr4** | 3399.0 | 3478.5 | 88.0 | 91.0 |
| **Chr5** | 3351.0 | 3421.0 | 85.0 | 85.0 |
| **Chr6** | 3360.0 | 3412.5 | 85.0 | 86.0 |
| **Chr7** | 3513.0 | 3446.0 | 80.0 | 79.0 |
| **Chr8** | 3248.5 | 3370.0 | 83.0 | 83.0 |
| **Chr9** | 3478.0 | 3384.0 | 80.0 | 81.0 |
| **Chr10** | 3404.0 | 3355.5 | 77.0 | 77.0 |
| **Chr11** | 3312.0 | 3321.0 | 75.0 | 76.0 |
| **Chr12** | 3372.0 | 3400.0 | 80.0 | 80.0 |
| **Chr13** | 3395.5 | 3626.0 | 93.0 | 90.0 |
| **Chr14** | 3377.0 | 3370.0 | 79.0 | 79.0 |
| **Chr15** | 3253.5 | 3318.0 | 77.0 | 77.0 |
| **Chr16** | 3707.0 | 3463.0 | 70.0 | 72.0 |
| **Chr17** | 3487.0 | 3548.0 | 67.0 | 67.0 |
| **Chr18** | 3297.0 | 3168.0 | 86.0 | 89.0 |
| **Chr19** | 3889.0 | 3747.5 | 62.0 | 60.0 |
| **Chr20** | 2905.0 | 3320.5 | 70.0 | 71.0 |
| **Chr21** | 3065.0 | 3248.0 | 79.0 | 78.0 |
| **Chr22** | 3085.5 | 3148.5 | 64.0 | 65.0 |
| **ChrX** | 3239.0 | 3319.0 | 84.0 | 85.0 |
| **ChrY** | 2322.0 | 2413.0 | 98.0 | 100.0 |
| **Median** | 3366.0 | 3370.0 | 80.0 | 79.5 |
| **Standard deviation** | 281.5 | 232.4 | 8.4 | 8.7 |
| **Median (whole genome)** | 3348 | 3330.50 | 79 | 79 |

### Supplementary Figures

**Figure S1.**
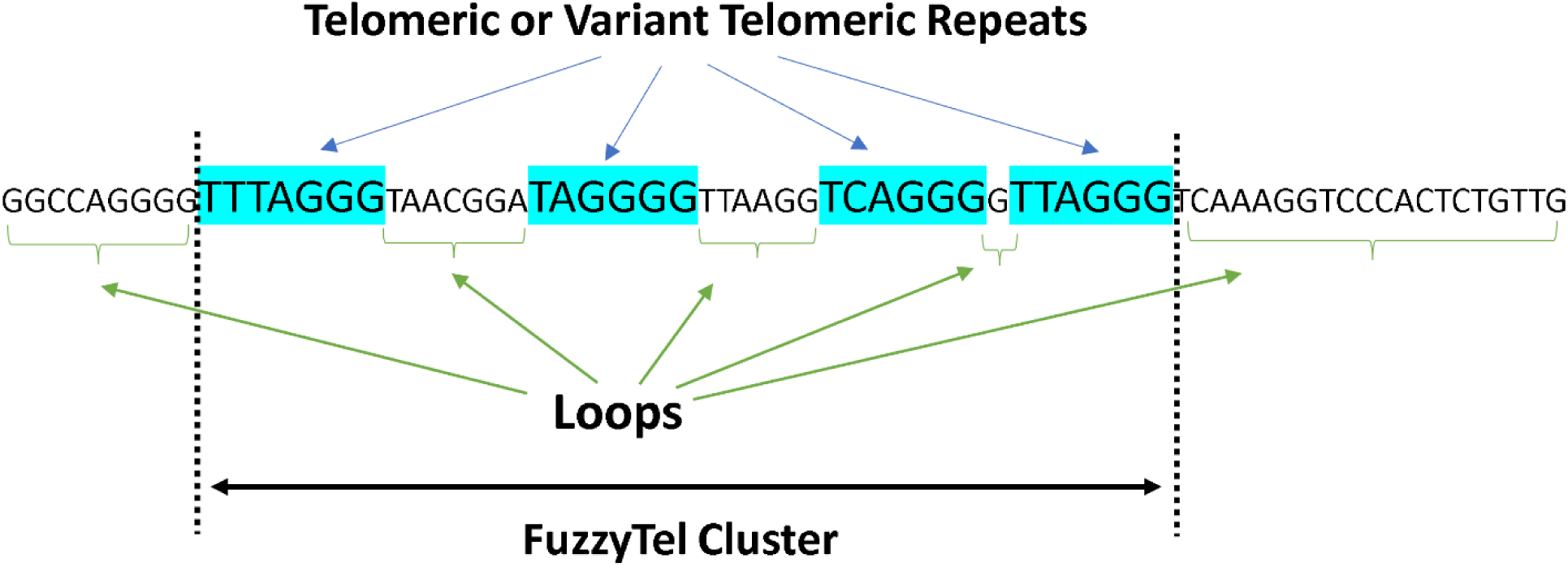
Schematic illustration of diffuse repeat cluster represented by the FuzzyTel pattern, G-rich orientation is shown.

**Figure S2.**
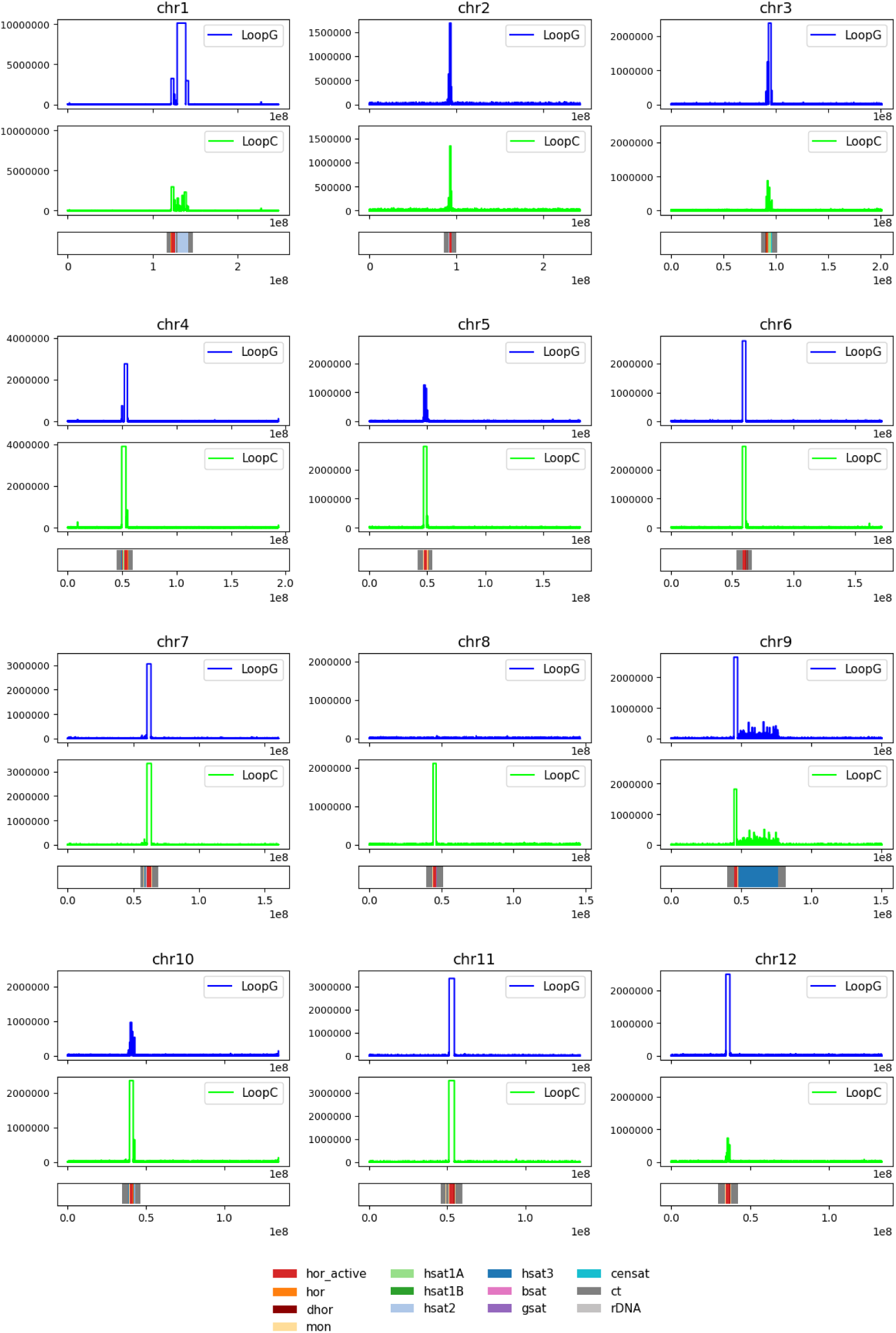
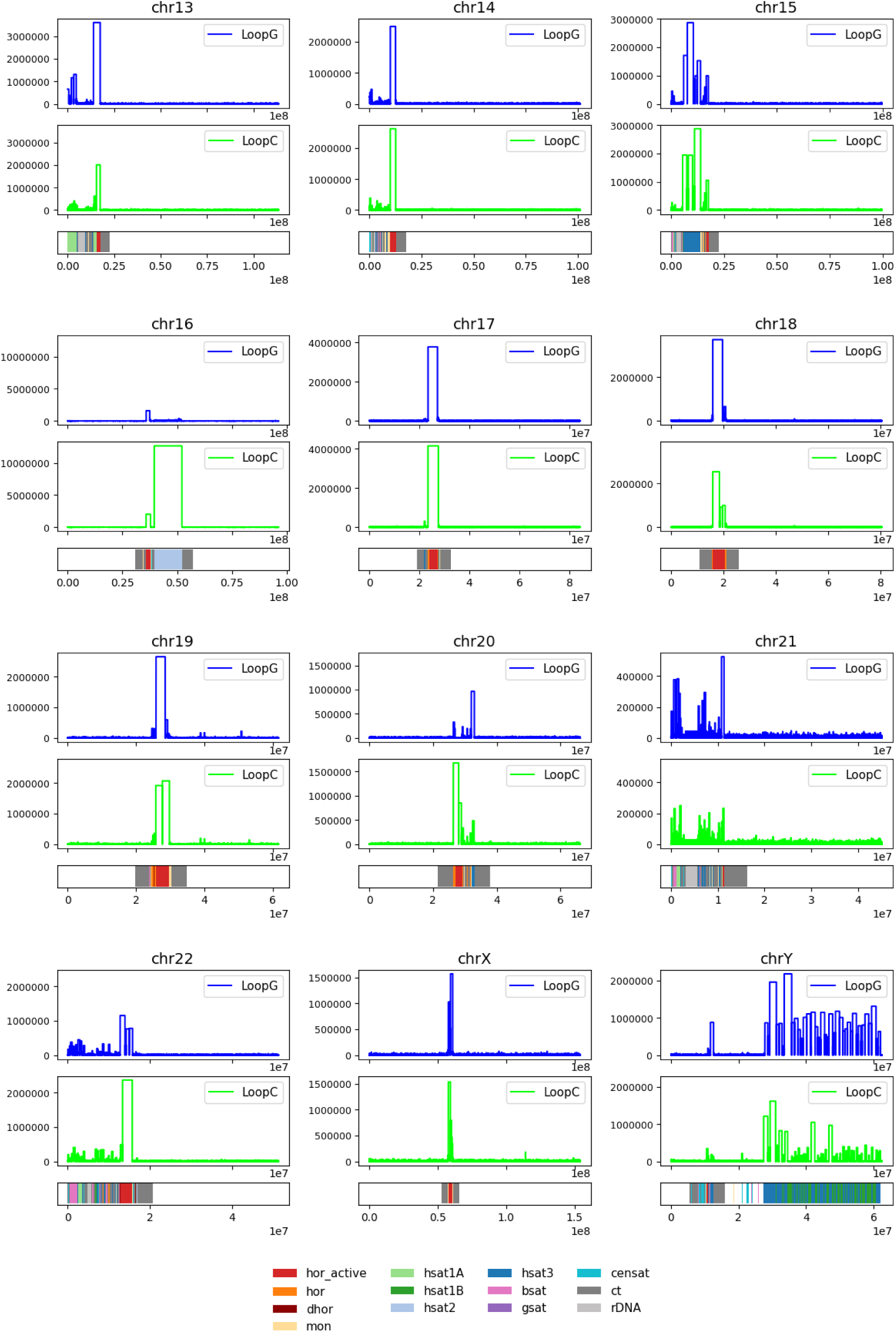
Position of large loops lacking canonical telomeric motif in all chromosomes of the human T2T-CHM13v2.0 genome. LoopG – matched pattern TTAGGG, LoopC – matched pattern CCCTAA

**Figure S3.**
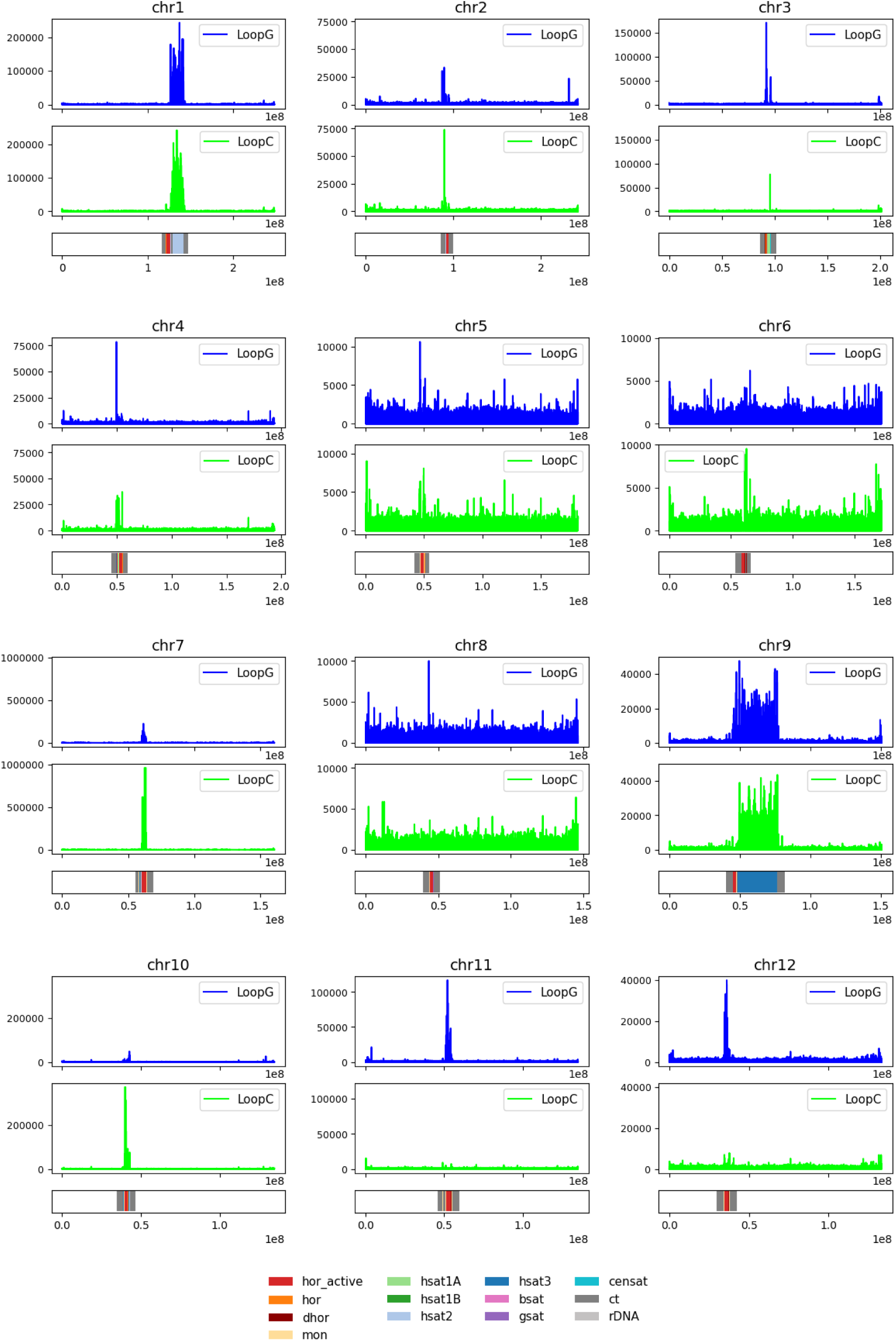
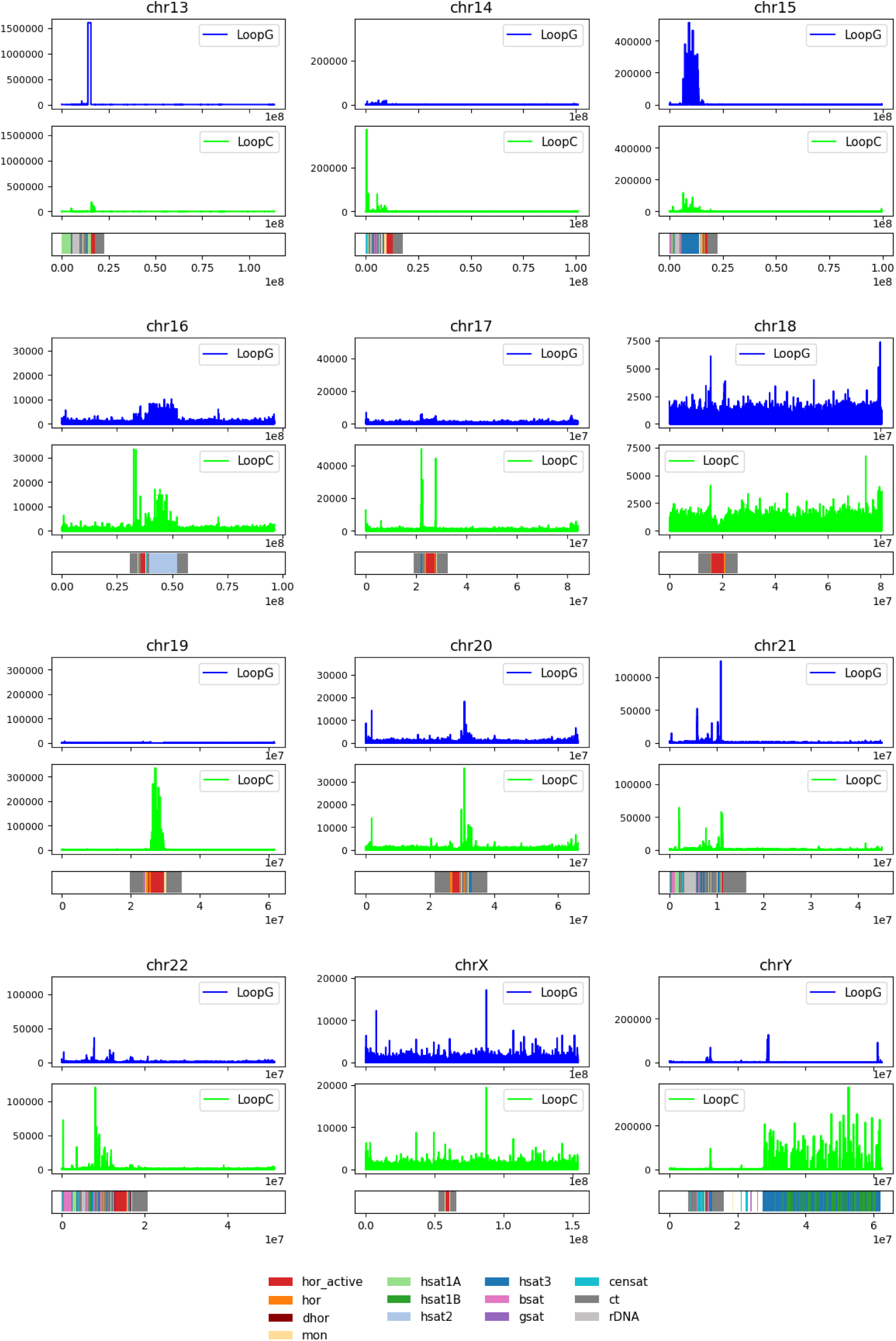
Position of large loops lacking canonical telomeric motif in all chromosomes of the human T2T-CHM13v2.0 genome. LoopG – matched pattern FuzzyTel G-rich orientation, LoopC – matched pattern FuzzyTel C-rich orientation

**Figure S4.**
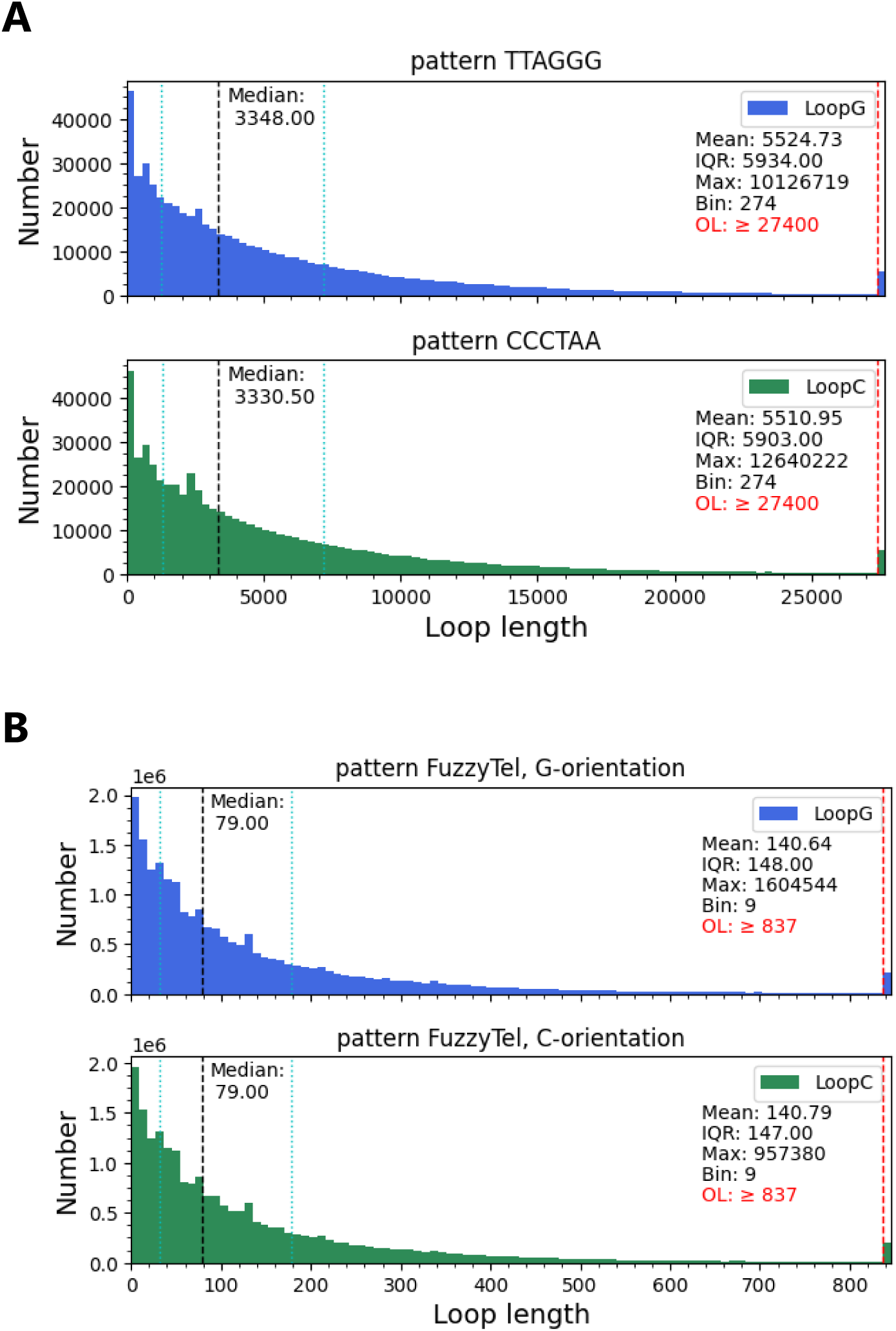
Loop lengths distribution between telomeric-like repeats. (A) Distribution of loop lengths determined by the canonical telomeric pattern. (B) Distribution of loop lengths determined by FuzzyTel pattern. Median is shown as dashed black line, Q1 and Q3 are dotted cyan lines, IQR – interquartile range, Max – maximal loop length, Bin – size of bins used to plot the data, OL – outliers, the number indicate the size of the loops which fall above 99^th^ percentile of the distribution. Human T2T-CHM13v2.0 genome was used for calculation. LoopG and LoopC denote G-rich and C-rich orientation of the pattern which was used for matching.

**Figure S5.**
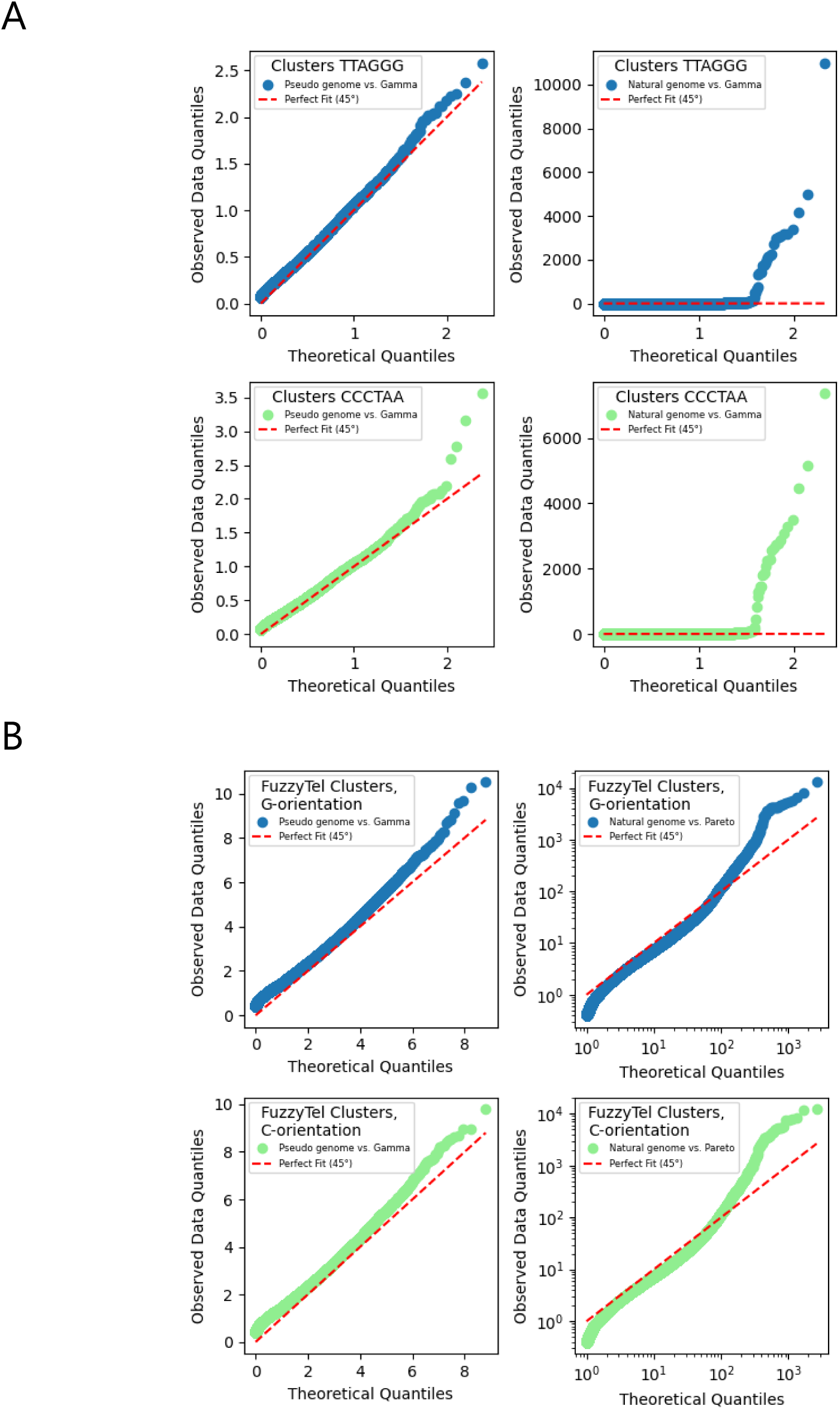
QQ-plots of the CS (Cluster Score) quantiles for the natural and pseudo genomes vs. theoretical quantiles of Gamma and Pareto distributions. (A) Clusters of canonical telomeric repeats and (B) Clusters formed by FuzzyTel pattern.

**Figure S6.**
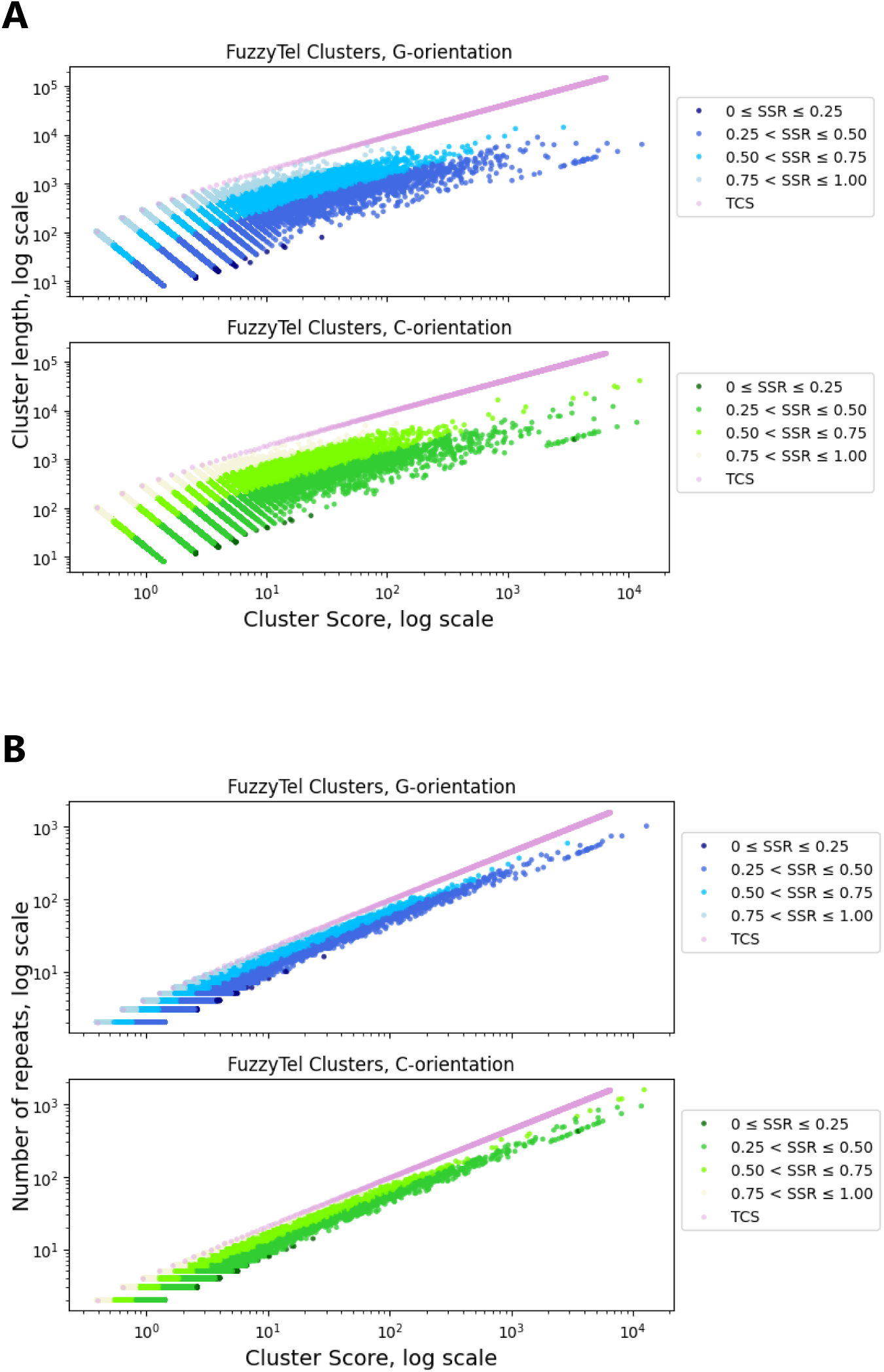
Relationship between (A) CS (Cluster Score) and Cluster length and (B) CS (Cluster Score) and number of repeats for the clusters formed by FuzzyTel pattern. TCS represents a simulation of a pattern generated by random choice of telomeric-like repeats spaced equally by the 90 nt intervals. Double decimal logarithm scale.

**Figure S7.**
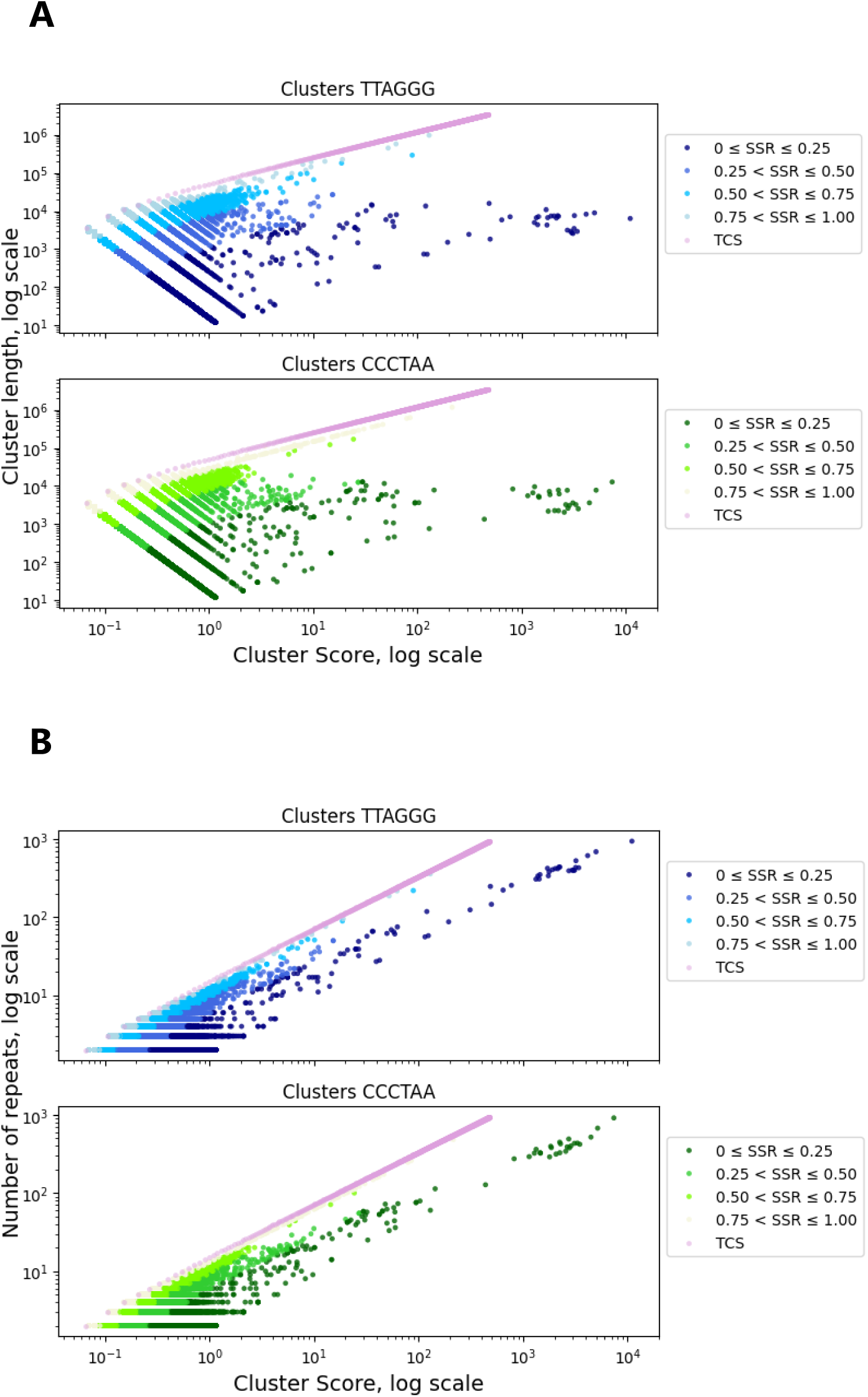
Relationship between (A) CS (Cluster Score) and Cluster length and (B) CS (Cluster Score) and number of repeats for canonical telomeric clusters. TCS represents a simulation of a pattern generated by telomeric repeats spaced equally by the 3660 nt intervals. Double decimal logarithm scale.

**Figure S8.**
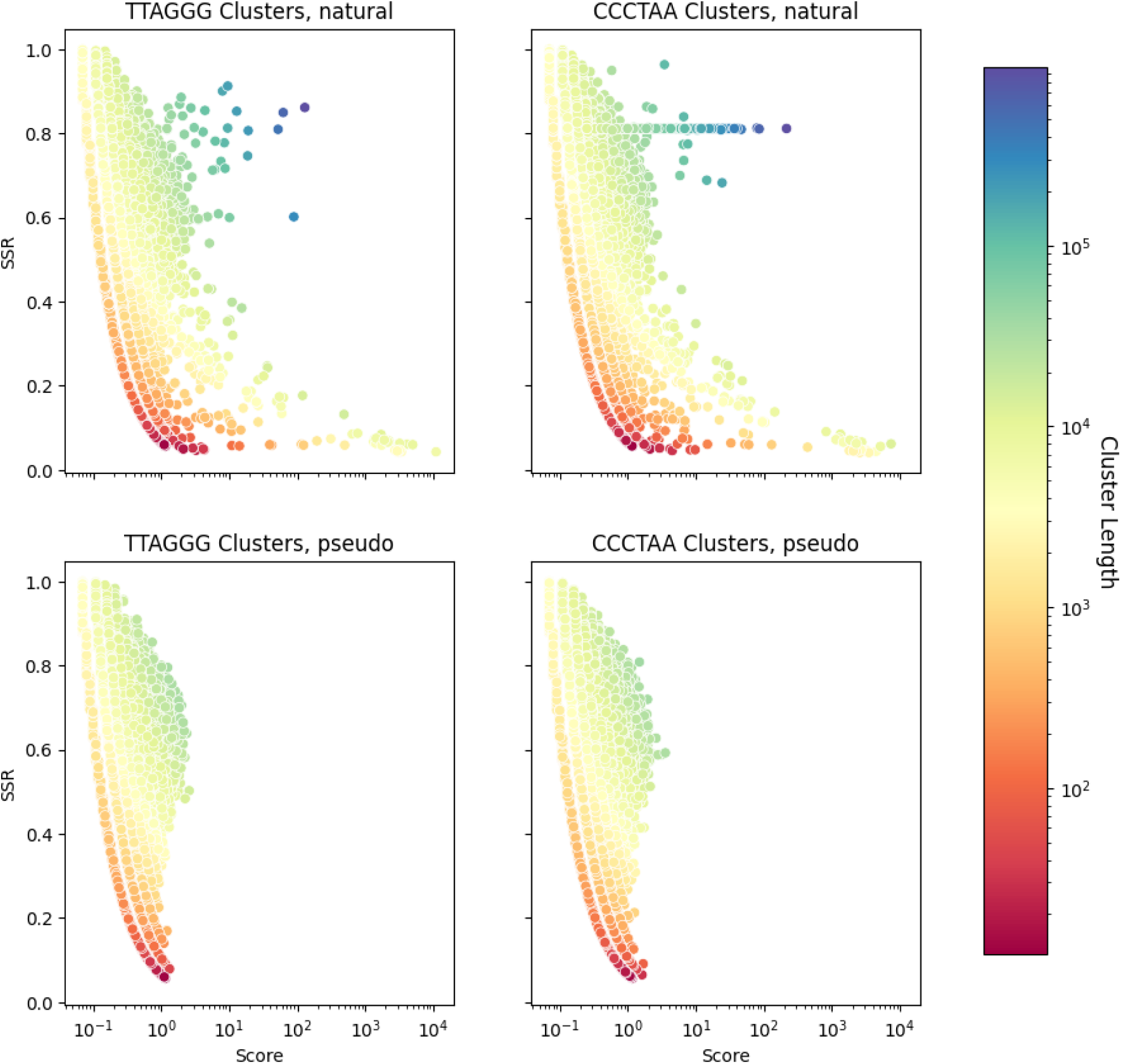
CS (Score) vs. SSR distribution with color reflecting cluster length. Clusters generated by matching the canonical telomeric pattern in the T2T-CHM13v2.0 natural human genome and pseudo genome are shown for the G-rich and C-rich orientations. X-axis is decimal logarithmic.

**Figure S9.**
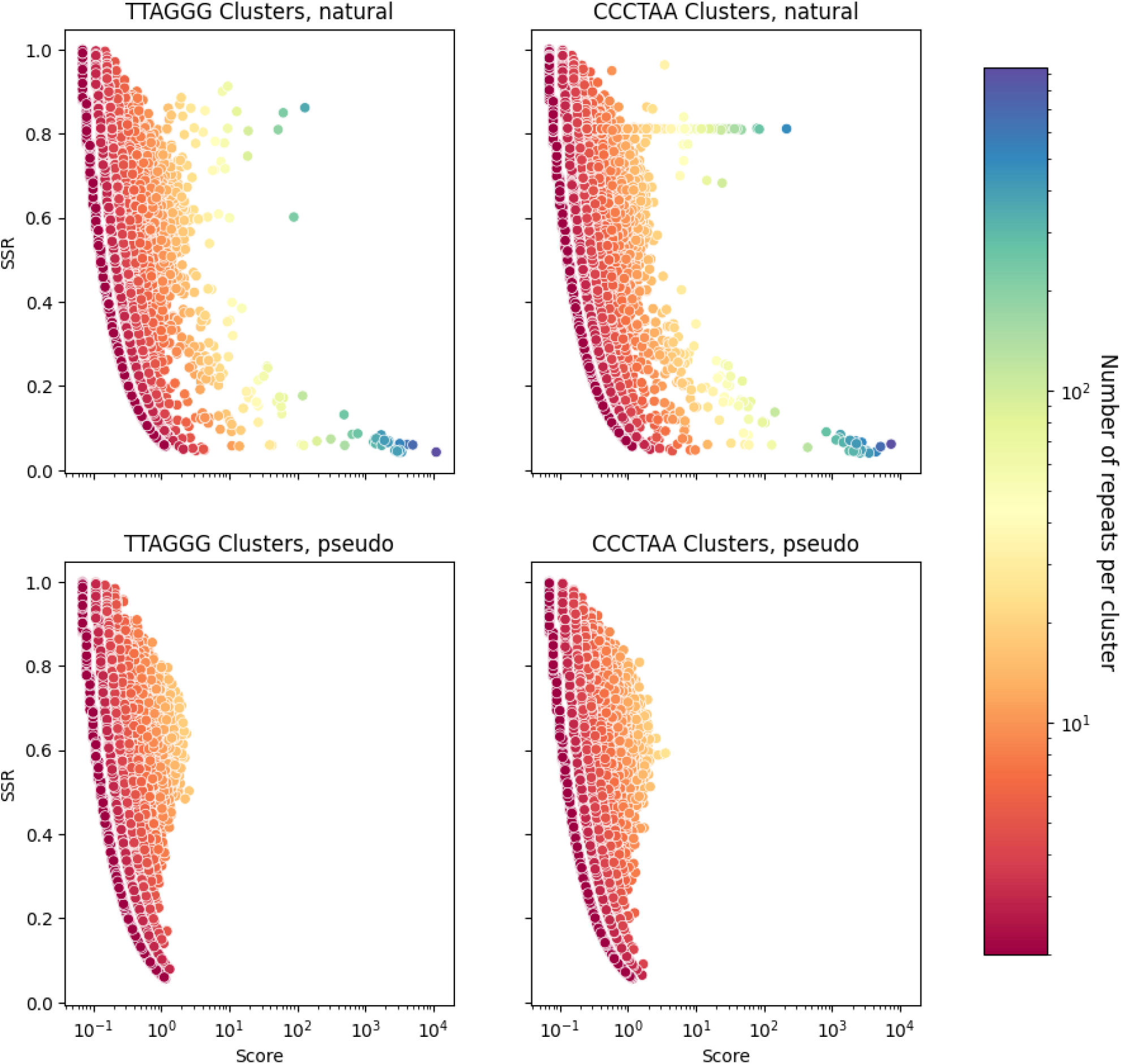
CS (Score) vs. SSR distribution with color reflecting number of repeats per cluster. Clusters generated by matching the canonical telomeric pattern in the T2T-CHM13v2.0 genome and pseudo genome are shown for the two G-rich and C-rich orientations. X-axis is decimal logarithmic.

**Figure S10.**
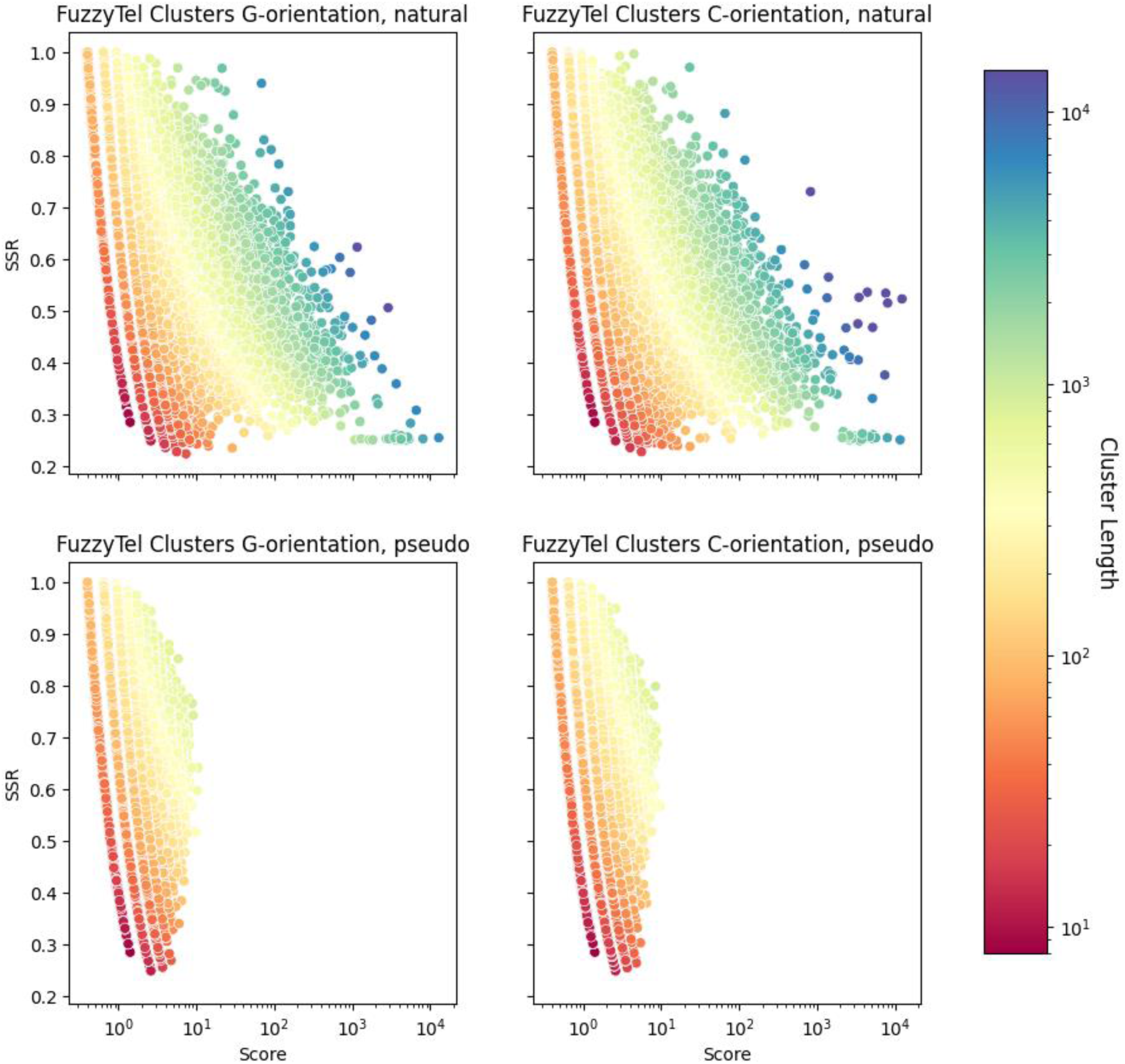
CS (Score) vs. SSR distribution with color reflecting cluster length. Clusters generated by matching FuzzyTel pattern in the T2T-CHM13v2.0 genome and pseudo genome are shown for the two G-rich and C-rich orientations. X-axis is decimal logarithmic.

**Figure S11.**
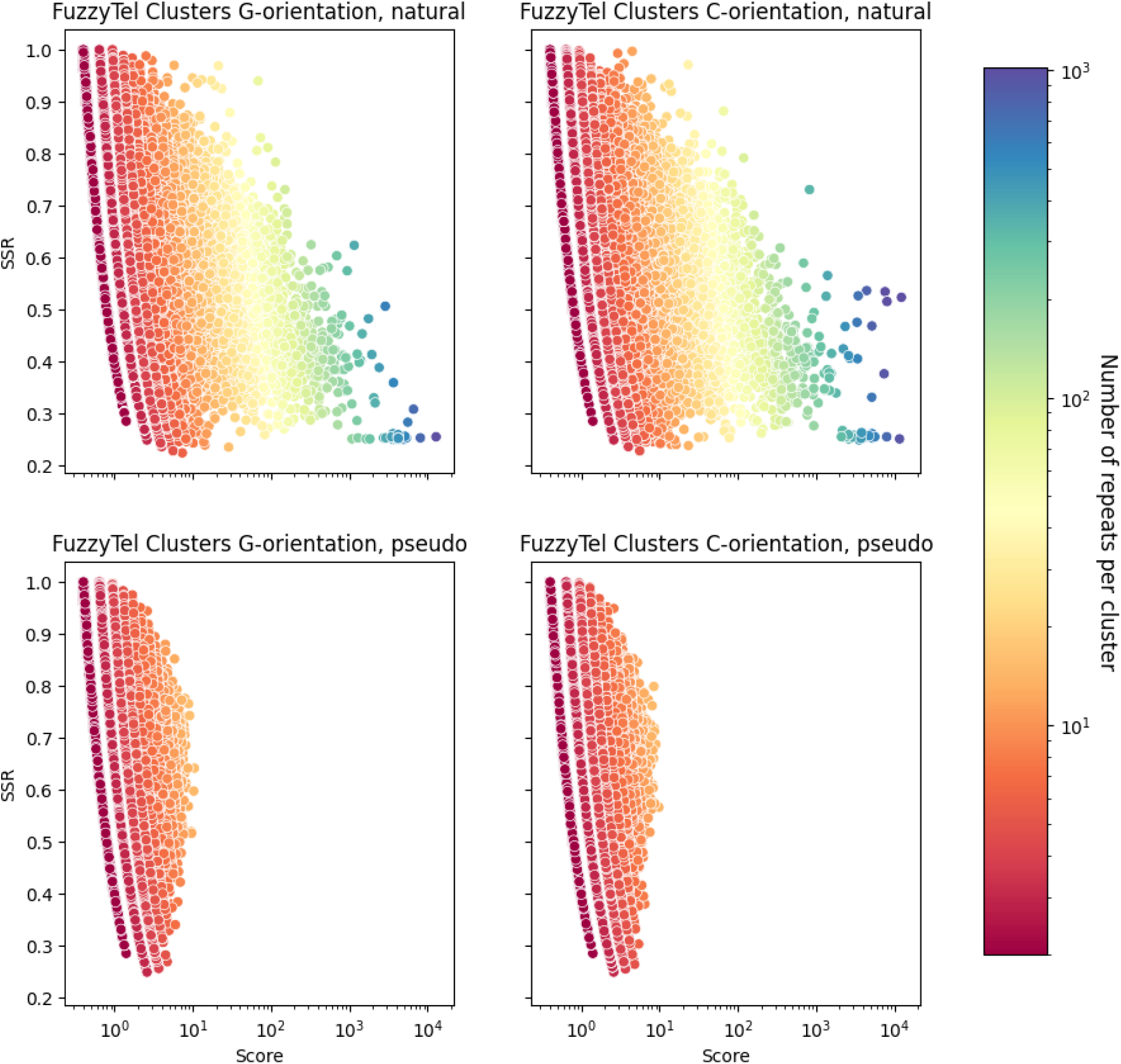
CS (Score) vs. SSR distribution with color reflecting number of repeats per cluster. Clusters generated by matching FuzzyTel pattern in the T2T-CHM13v2.0 genome and pseudo genome are shown for the two G-rich and C-rich orientations. X-axis is decimal logarithmic.

**Figure S12.**
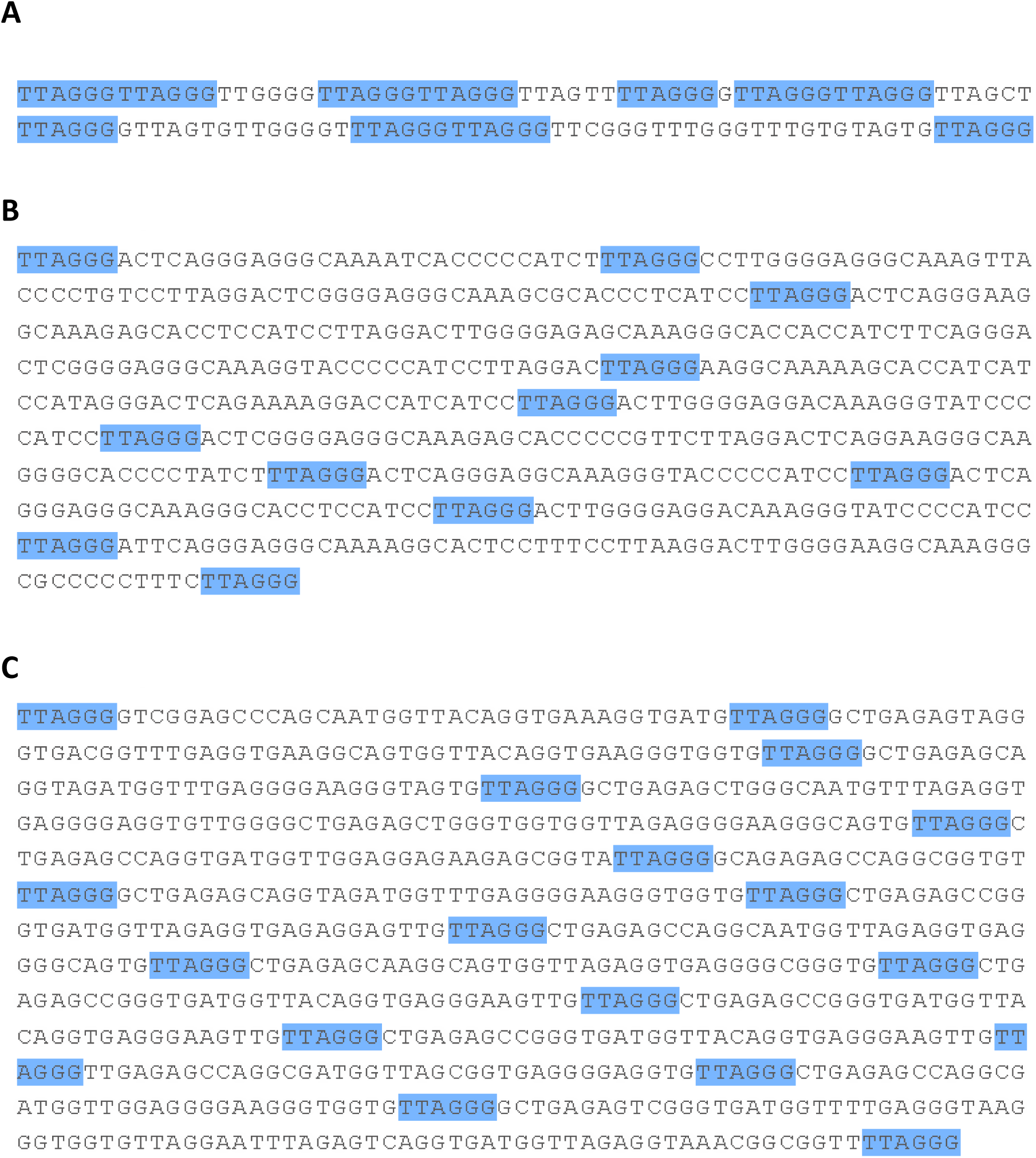
Examples of clusters enriched with TTAGGG motif in T2T-CHM13v2.0 genome with CS ≥1.2 and SSR ≤ 0.5. A. chr9 150394480-150394602; B. chr10 129922360-129922926; C. chr11 120816613-120817402.

**Figure S13.**
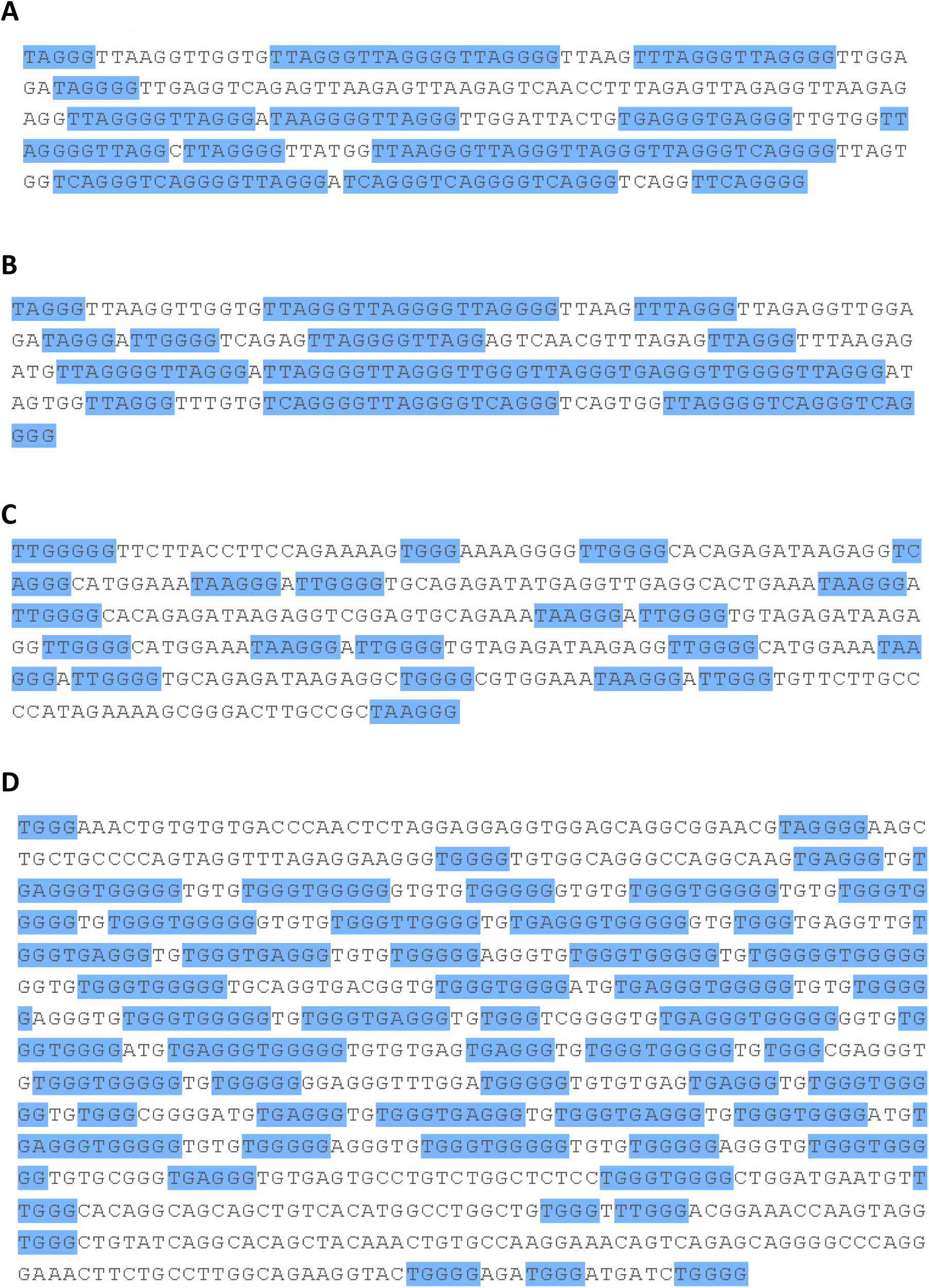
Examples of clusters enriched by FuzzyTel G-rich orientation pattern in T2T-CHM13v2.0 genome with CS ≥10 and SSR ≤ 0.5. A. chr1 127776462-127776760; B. chr9 40801838-40802085; C. chr8 12053913-12054248; chr5 176574224-176575127.

**Figure S14.**
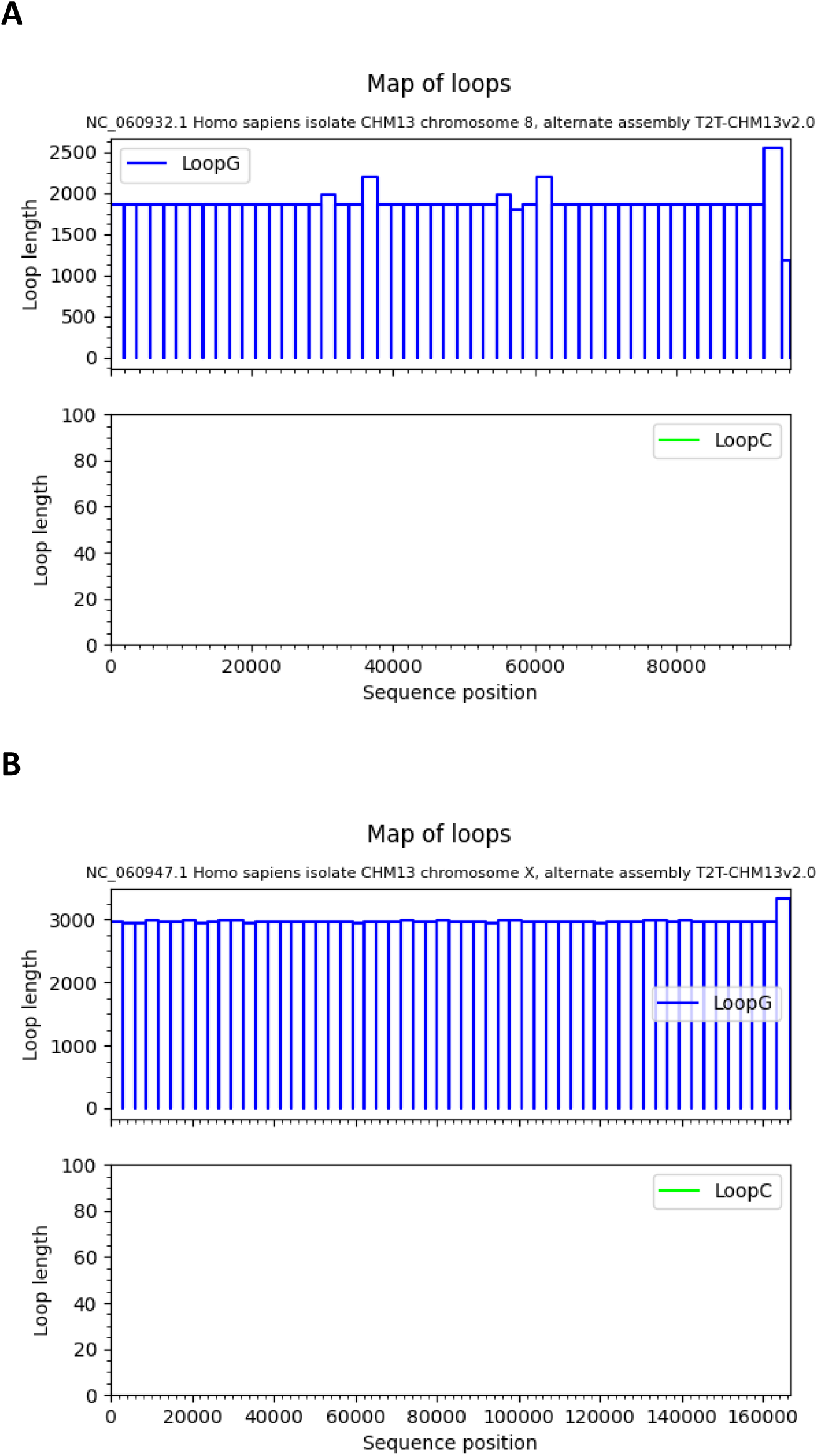
Examples of sparse long clusters characterized by near regular distribution of TTAGGG motif in T2T-CHM13v2.0 genome enriched with parameters CS ≥1.2 and SSR > 0.5. A. chr8 44243811-44339964; B chr X 114134178-114300842. Sequence position on the plots is given from the starting point of clusters.

**Figure S15.**
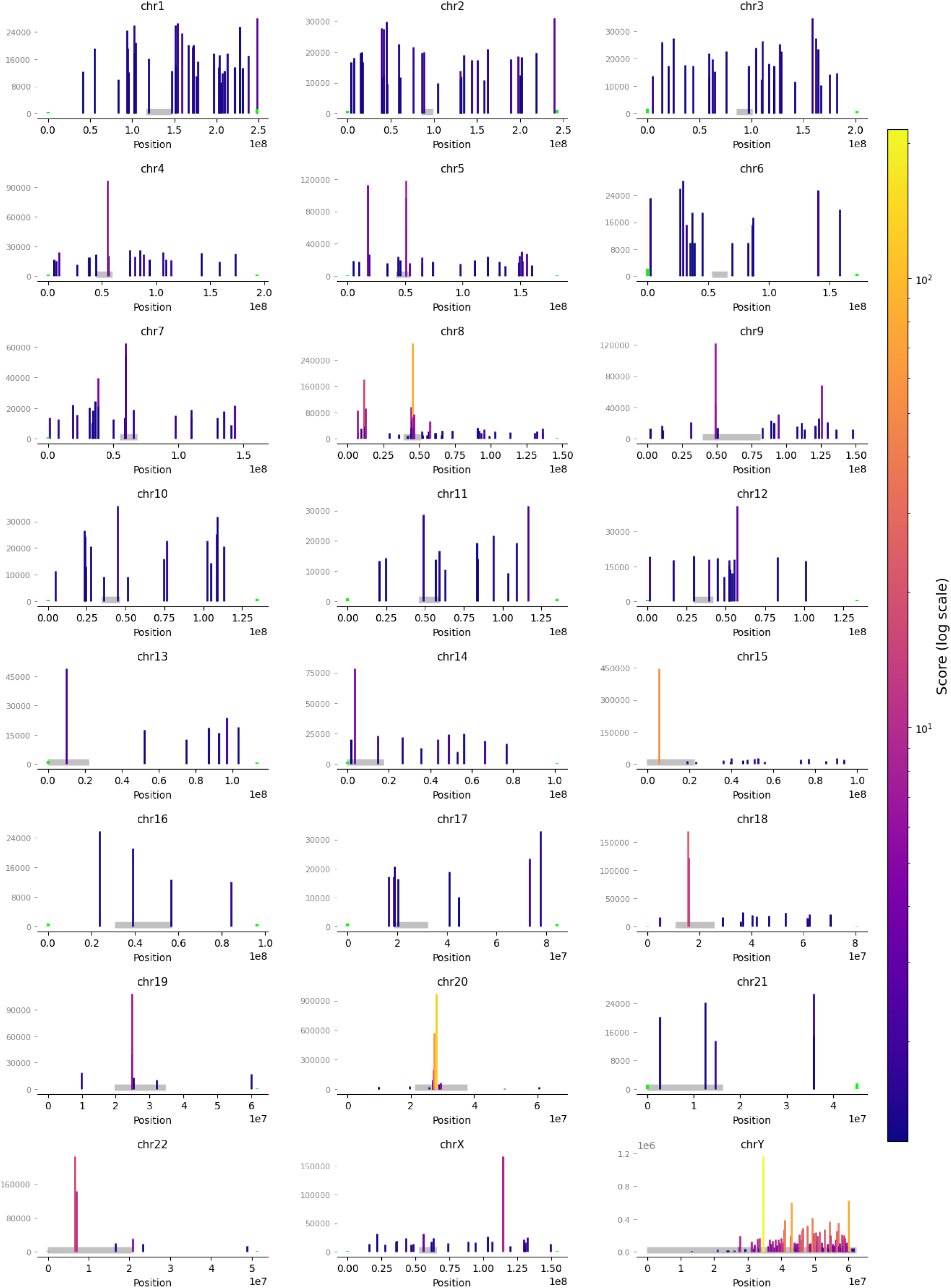
Distribution of sparse clusters CS ≥1.2 and SSR > 0.5 containing canonical telomeric repeats in different chromosomes

**Figure S16.**
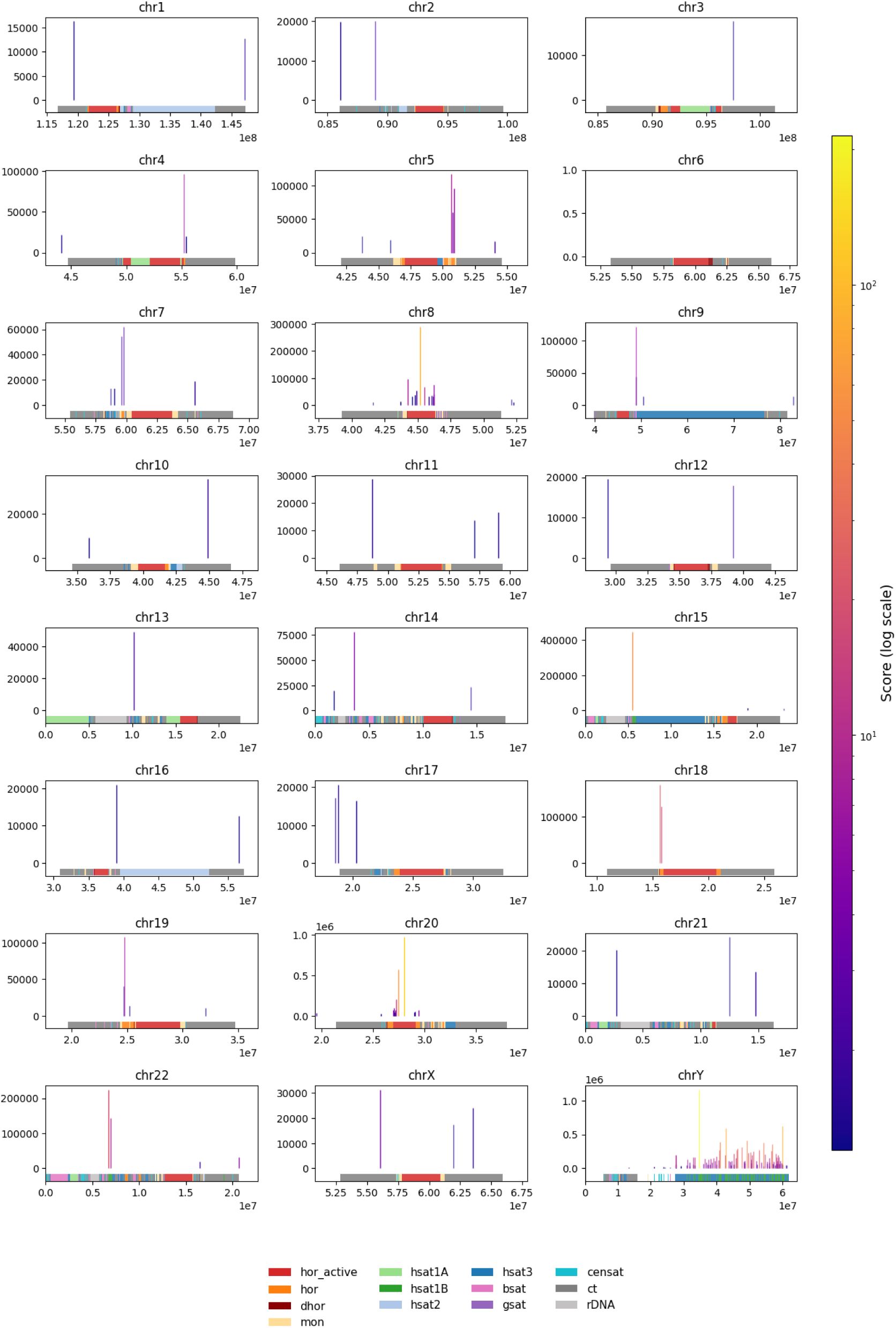
Sparse clusters containing canonical telomeric repeats (CS ≥1.2 and SSR > 0.5) in centromeric regions

**Figure S17.**
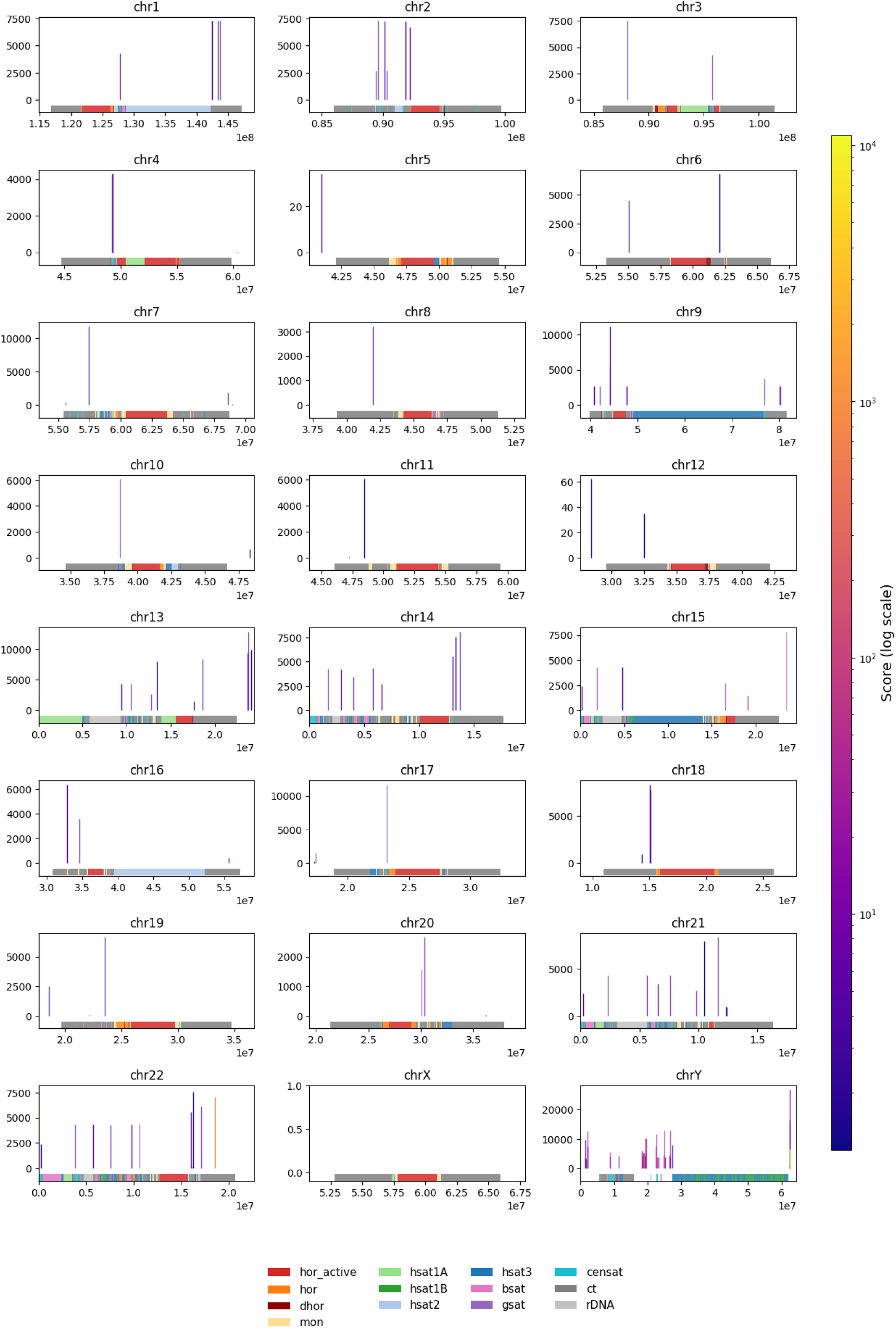
Dense clusters containing canonical telomeric repeats (CS ≥1.2 and SSR ≤ 0.5) in centromeric regions

**Figure S18.**
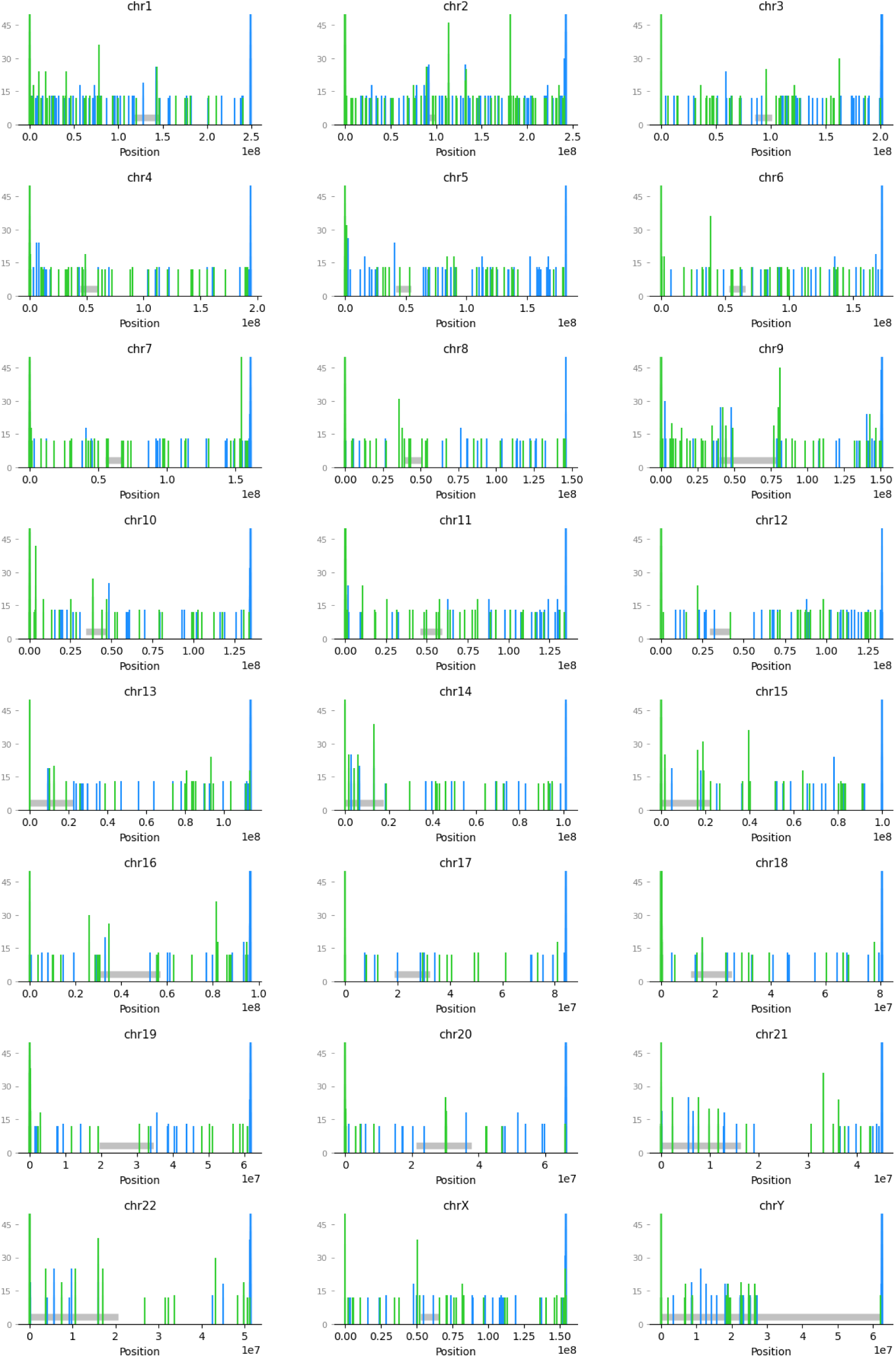
Distribution of tandem canonical telomeric clusters with R ≥ 2 and loop length ≤ 1 along chromosomes of the T2T-CHM13v2.0 genome. TTAGGG are blue, CCCTAA are green, centromere positions are gray shadows. Telomeres are represented on both sides and are truncated - y axes are set up to 50 nt to enlarge ITSs.

