## Supplementary file1 for "FuzzyClusTeR: a web server for analysis of tandem and diffuse DNA repeat clusters with application to telomeric-like repeats"

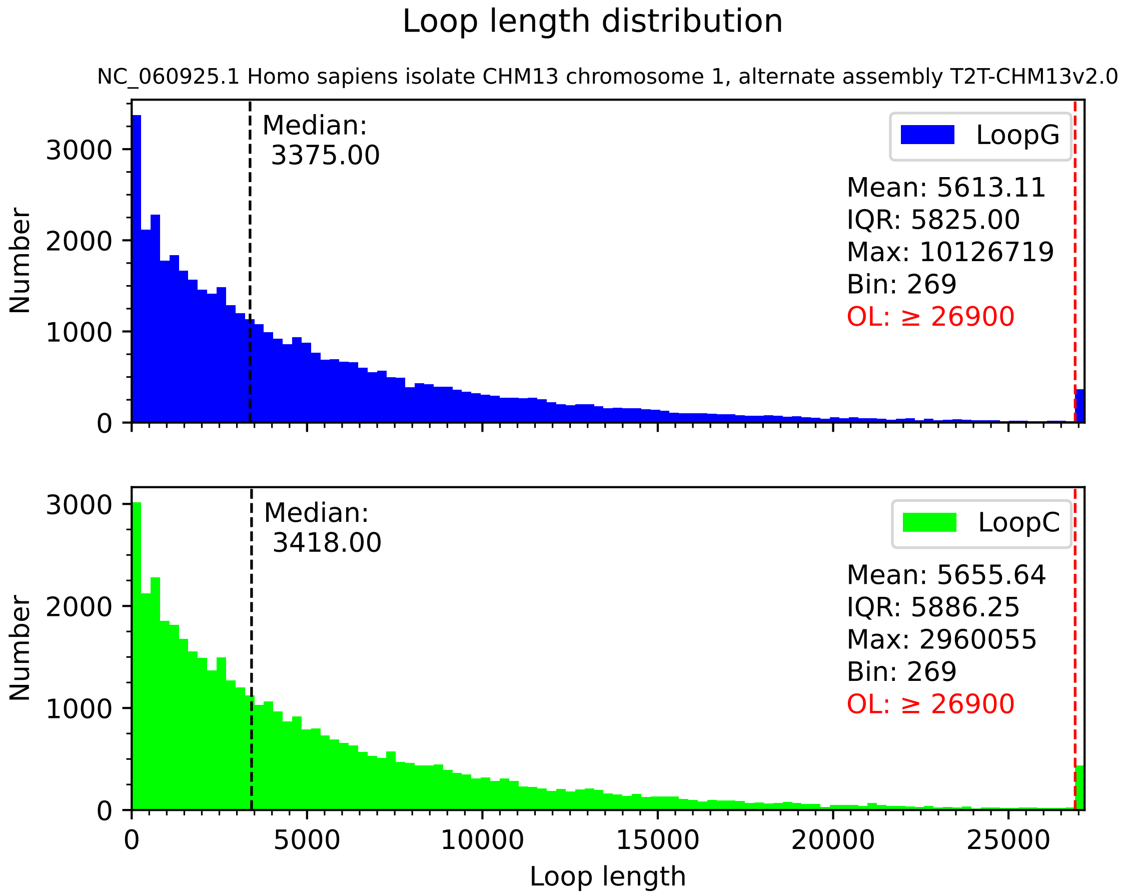
Loop length distribution
NC_060925.1 Homo sapiens isolate CHM13 chromosome 1, alternate assembly T2T-CHM13v2.0
Median:
3375.00
LoopG
Mean: 5613.11
IQR: 5825.00
Max: 10126719
Bin: 269
OL: ≥ 26900
Number
0
1000
2000
3000
Median:
3418.00
LoopC
Mean: 5655.64
IQR: 5886.25
Max: 2960055
Bin: 269
OL: ≥ 26900
Number
0
1000
2000
3000
0
5000
10000
15000
20000
25000
Loop length

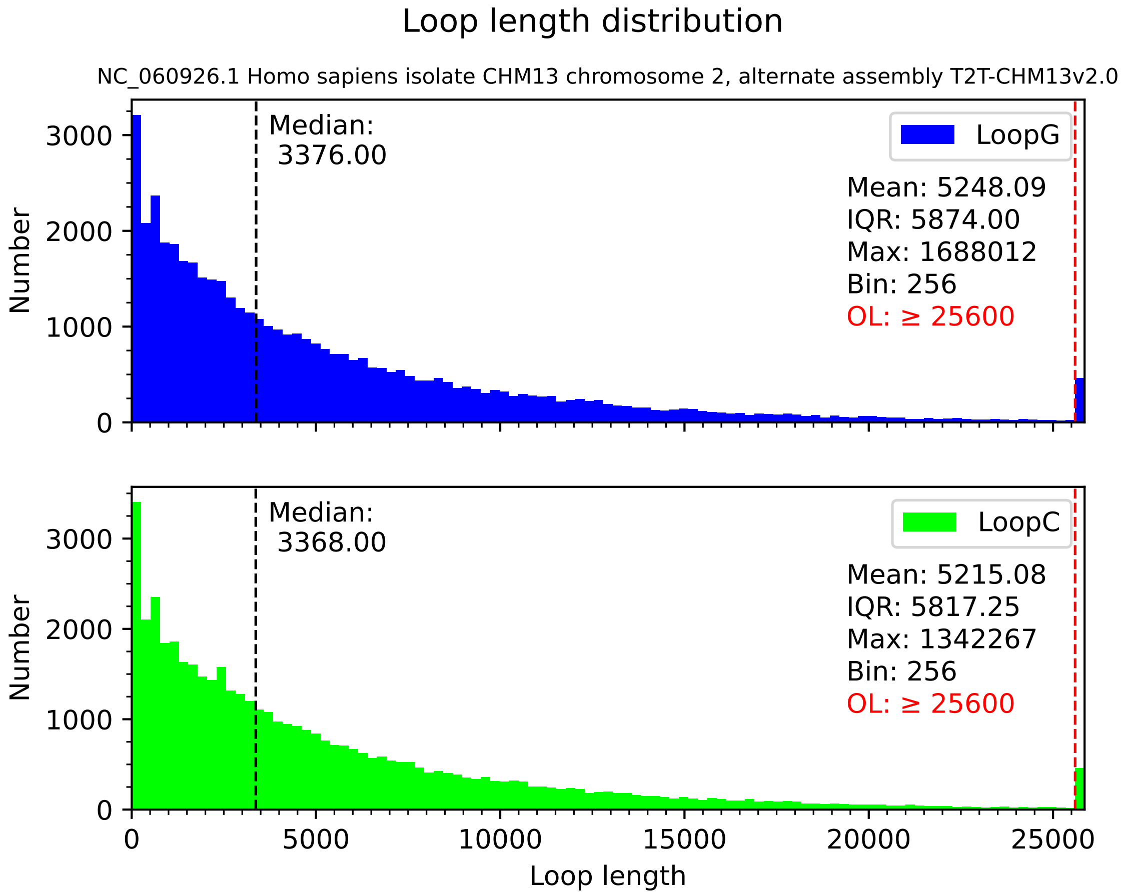
Loop length distribution
NC_060926.1 Homo sapiens isolate CHM13 chromosome 2, alternate assembly T2T-CHM13v2.0
Median:
3376.00
LoopG
Mean: 5248.09
IQR: 5874.00
Max: 1688012
Bin: 256
OL: ≥ 25600
Number
0
1000
2000
3000
Median:
3368.00
LoopC
Mean: 5215.08
IQR: 5817.25
Max: 1342267
Bin: 256
OL: ≥ 25600
0
5000
10000
15000
20000
25000
Loop length

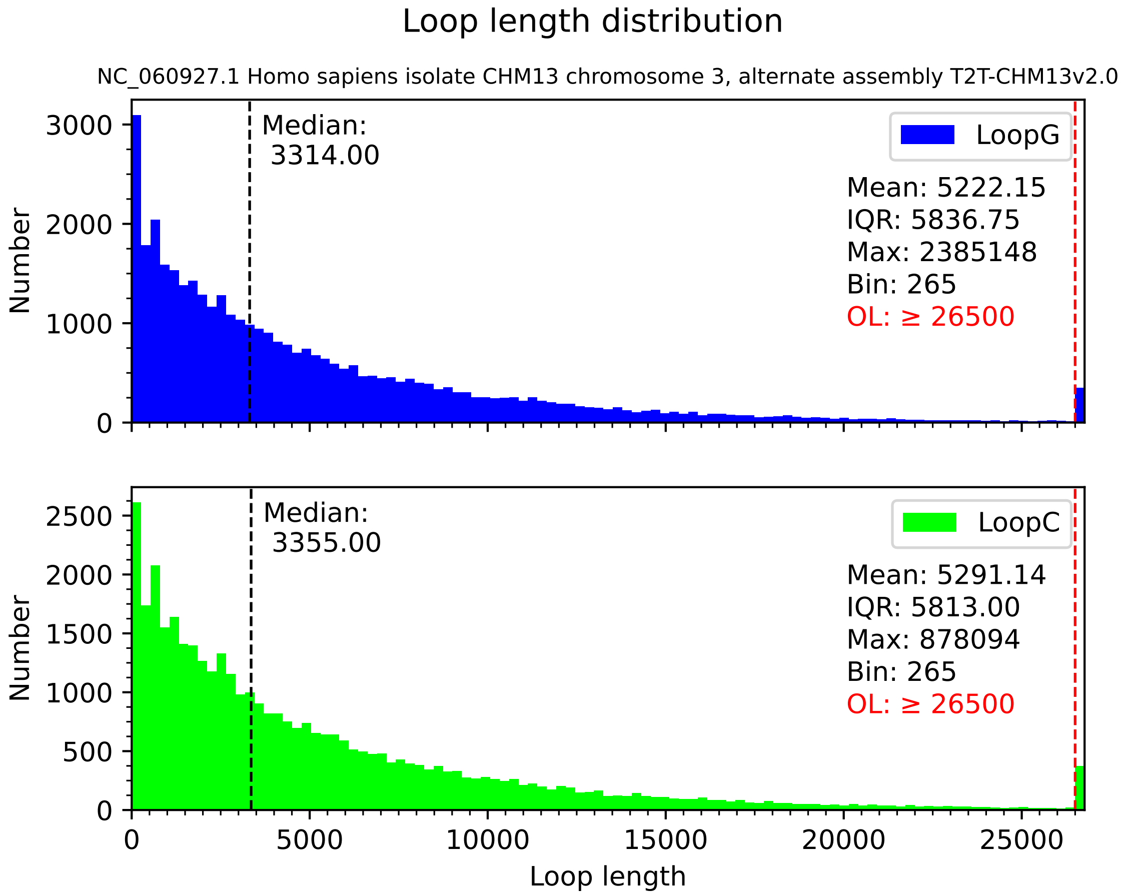
Loop length distribution
NC_060927.1 Homo sapiens isolate CHM13 chromosome 3, alternate assembly T2T-CHM13v2.0
Median:
3314.00
LoopG
Mean: 5222.15
IQR: 5836.75
Max: 2385148
Bin: 265
OL: ≥ 26500
Number
0
1000
2000
3000
Median:
3355.00
LoopC
Mean: 5291.14
IQR: 5813.00
Max: 878094
Bin: 265
OL: ≥ 26500
Number
0
500
1000
1500
2000
2500
0
5000
10000
15000
20000
25000
Loop length

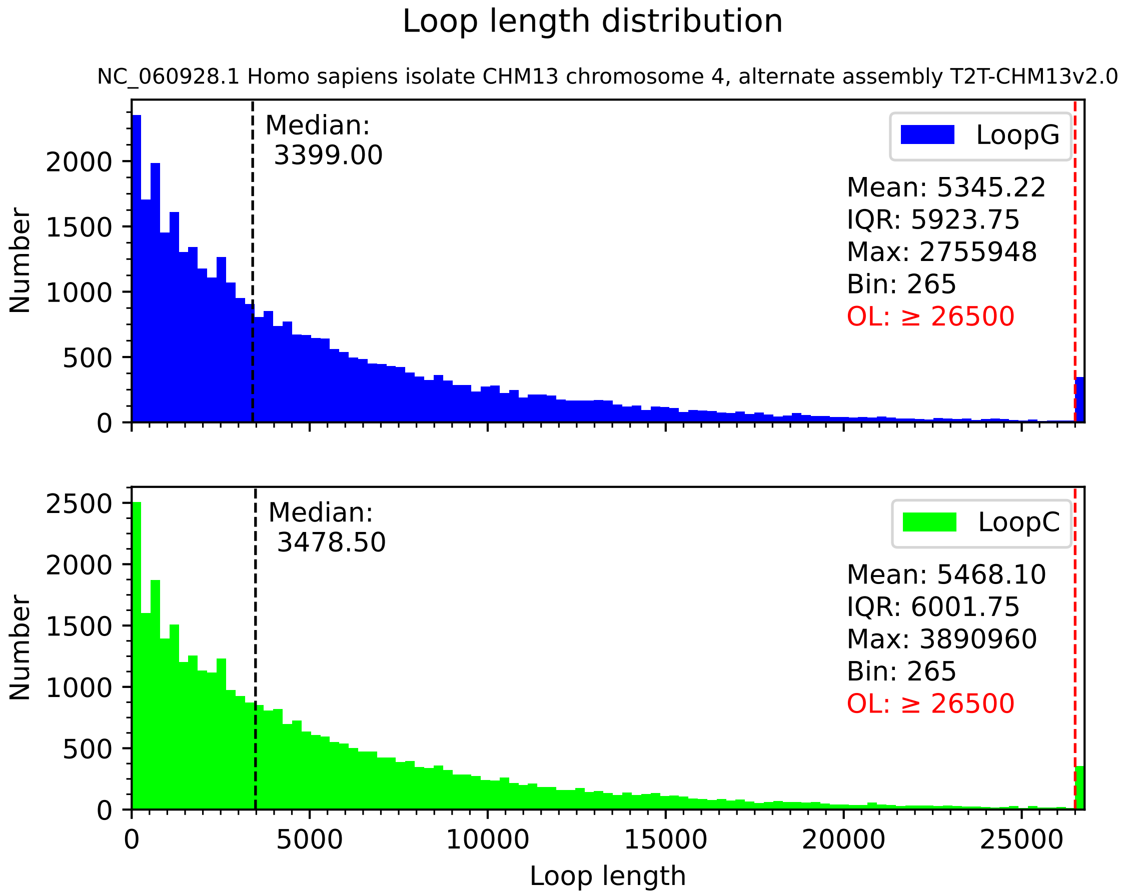
Loop length distribution
NC_060928.1 Homo sapiens isolate CHM13 chromosome 4, alternate assembly T2T-CHM13v2.0
Median:
3399.00
LoopG
Mean: 5345.22
IQR: 5923.75
Max: 2755948
Bin: 265
OL: ≥ 26500
Number
0
500
1000
1500
2000
Median:
3478.50
LoopC
Mean: 5468.10
IQR: 6001.75
Max: 3890960
Bin: 265
OL: ≥ 26500
Number
0
500
1000
1500
2000
2500
0
5000
10000
15000
20000
25000
Loop length

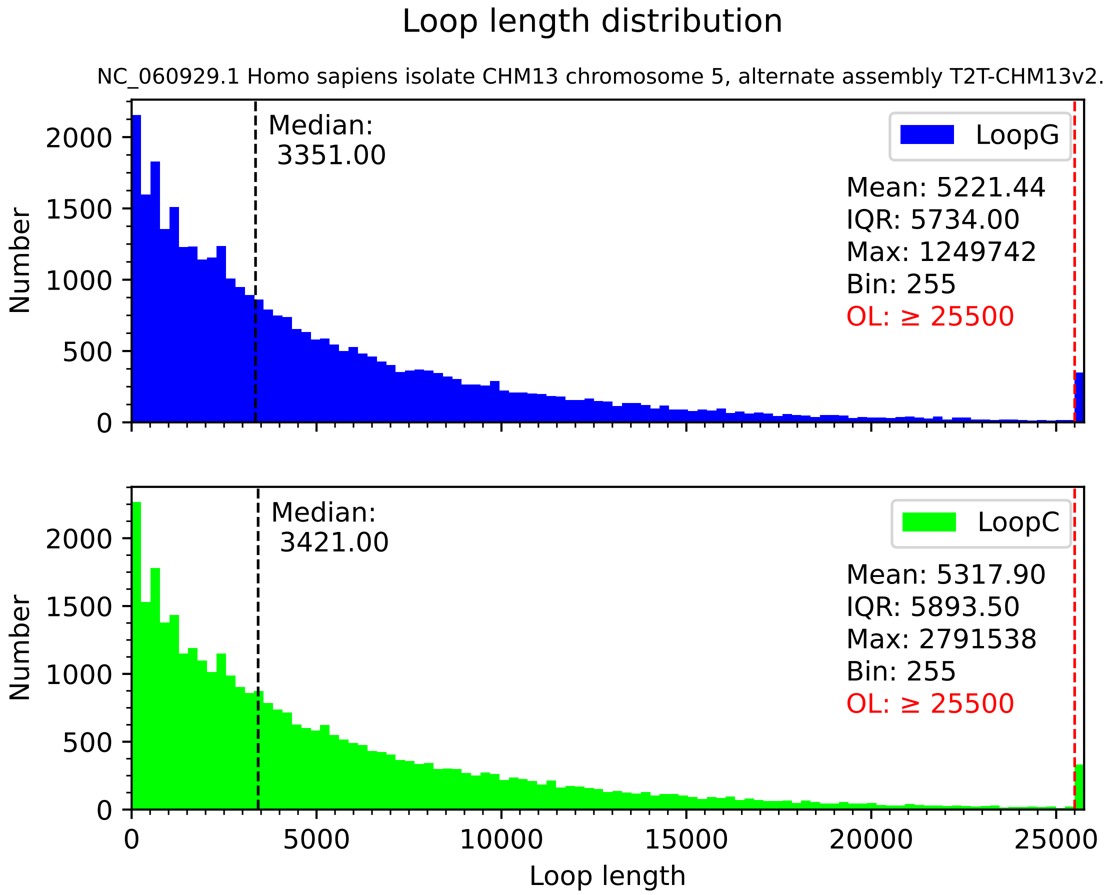

Loop length distribution
NC_060929.1 Homo sapiens isolate CHM13 chromosome 5, alternate assembly T2T-CHM13v2.0
Median:
3351.00
LoopG
Mean: 5221.44
IQR: 5734.00
Max: 1249742
Bin: 255
OL: ≥ 25500
Median:
3421.00
LoopC
Mean: 5317.90
IQR: 5893.50
Max: 2791538
Bin: 255
OL: ≥ 25500
Number
Loop length
0
500
1000
1500
2000
5000
10000
15000
20000
25000

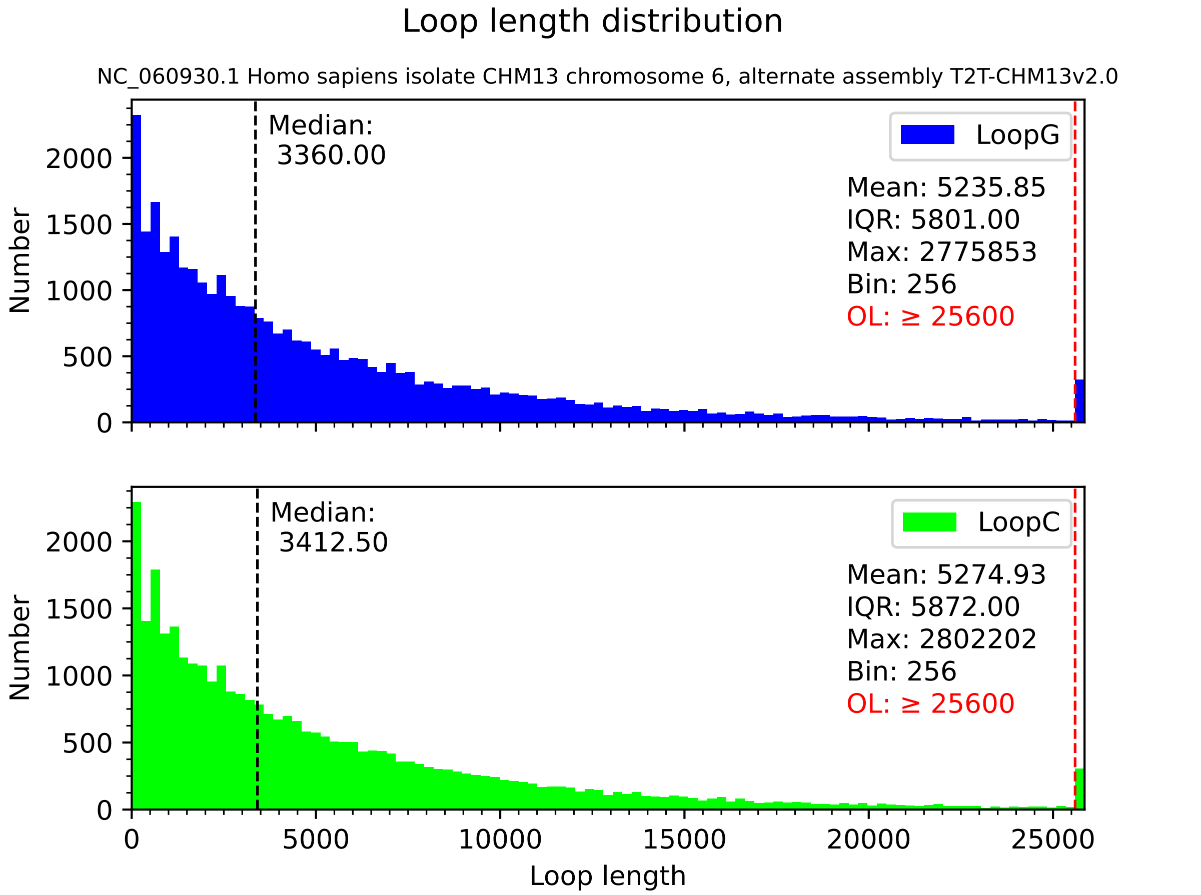
Loop length distribution
NC_060930.1 Homo sapiens isolate CHM13 chromosome 6, alternate assembly T2T-CHM13v2.0
Median:
3360.00
LoopG
Mean: 5235.85
IQR: 5801.00
Max: 2775853
Bin: 256
OL: ≥ 25600
Median:
3412.50
LoopC
Mean: 5274.93
IQR: 5872.00
Max: 2802202
Bin: 256
OL: ≥ 25600
Number
Number
0
500
1000
1500
2000
0
5000
10000
15000
20000
25000
Loop length

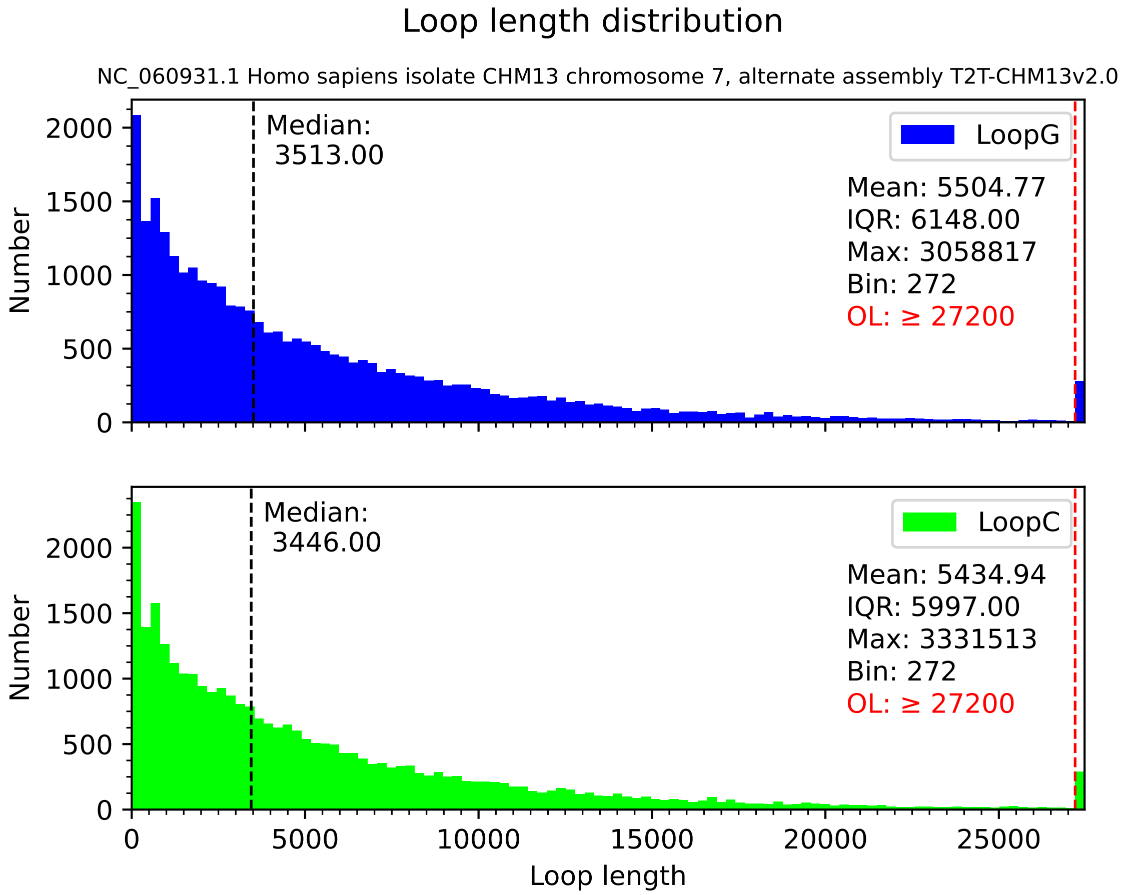
Loop length distribution
NC_060931.1 Homo sapiens isolate CHM13 chromosome 7, alternate assembly T2T-CHM13v2.0
Median:
3513.00
LoopG
Mean: 5504.77
IQR: 6148.00
Max: 3058817
Bin: 272
OL: ≥ 27200
Median:
3446.00
LoopC
Mean: 5434.94
IQR: 5997.00
Max: 3331513
Bin: 272
OL: ≥ 27200
Number
Loop length
0
500
1000
1500
2000
5000
10000
15000
20000
25000

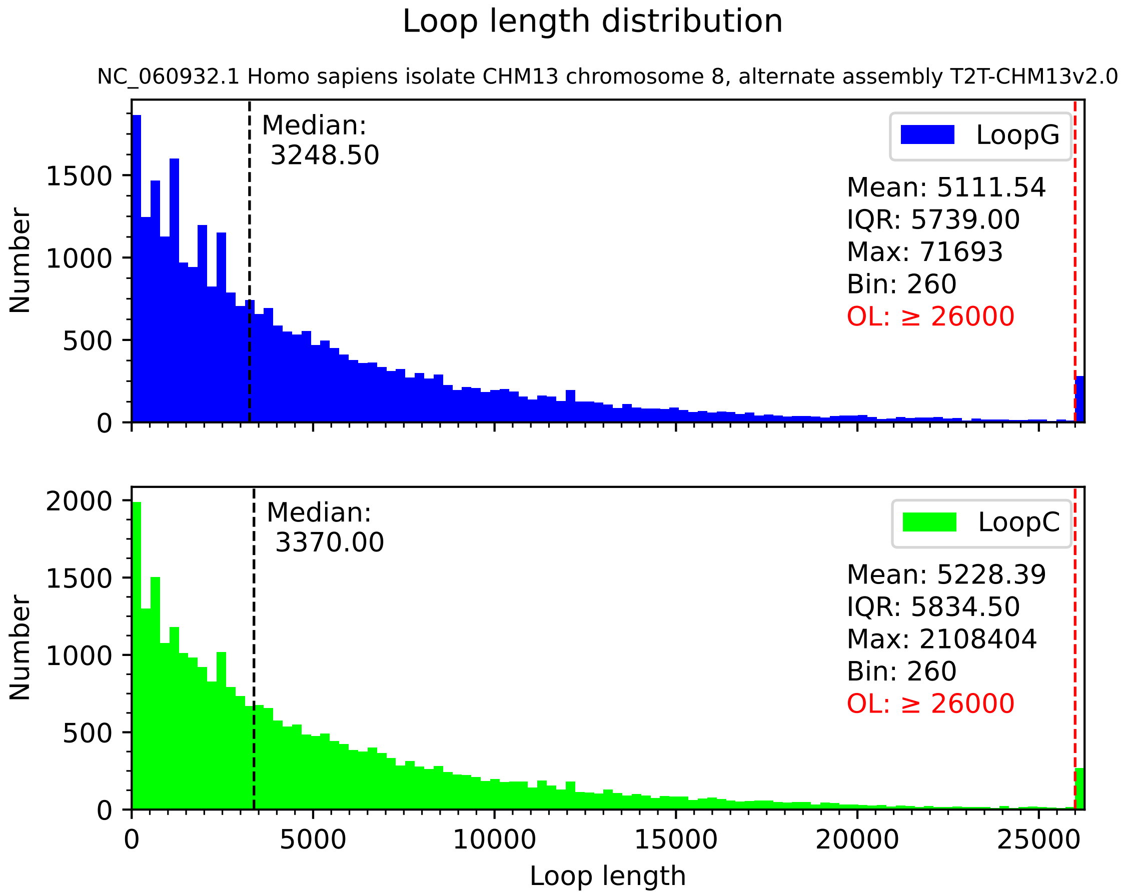
Loop length distribution
NC_060932.1 Homo sapiens isolate CHM13 chromosome 8, alternate assembly T2T-CHM13v2.0
Median:
3248.50
LoopG
Mean: 5111.54
IQR: 5739.00
Max: 71693
Bin: 260
OL: ≥ 26000
Median:
3370.00
LoopC
Mean: 5228.39
IQR: 5834.50
Max: 2108404
Bin: 260
OL: ≥ 26000
Number
Loop length
0
500
1000
1500
2000
5000
10000
15000
20000
25000

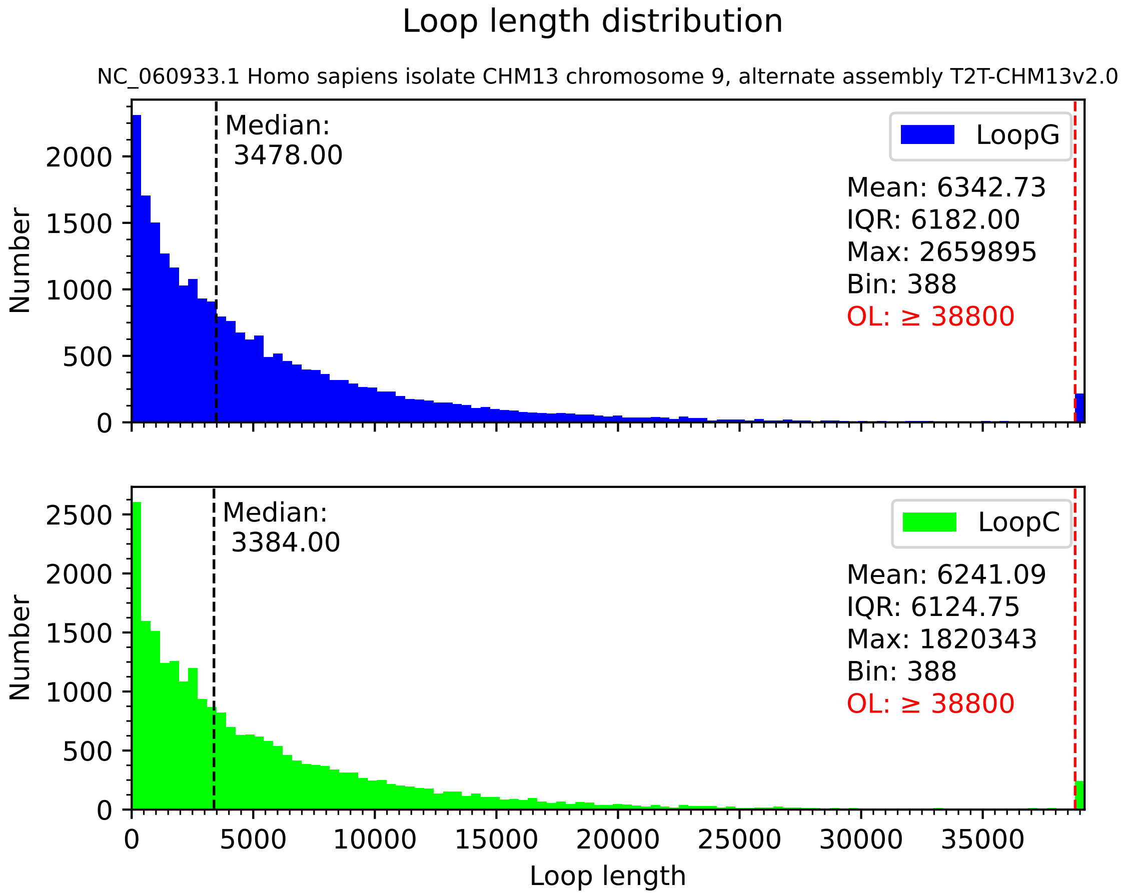
Loop length distribution
NC_060933.1 Homo sapiens isolate CHM13 chromosome 9, alternate assembly T2T-CHM13v2.0
Median:
3478.00
LoopG
Mean: 6342.73
IQR: 6182.00
Max: 2659895
Bin: 388
OL: ≥ 38800
Number
0
500
1000
1500
2000
Median:
3384.00
LoopC
Mean: 6241.09
IQR: 6124.75
Max: 1820343
Bin: 388
OL: ≥ 38800
Number
0
500
1000
1500
2000
2500
0
5000
10000
15000
20000
25000
30000
35000
Loop length

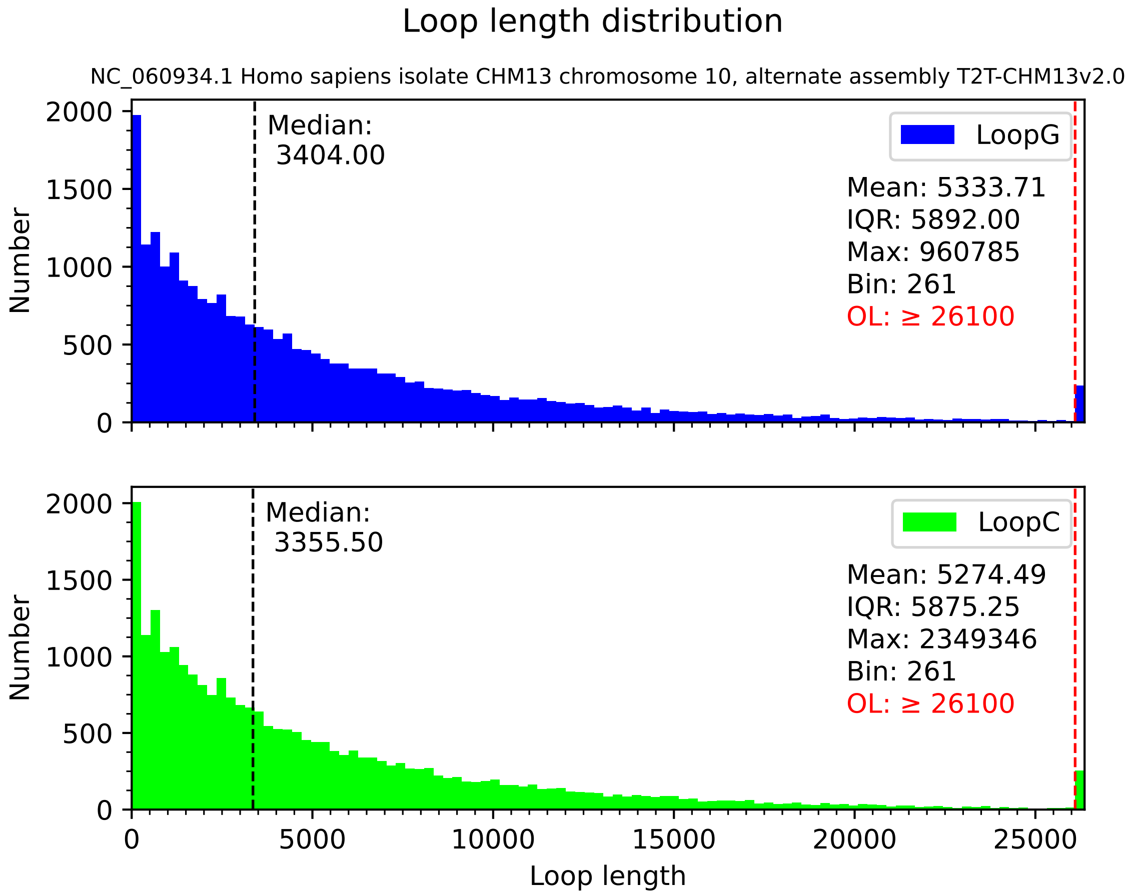

Loop length distribution
NC_060934.1 Homo sapiens isolate CHM13 chromosome 10, alternate assembly T2T-CHM13v2.0
Median:
3404.00
LoopG
Mean: 5333.71
IQR: 5892.00
Max: 960785
Bin: 261
OL: ≥ 26100
Number
0
500
1000
1500
2000
Median:
3355.50
LoopC
Mean: 5274.49
IQR: 5875.25
Max: 2349346
Bin: 261
OL: ≥ 26100
Number
0
500
1000
1500
2000
0
5000
10000
15000
20000
25000
Loop length

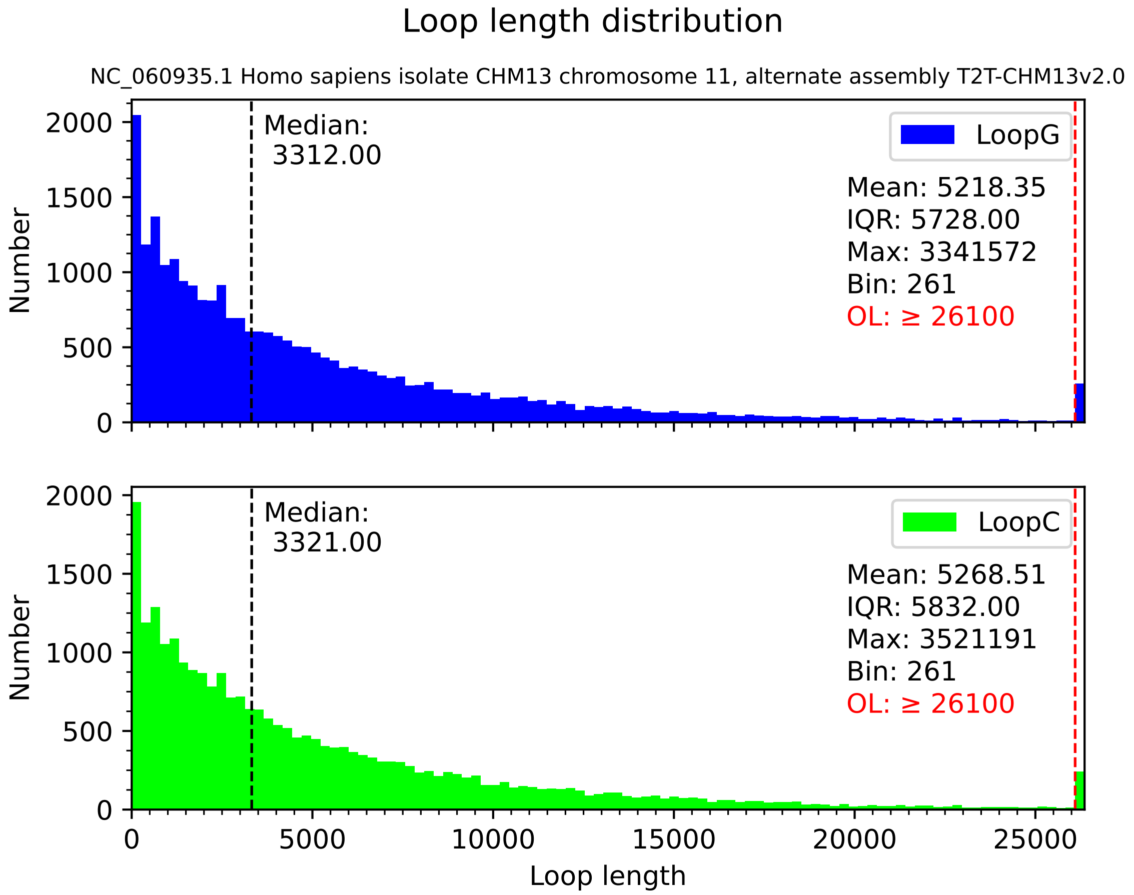
Loop length distribution
NC_060935.1 Homo sapiens isolate CHM13 chromosome 11, alternate assembly T2T-CHM13v2.0
Median:
3312.00
LoopG
Mean: 5218.35
IQR: 5728.00
Max: 3341572
Bin: 261
OL: ≥ 26100
Median:
3321.00
LoopC
Mean: 5268.51
IQR: 5832.00
Max: 3521191
Bin: 261
OL: ≥ 26100
Number
Number
Loop length
0
500
1000
1500
2000
0
5000
10000
15000
20000
25000

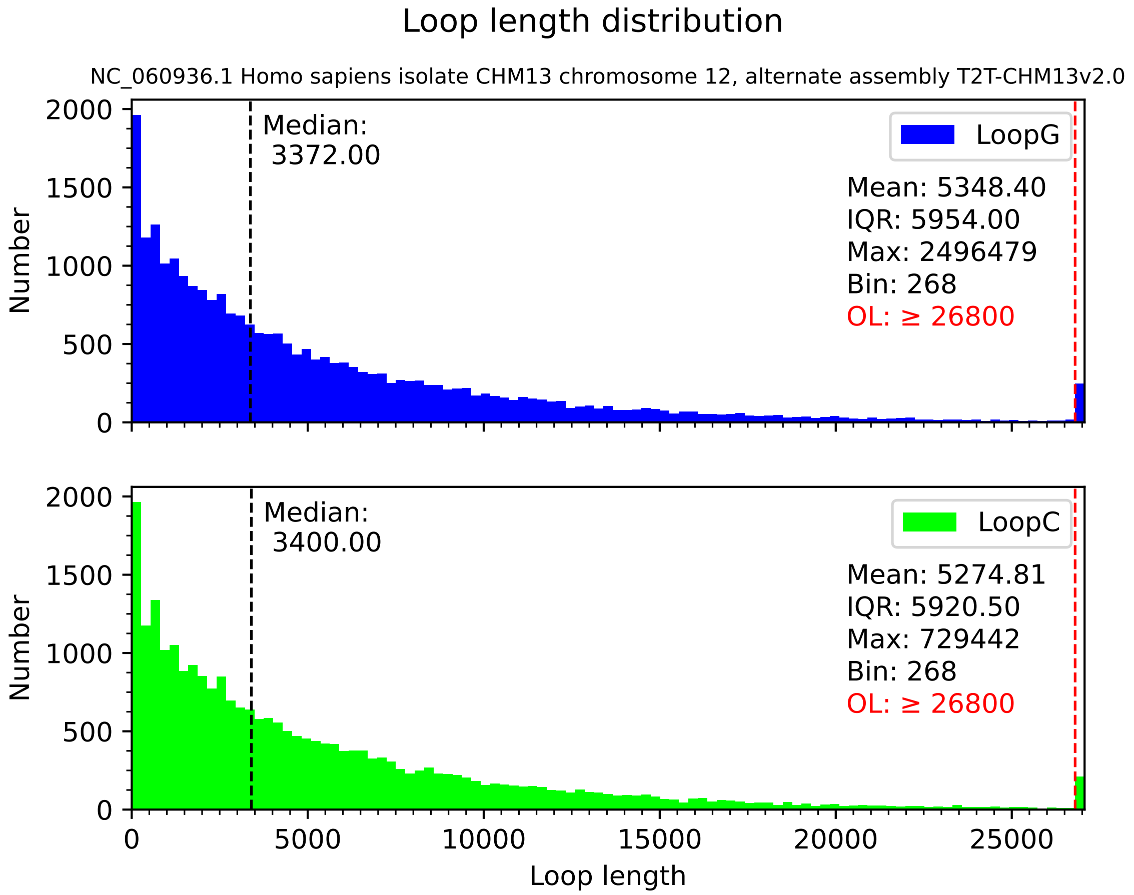

Loop length distribution
NC_060936.1 Homo sapiens isolate CHM13 chromosome 12, alternate assembly T2T-CHM13v2.0
Number
Median:
3372.00
LoopG
Mean: 5348.40
IQR: 5954.00
Max: 2496479
Bin: 268
OL: ≥ 26800
Median:
3400.00
LoopC
Mean: 5274.81
IQR: 5920.50
Max: 729442
Bin: 268
OL: ≥ 26800
0
500
1000
1500
2000
0
5000
10000
15000
20000
25000
Loop length

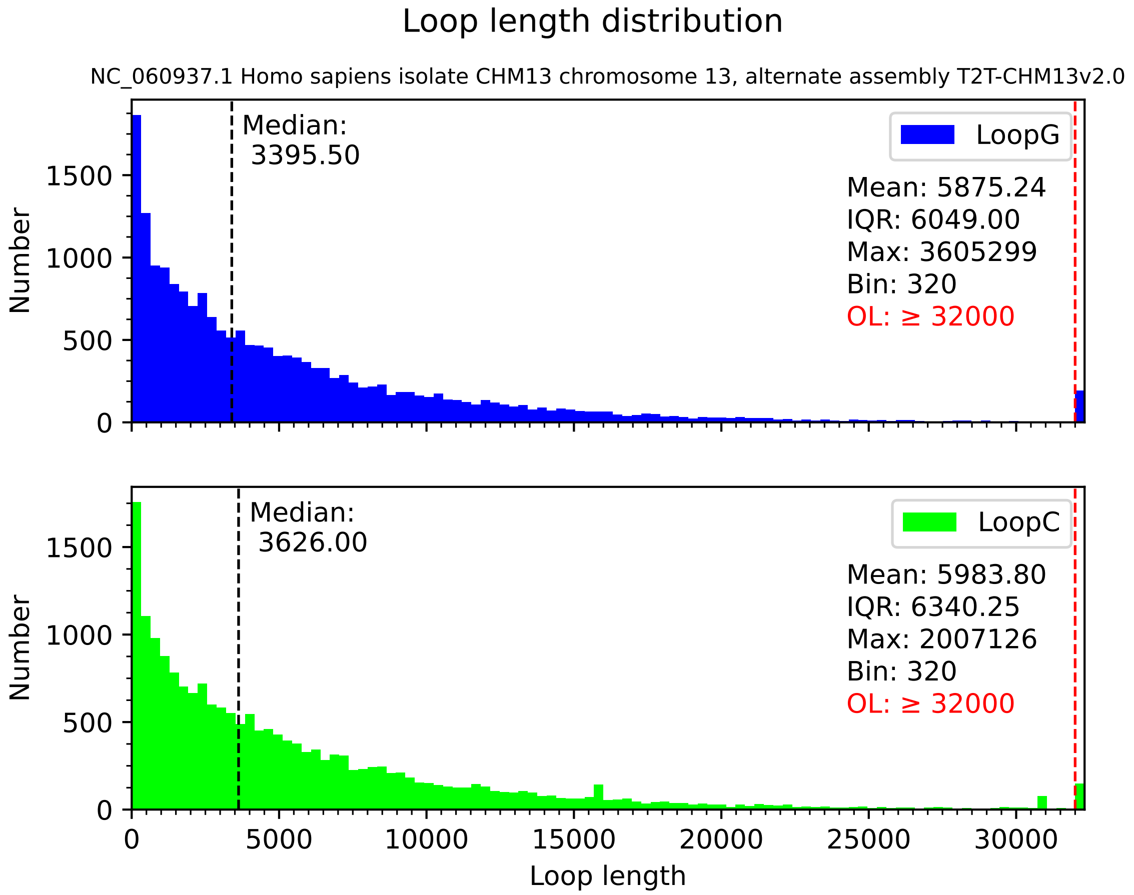

Loop length distribution
NC_060937.1 Homo sapiens isolate CHM13 chromosome 13, alternate assembly T2T-CHM13v2.0
Median:
3395.50
LoopG
Mean: 5875.24
IQR: 6049.00
Max: 3605299
Bin: 320
OL: ≥ 32000
Number
0
500
1000
1500
Median:
3626.00
LoopC
Mean: 5983.80
IQR: 6340.25
Max: 2007126
Bin: 320
OL: ≥ 32000
Number
0
500
1000
1500
0
5000
10000
15000
20000
25000
30000
Loop length

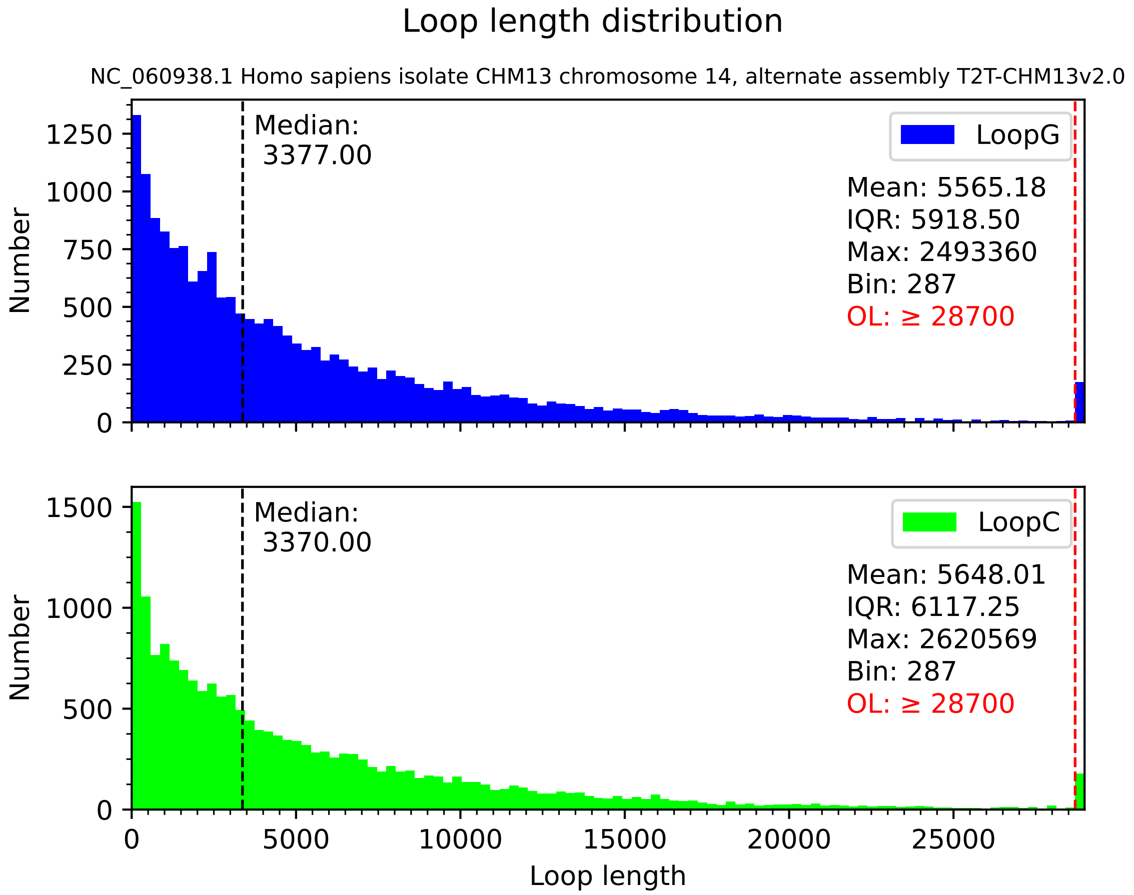
Loop length distribution
NC_060938.1 Homo sapiens isolate CHM13 chromosome 14, alternate assembly T2T-CHM13v2.0
Median:
3377.00
LoopG
Mean: 5565.18
IQR: 5918.50
Max: 2493360
Bin: 287
OL: ≥ 28700
Number
0
250
500
750
1000
1250
Median:
3370.00
LoopC
Mean: 5648.01
IQR: 6117.25
Max: 2620569
Bin: 287
OL: ≥ 28700
Number
0
500
1000
1500
0
5000
10000
15000
20000
25000
Loop length

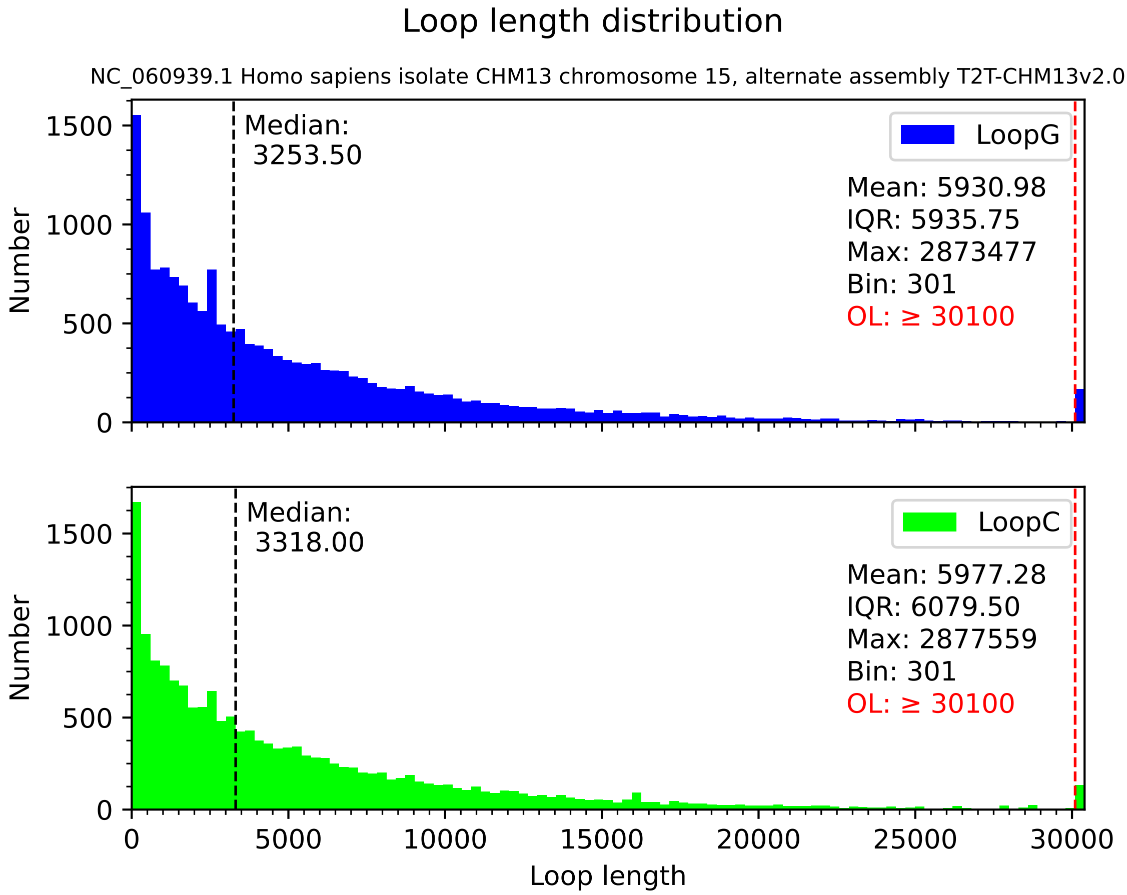

Loop length distribution
NC_060939.1 Homo sapiens isolate CHM13 chromosome 15, alternate assembly T2T-CHM13v2.0
Median: 3253.50
LoopG
Mean: 5930.98
IQR: 5935.75
Max: 2873477
Bin: 301
OL: ≥ 30100
Median: 3318.00
LoopC
Mean: 5977.28
IQR: 6079.50
Max: 2877559
Bin: 301
OL: ≥ 30100
Number
Loop length
0
500
1000
1500
5000
10000
15000
20000
25000
30000

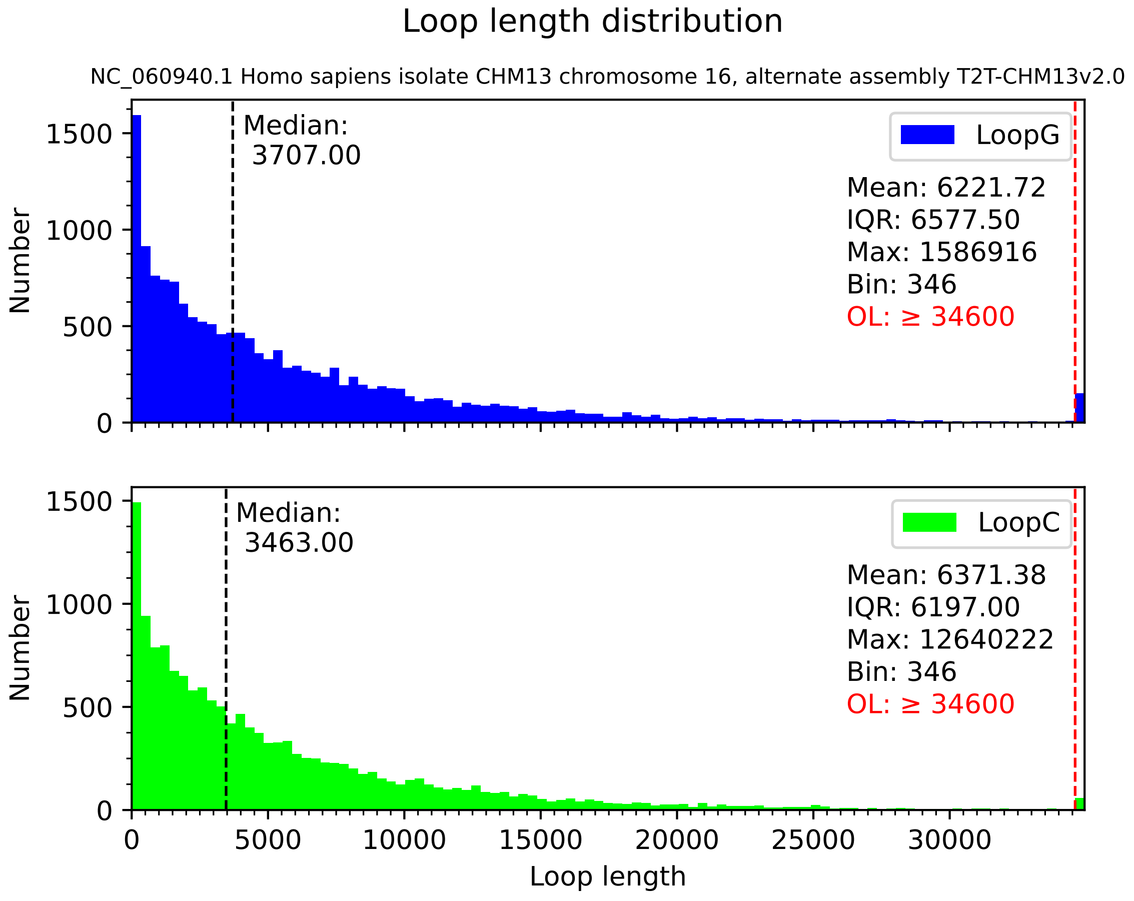
Loop length distribution
NC_060940.1 Homo sapiens isolate CHM13 chromosome 16, alternate assembly T2T-CHM13v2.0
Median:
3707.00
LoopG
Mean: 6221.72
IQR: 6577.50
Max: 1586916
Bin: 346
OL: ≥ 34600
Median:
3463.00
LoopC
Mean: 6371.38
IQR: 6197.00
Max: 12640222
Bin: 346
OL: ≥ 34600
Number
Loop length
0
500
1000
1500
0
5000
10000
15000
20000
25000
30000

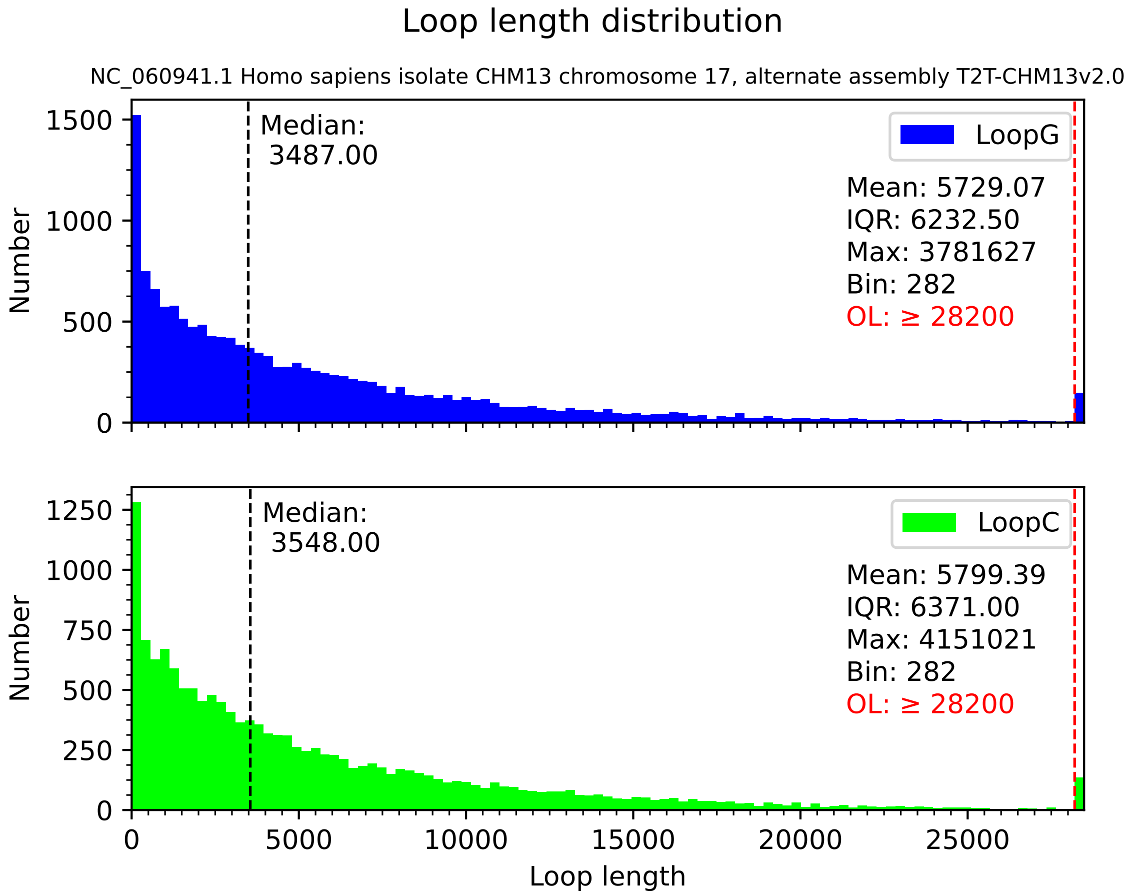

Loop length distribution
NC_060941.1 Homo sapiens isolate CHM13 chromosome 17, alternate assembly T2T-CHM13v2.0
Median:
3487.00
LoopG
Mean: 5729.07
IQR: 6232.50
Max: 3781627
Bin: 282
OL: ≥ 28200
Number
0
500
1000
1500
Median:
3548.00
LoopC
Mean: 5799.39
IQR: 6371.00
Max: 4151021
Bin: 282
OL: ≥ 28200
Number
0
250
500
750
1000
1250
0
5000
10000
15000
20000
25000
Loop length

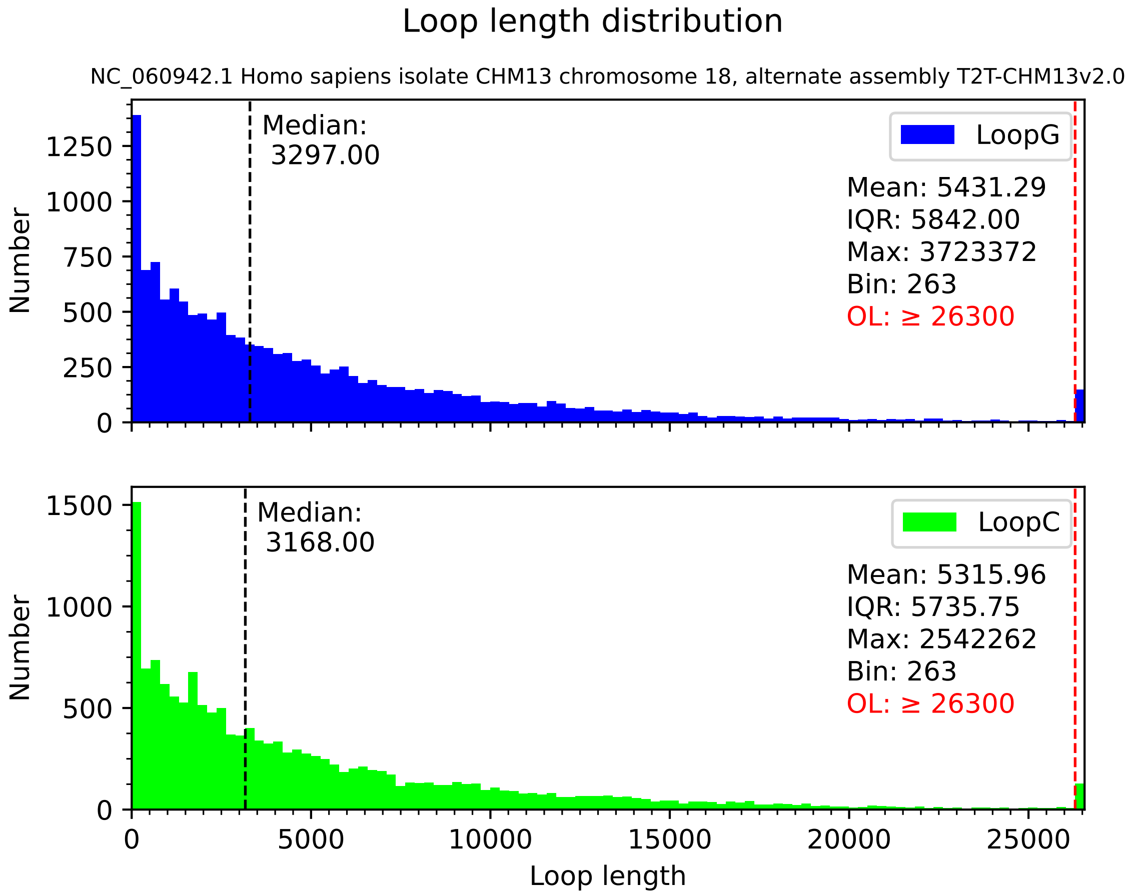
Loop length distribution
NC_060942.1 Homo sapiens isolate CHM13 chromosome 18, alternate assembly T2T-CHM13v2.0
Median:
3297.00
LoopG
Mean: 5431.29
IQR: 5842.00
Max: 3723372
Bin: 263
OL: ≥ 26300
Number
Median:
3168.00
LoopC
Mean: 5315.96
IQR: 5735.75
Max: 2542262
Bin: 263
OL: ≥ 26300
Number
Loop length

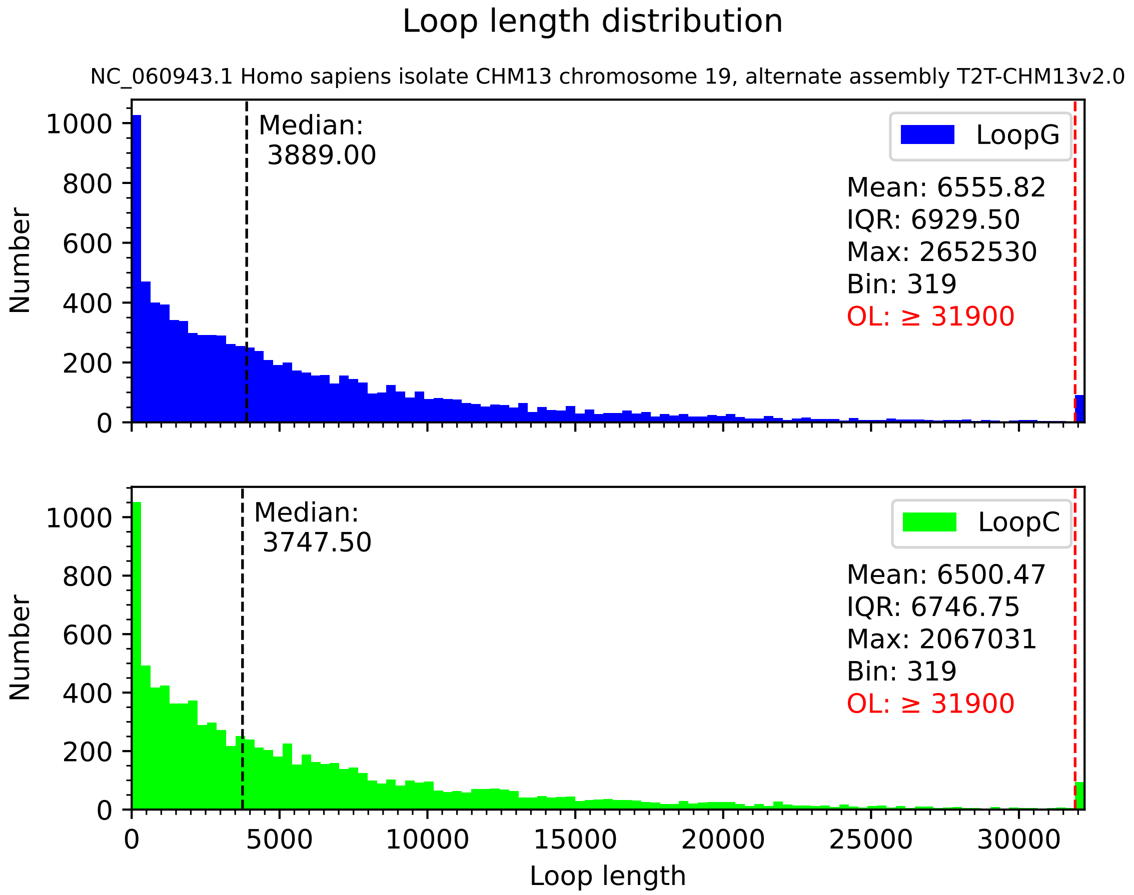

Loop length distribution
NC_060943.1 Homo sapiens isolate CHM13 chromosome 19, alternate assembly T2T-CHM13v2.0
Median:
3889.00
LoopG
Mean: 6555.82
IQR: 6929.50
Max: 2652530
Bin: 319
OL: ≥ 31900
Median:
3747.50
LoopC
Mean: 6500.47
IQR: 6746.75
Max: 2067031
Bin: 319
OL: ≥ 31900
Number
Loop length
0
200
400
600
800
1000
0
5000
10000
15000
20000
25000
30000

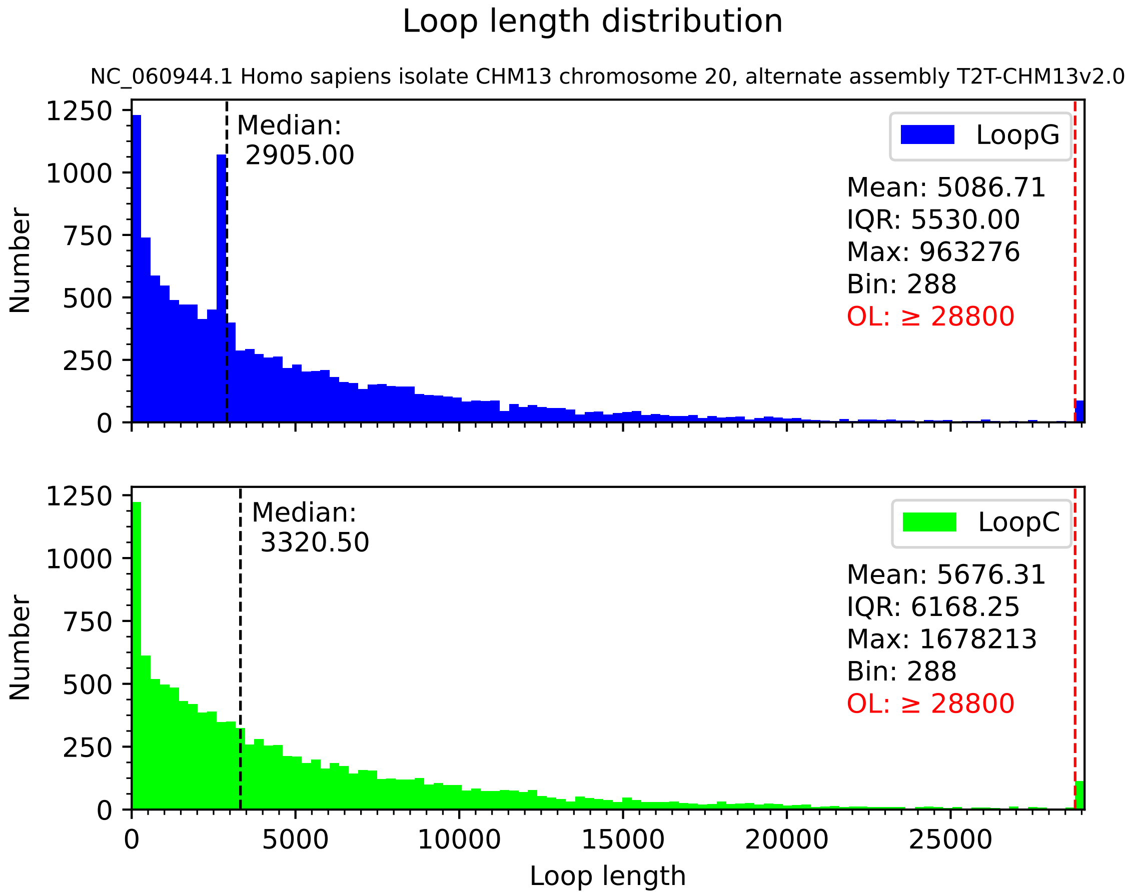

Loop length distribution
NC_060944.1 Homo sapiens isolate CHM13 chromosome 20, alternate assembly T2T-CHM13v2.0
Median:
2905.00
LoopG
Mean: 5086.71
IQR: 5530.00
Max: 963276
Bin: 288
OL: ≥ 28800
Number
0
250
500
750
1000
1250
Median:
3320.50
LoopC
Mean: 5676.31
IQR: 6168.25
Max: 1678213
Bin: 288
OL: ≥ 28800
Number
0
250
500
750
1000
1250
0
5000
10000
15000
20000
25000
Loop length

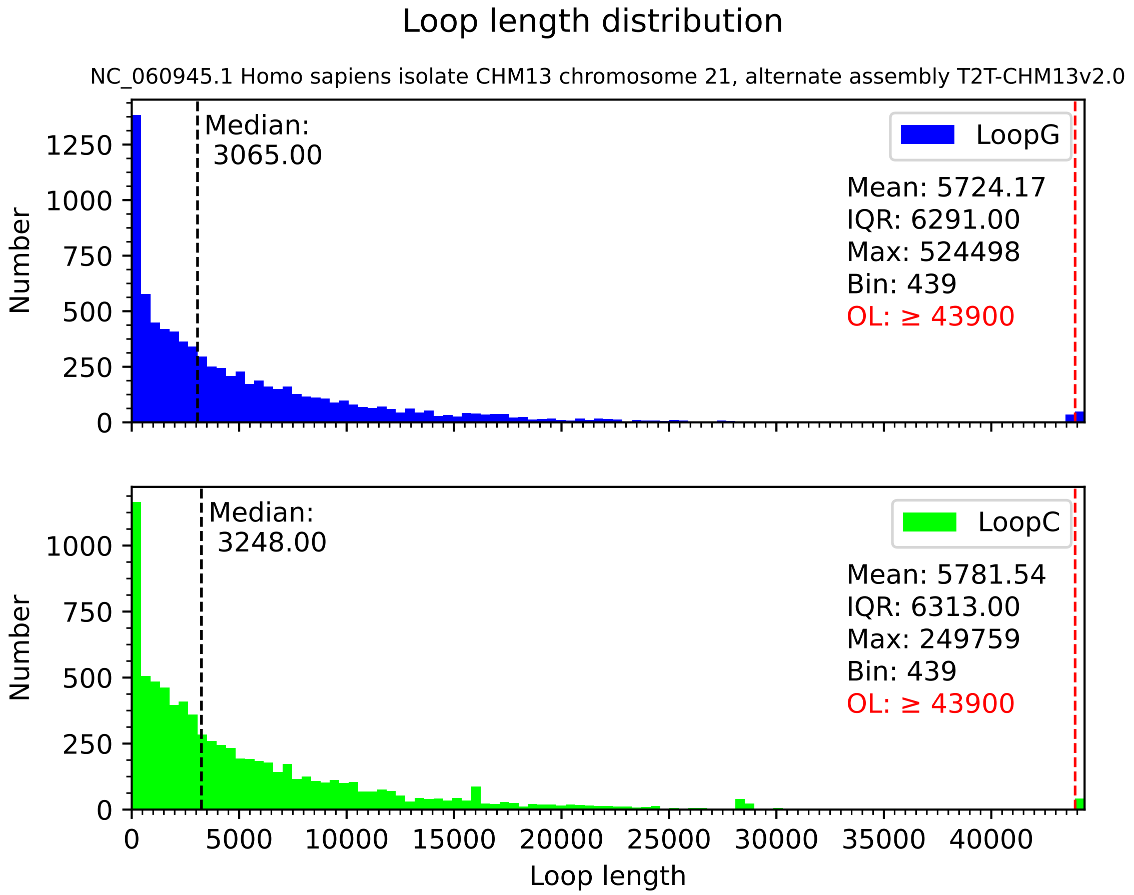

Loop length distribution
NC_060945.1 Homo sapiens isolate CHM13 chromosome 21, alternate assembly T2T-CHM13v2.0
Median:
3065.00
LoopG
Mean: 5724.17
IQR: 6291.00
Max: 524498
Bin: 439
OL: ≥ 43900
Number
0
250
500
750
1000
1250
Median:
3248.00
LoopC
Mean: 5781.54
IQR: 6313.00
Max: 249759
Bin: 439
OL: ≥ 43900
Number
0
250
500
750
1000
0
5000
10000
15000
20000
25000
30000
35000
40000
Loop length

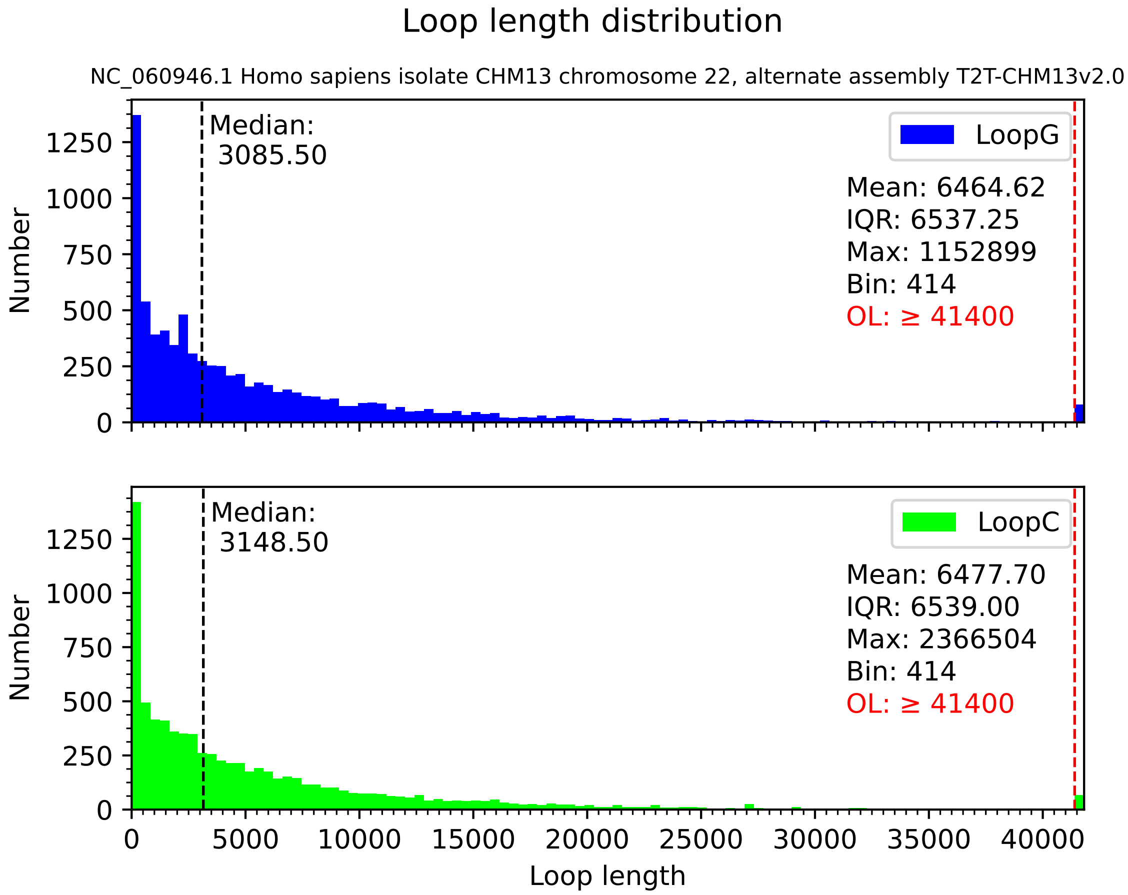
Loop length distribution
NC_060946.1 Homo sapiens isolate CHM13 chromosome 22, alternate assembly T2T-CHM13v2.0
Median:
3085.50
LoopG
Mean: 6464.62
IQR: 6537.25
Max: 1152899
Bin: 414
OL: ≥ 41400
Number
0
250
500
750
1000
1250
Median:
3148.50
LoopC
Mean: 6477.70
IQR: 6539.00
Max: 2366504
Bin: 414
OL: ≥ 41400
Number
0
250
500
750
1000
1250
0
5000
10000
15000
20000
25000
30000
35000
40000
Loop length

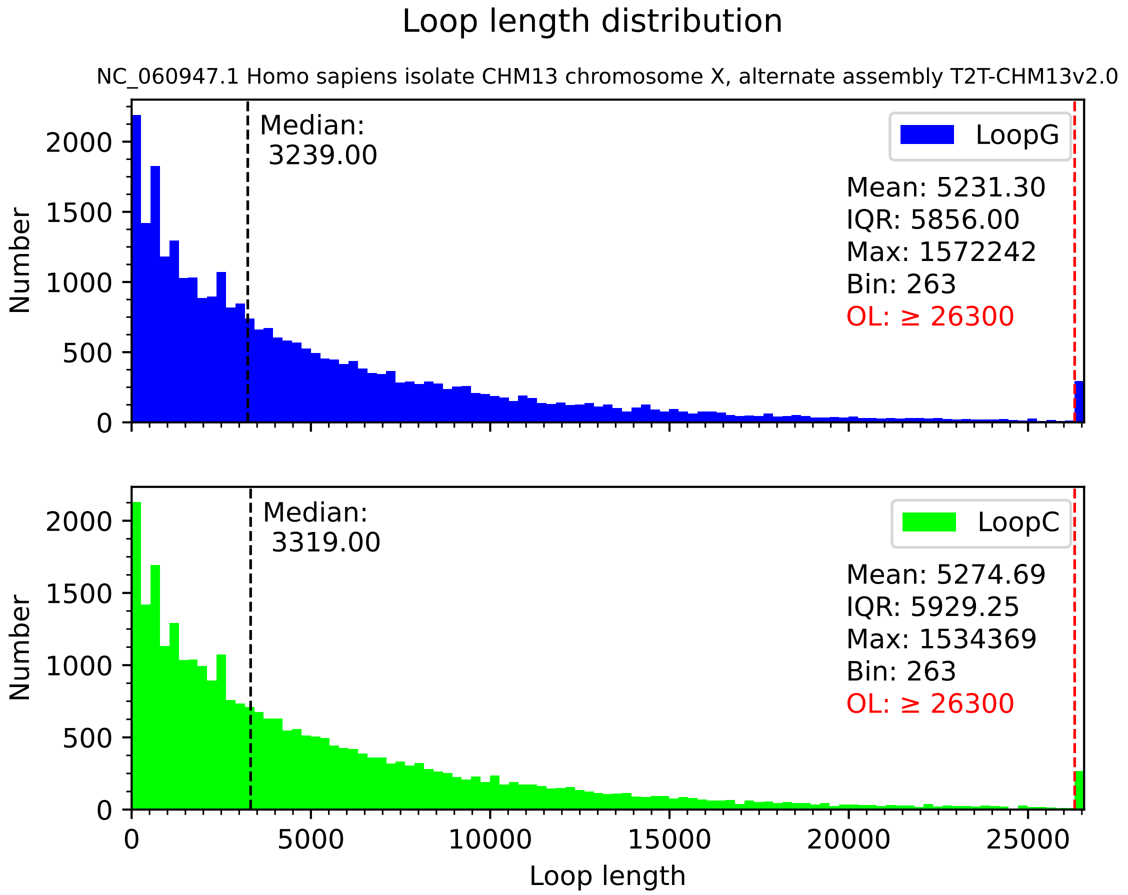

Loop length distribution
NC_060947.1 Homo sapiens isolate CHM13 chromosome X, alternate assembly T2T-CHM13v2.0
Median:
3239.00
LoopG
Mean: 5231.30
IQR: 5856.00
Max: 1572242
Bin: 263
OL: ≥ 26300
Median:
3319.00
LoopC
Mean: 5274.69
IQR: 5929.25
Max: 1534369
Bin: 263
OL: ≥ 26300
Number
Loop length
0
500
1000
1500
2000
0
5000
10000
15000
20000
25000

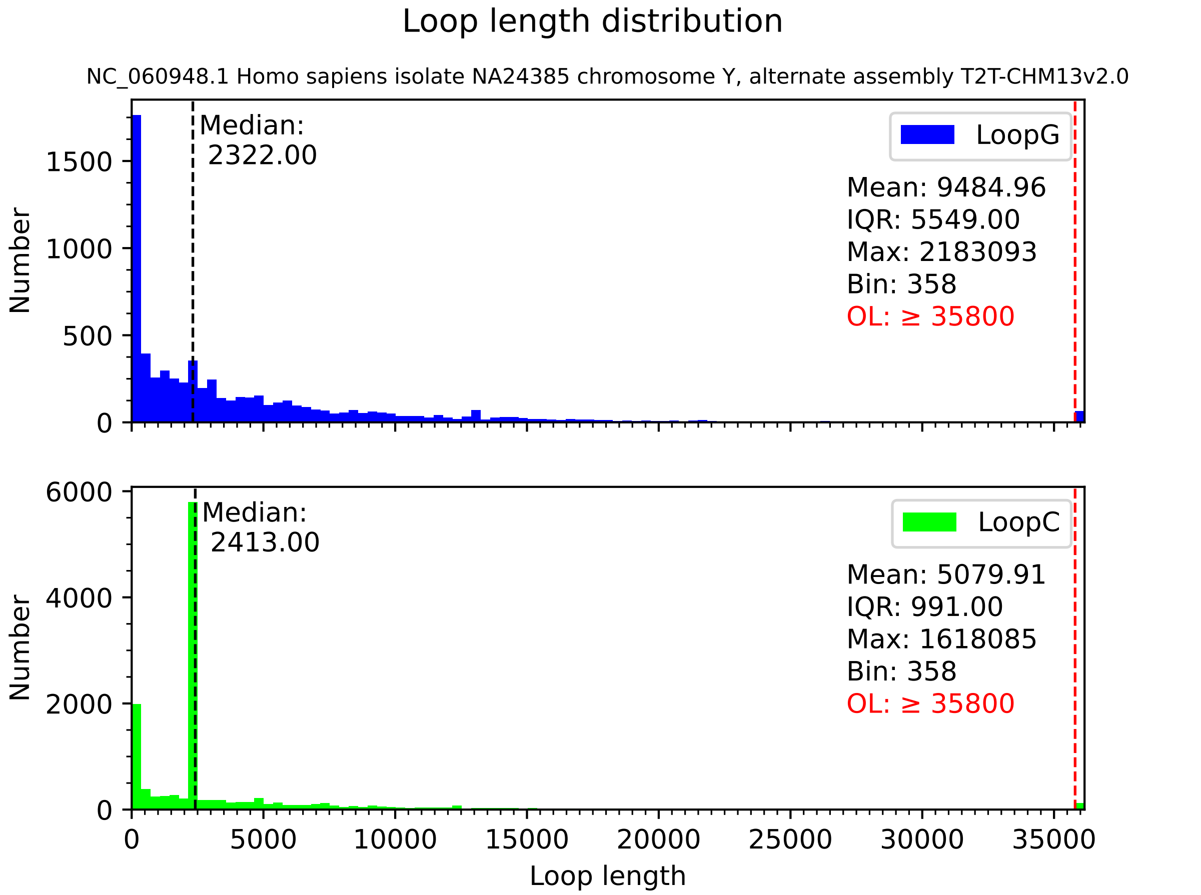
Loop length distribution
NC_060948.1 Homo sapiens isolate NA24385 chromosome Y, alternate assembly T2T-CHM13v2.0
LoopG
Median:
2322.00
Mean: 9484.96
IQR: 5549.00
Max: 2183093
Bin: 358
OL: ≥ 35800
Number
0
500
1000
1500
LoopC
Median:
2413.00
Mean: 5079.91
IQR: 991.00
Max: 1618085
Bin: 358
OL: ≥ 35800
Number
0
2000
4000
6000
0
5000
10000
15000
20000
25000
30000
35000
Loop length
