## Supplementary file 2 for "FuzzyClusTeR: a web server for analysis of tandem and diffuse DNA repeat clusters with application to telomeric-like repeats"

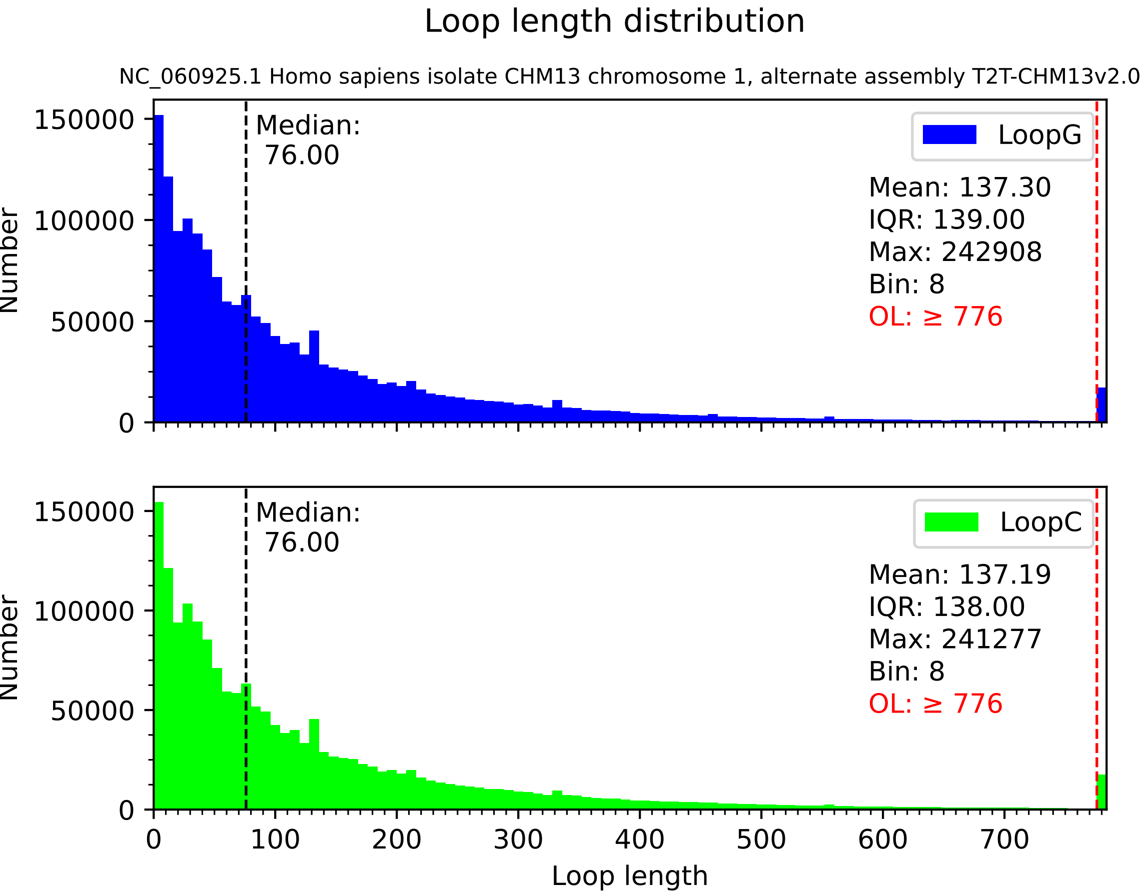
Loop length distribution
NC_060925.1 Homo sapiens isolate CHM13 chromosome 1, alternate assembly T2T-CHM13v2.0
Median:
76.00
LoopG
Mean: 137.30
IQR: 139.00
Max: 242908
Bin: 8
OL: ≥ 776
Median:
76.00
LoopC
Mean: 137.19
IQR: 138.00
Max: 241277
Bin: 8
OL: ≥ 776
Number
Loop length
0
50000
100000
150000
0
100
200
300
400
500
600
700

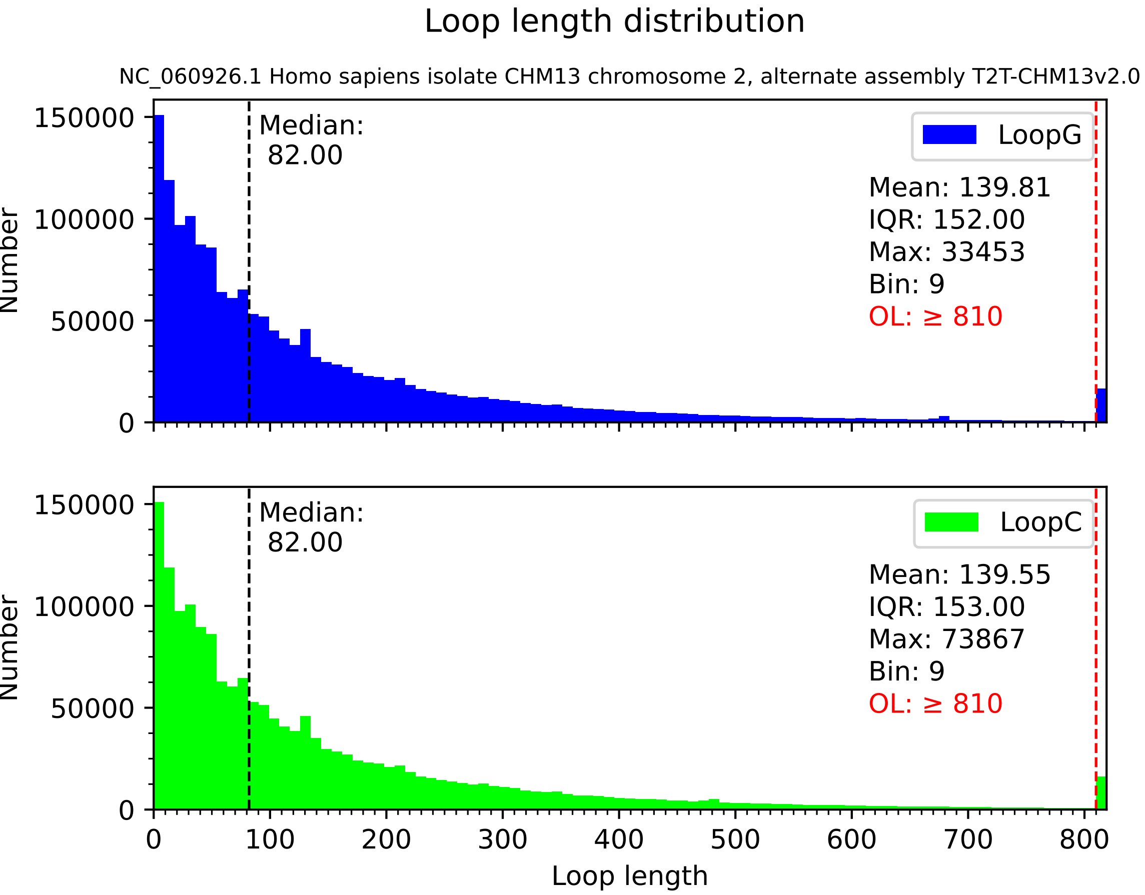

Loop length distribution
NC_060926.1 Homo sapiens isolate CHM13 chromosome 2, alternate assembly T2T-CHM13v2.0
Median:
82.00
LoopG
Mean: 139.81
IQR: 152.00
Max: 33453
Bin: 9
OL: ≥ 810
Median:
82.00
LoopC
Mean: 139.55
IQR: 153.00
Max: 73867
Bin: 9
OL: ≥ 810
Number
Number
Loop length
0
50000
100000
150000
0
50000
100000
150000
0
100
200
300
400
500
600
700
800

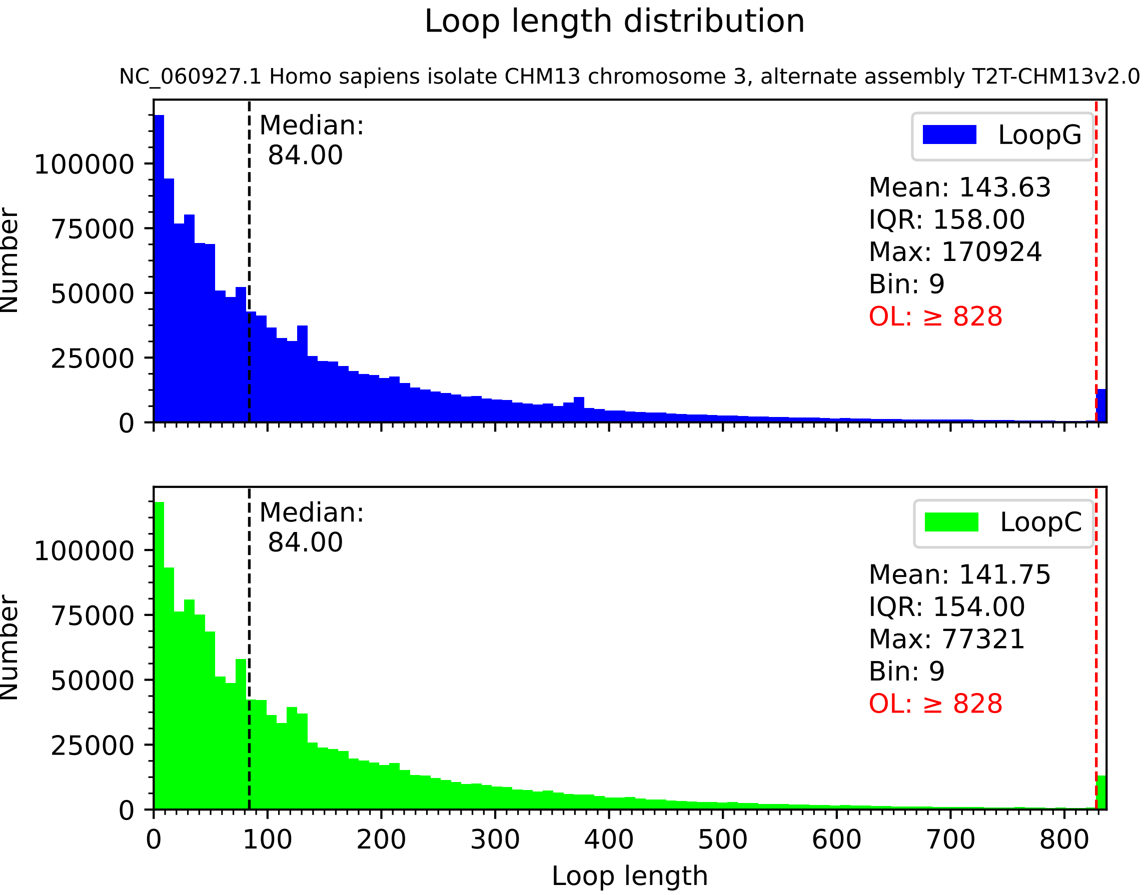

Loop length distribution
NC_060927.1 Homo sapiens isolate CHM13 chromosome 3, alternate assembly T2T-CHM13v2.0
Median:
84.00
LoopG
Mean: 143.63
IQR: 158.00
Max: 170924
Bin: 9
OL: ≥ 828
Median:
84.00
LoopC
Mean: 141.75
IQR: 154.00
Max: 77321
Bin: 9
OL: ≥ 828
Number
Loop length
0
25000
50000
75000
100000
100
200
300
400
500
600
700
800

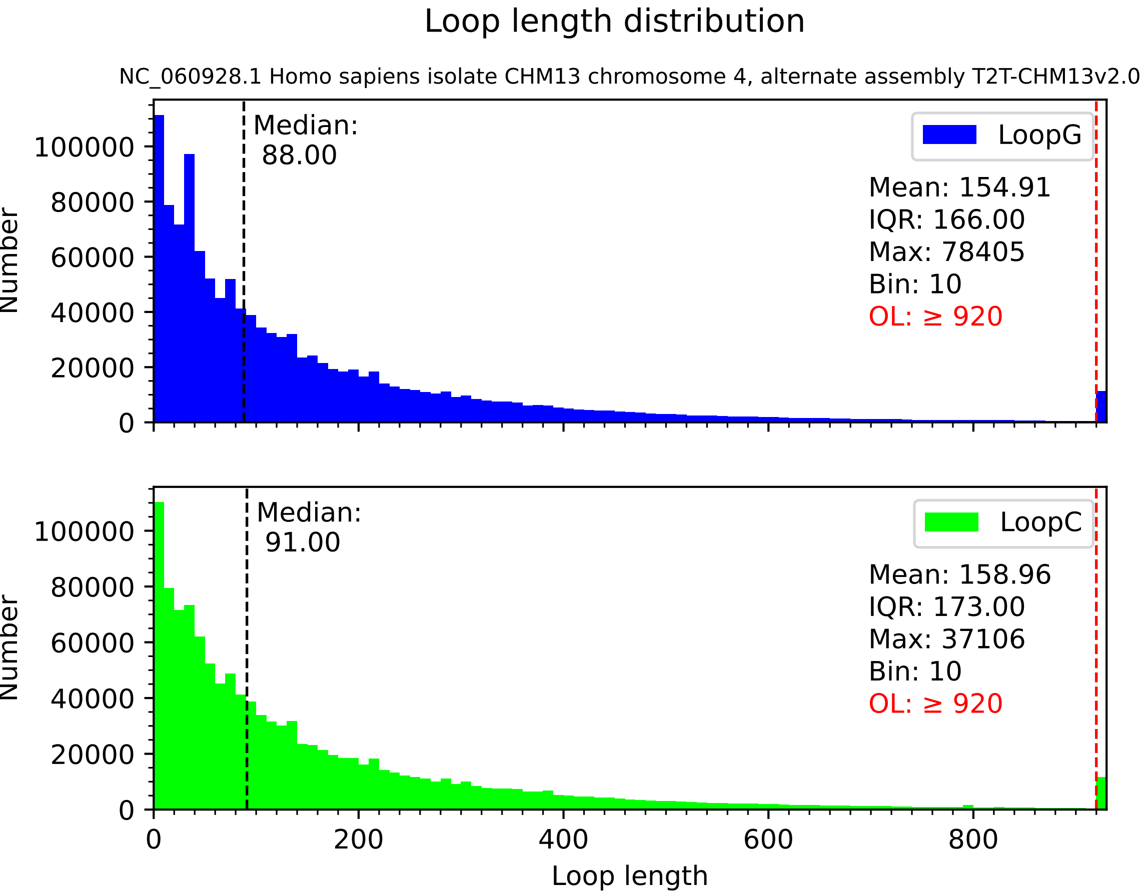
Loop length distribution
NC_060928.1 Homo sapiens isolate CHM13 chromosome 4, alternate assembly T2T-CHM13v2.0
Median:
88.00
LoopG
Mean: 154.91
IQR: 166.00
Max: 78405
Bin: 10
OL: ≥ 920
Number
Median:
91.00
LoopC
Mean: 158.96
IQR: 173.00
Max: 37106
Bin: 10
OL: ≥ 920
Number
Loop length
0
20000
40000
60000
80000
100000
0
200
400
600
800

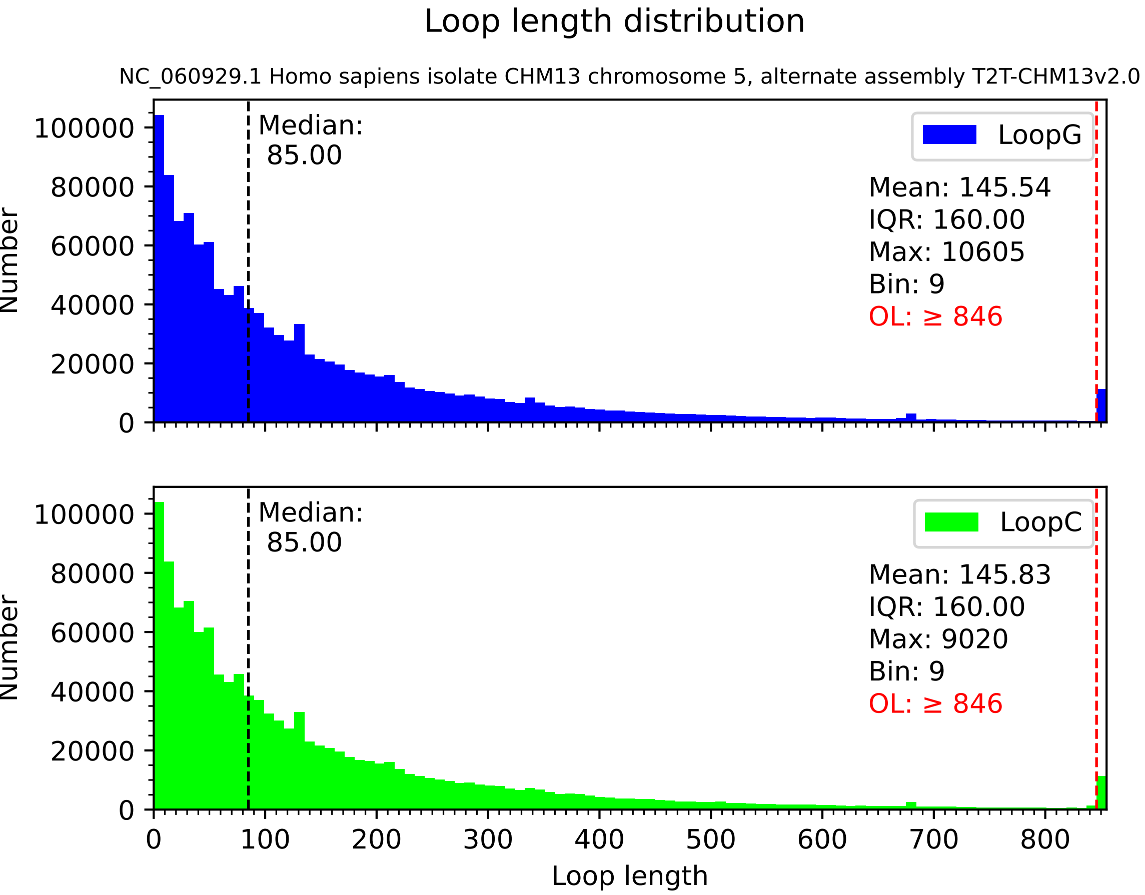

Loop length distribution
NC_060929.1 Homo sapiens isolate CHM13 chromosome 5, alternate assembly T2T-CHM13v2.0
Median: 85.00
LoopG
Mean: 145.54
IQR: 160.00
Max: 10605
Bin: 9
OL: ≥ 846
Number
0
20000
40000
60000
80000
100000
Median: 85.00
LoopC
Mean: 145.83
IQR: 160.00
Max: 9020
Bin: 9
OL: ≥ 846
Number
0
100
200
300
400
500
600
700
800
Loop length

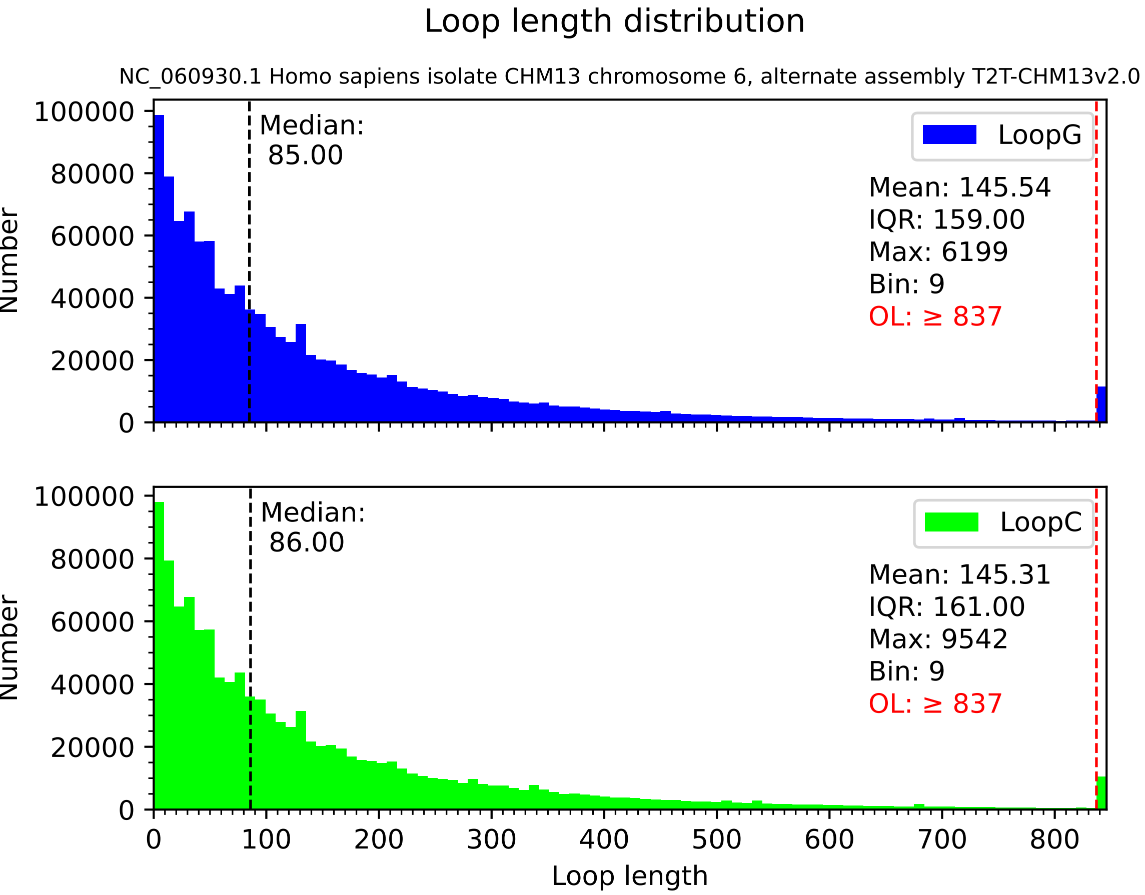

Loop length distribution
NC_060930.1 Homo sapiens isolate CHM13 chromosome 6, alternate assembly T2T-CHM13v2.0
Number
100000
80000
60000
40000
20000
0
Median:
85.00
LoopG
Mean: 145.54
IQR: 159.00
Max: 6199
Bin: 9
OL: ≥ 837
Number
100000
80000
60000
40000
20000
0
Median:
86.00
LoopC
Mean: 145.31
IQR: 161.00
Max: 9542
Bin: 9
OL: ≥ 837
0
100
200
300
400
500
600
700
800
Loop length

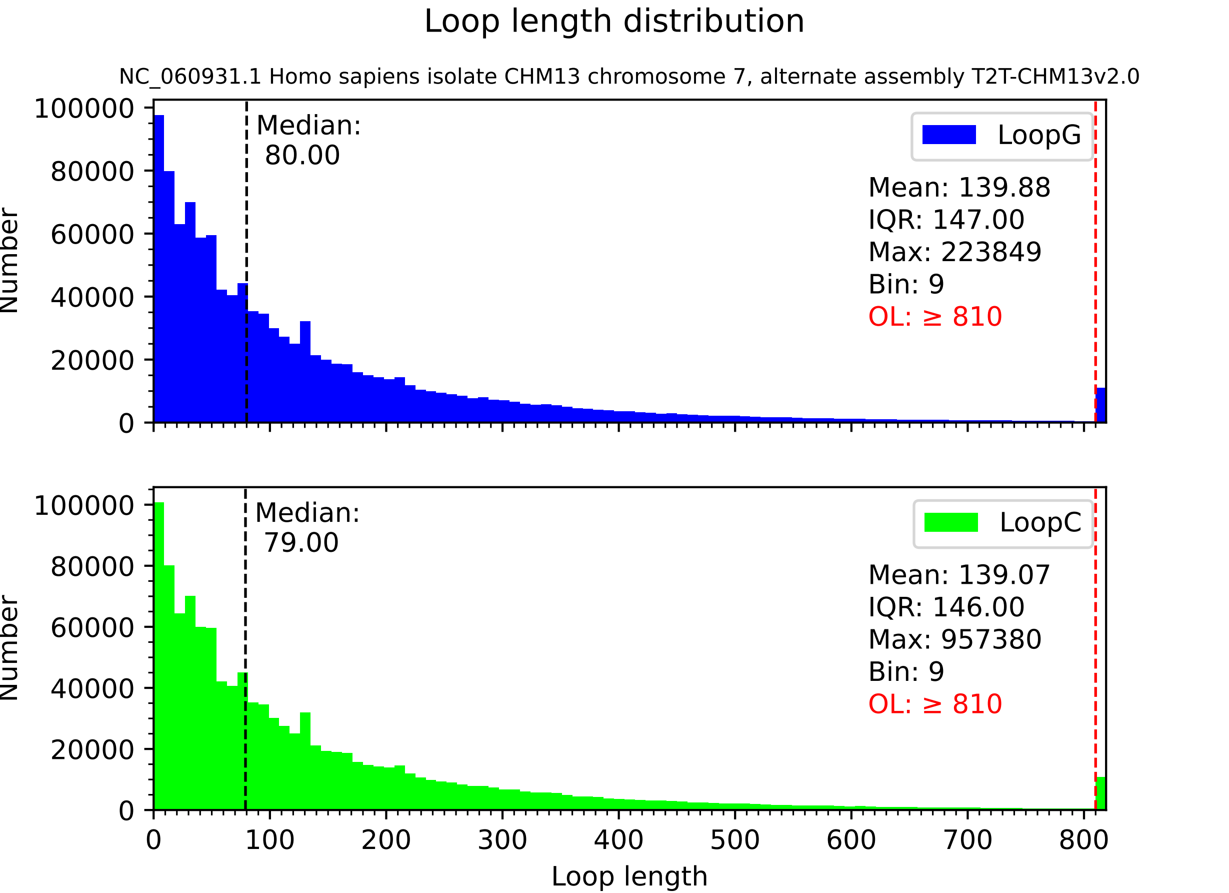

Loop length distribution
NC_060931.1 Homo sapiens isolate CHM13 chromosome 7, alternate assembly T2T-CHM13v2.0
Number
Median:
80.00
LoopG
Mean: 139.88
IQR: 147.00
Max: 223849
Bin: 9
OL: ≥ 810
Number
Median:
79.00
LoopC
Mean: 139.07
IQR: 146.00
Max: 957380
Bin: 9
OL: ≥ 810
Loop length

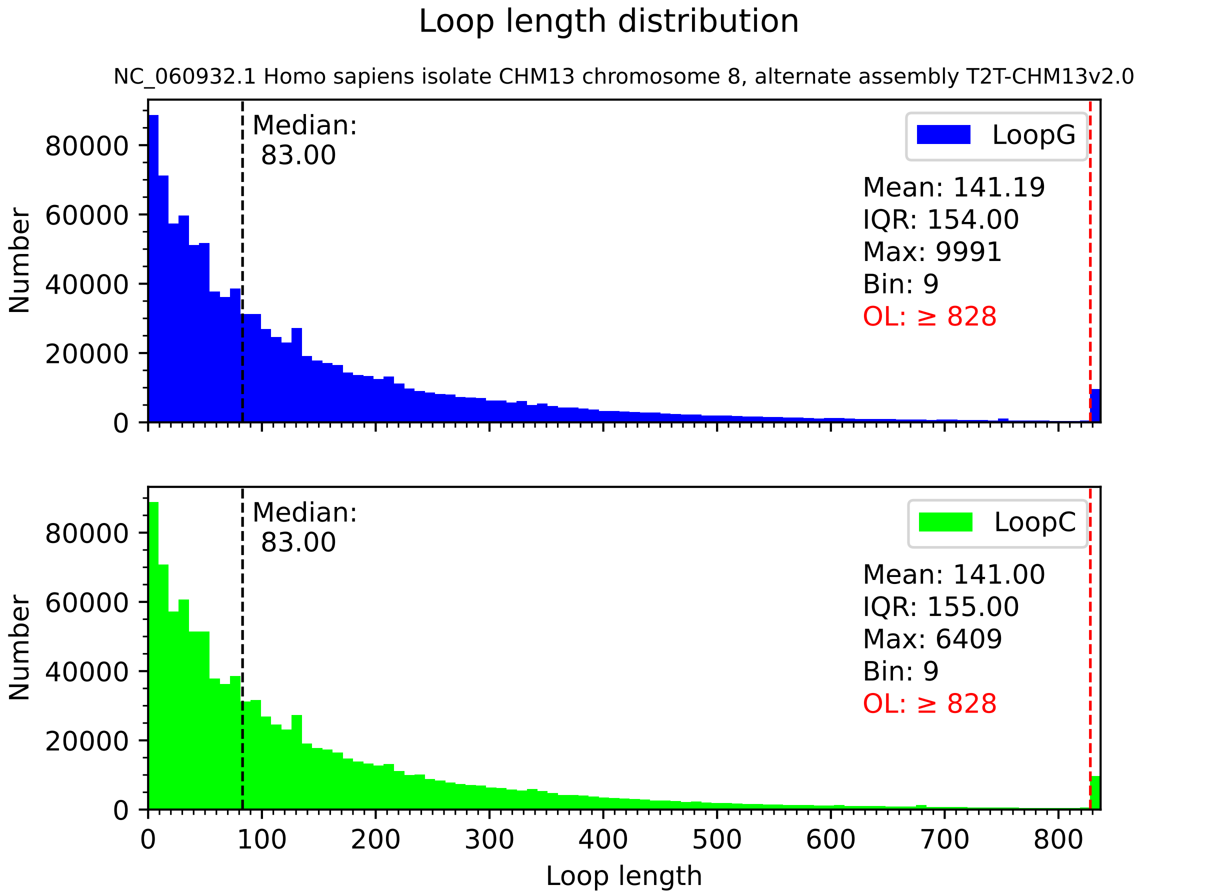
Loop length distribution
NC_060932.1 Homo sapiens isolate CHM13 chromosome 8, alternate assembly T2T-CHM13v2.0
Median:
83.00
LoopG
Mean: 141.19
IQR: 154.00
Max: 9991
Bin: 9
OL: ≥ 828
Median:
83.00
LoopC
Mean: 141.00
IQR: 155.00
Max: 6409
Bin: 9
OL: ≥ 828
Number
0
20000
40000
60000
80000
0
100
200
300
400
500
600
700
800
Loop length

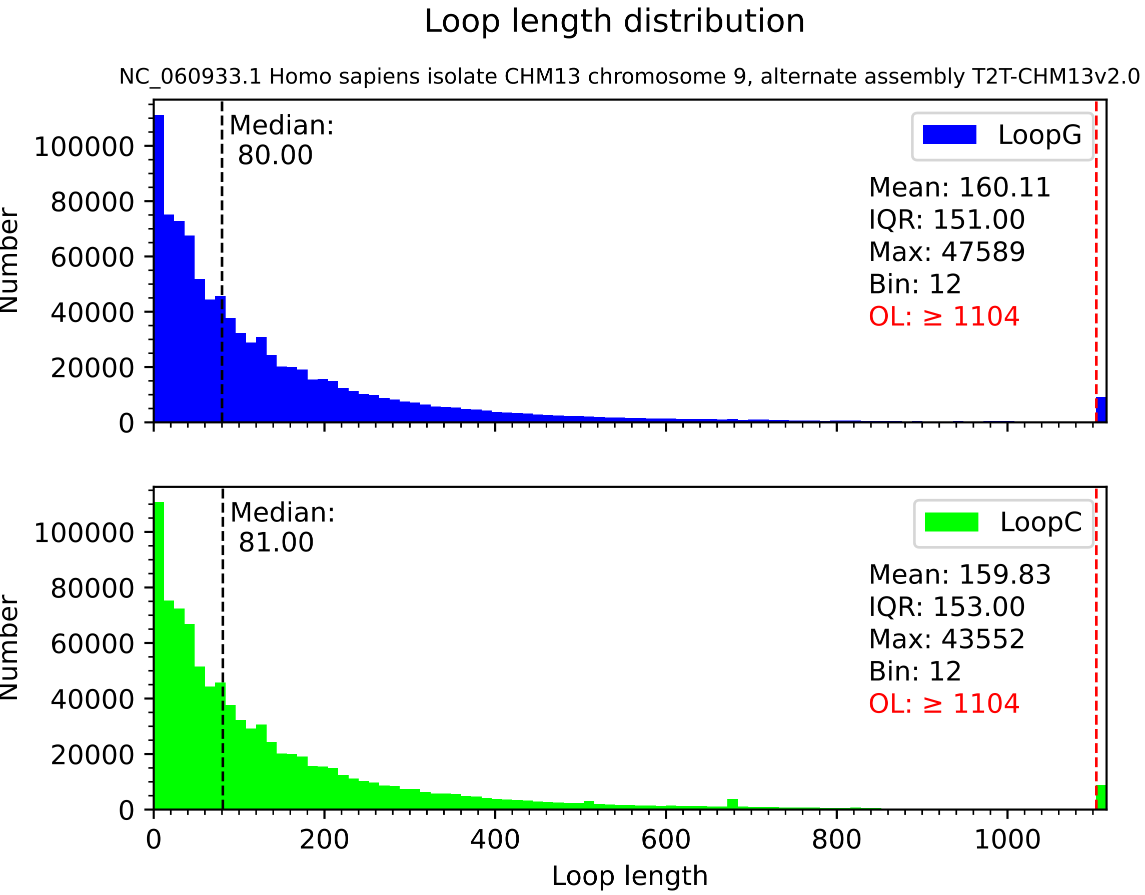

Loop length distribution
NC_060933.1 Homo sapiens isolate CHM13 chromosome 9, alternate assembly T2T-CHM13v2.0
Median:
80.00
LoopG
Mean: 160.11
IQR: 151.00
Max: 47589
Bin: 12
OL: ≥ 1104
Median:
81.00
LoopC
Mean: 159.83
IQR: 153.00
Max: 43552
Bin: 12
OL: ≥ 1104
Number
Loop length
0
20000
40000
60000
80000
100000
0
200
400
600
800
1000

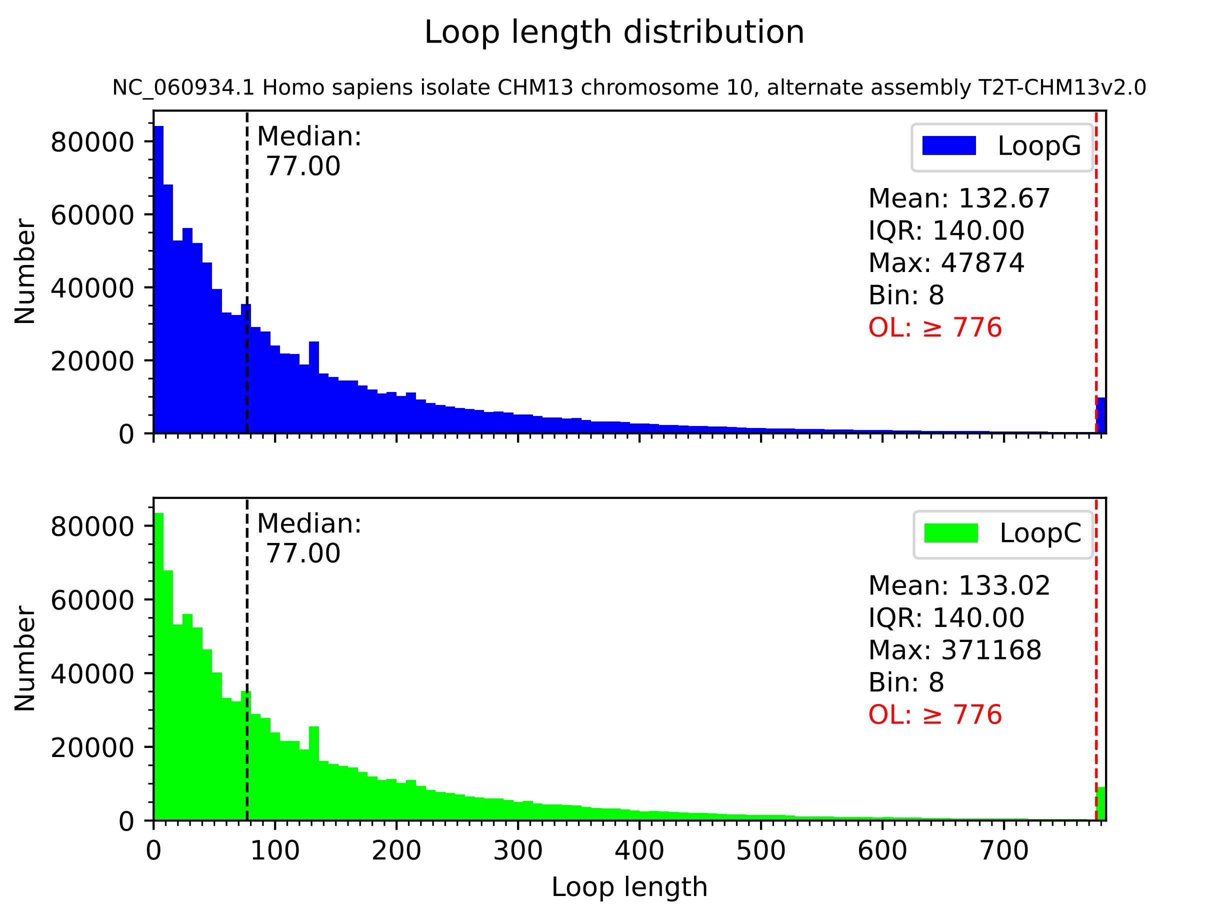

Loop length distribution
NC_060934.1 Homo sapiens isolate CHM13 chromosome 10, alternate assembly T2T-CHM13v2.0
Number
Median:
77.00
LoopG
Mean: 132.67
IQR: 140.00
Max: 47874
Bin: 8
OL: ≥ 776
Median:
77.00
LoopC
Mean: 133.02
IQR: 140.00
Max: 371168
Bin: 8
OL: ≥ 776
0
20000
40000
60000
80000
0
100
200
300
400
500
600
700
Loop length

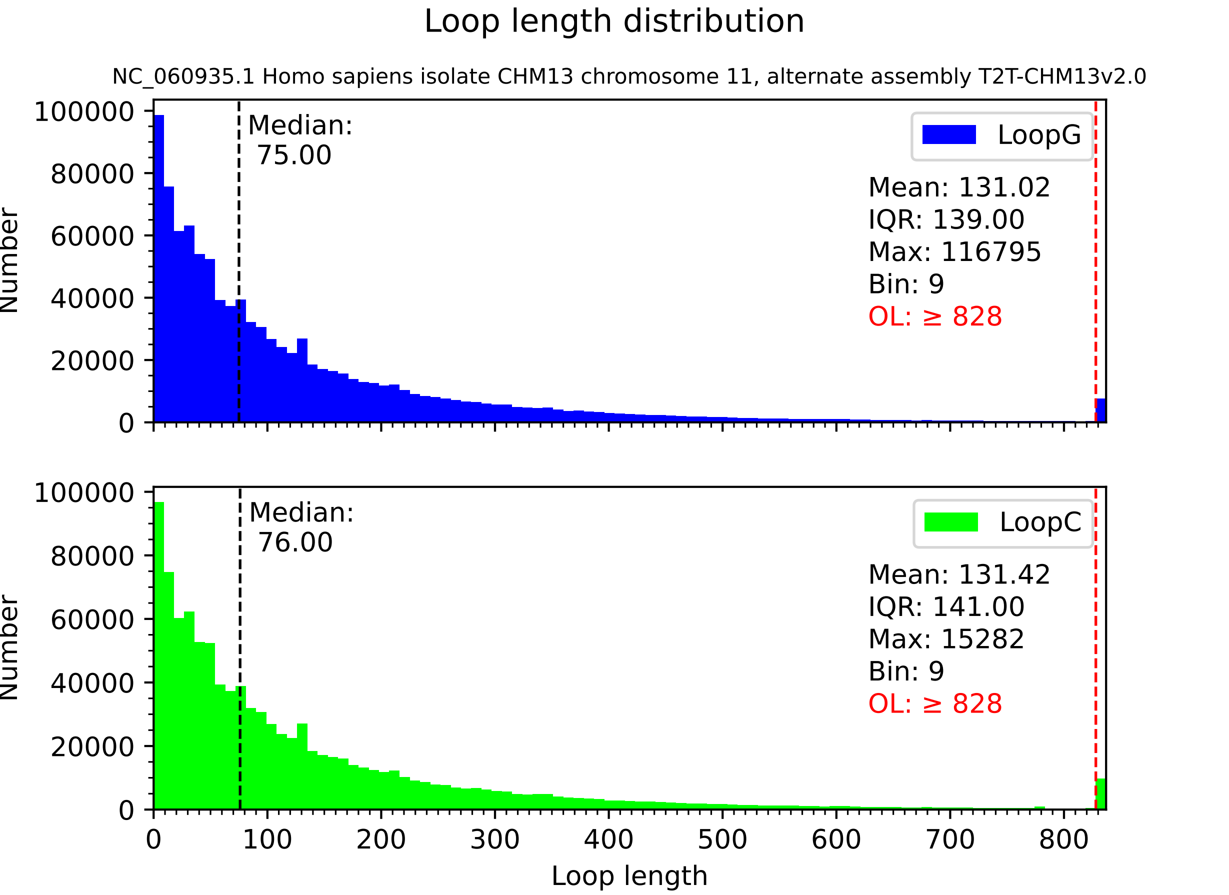

Loop length distribution
NC_060935.1 Homo sapiens isolate CHM13 chromosome 11, alternate assembly T2T-CHM13v2.0
Median:
75.00
LoopG
Mean: 131.02
IQR: 139.00
Max: 116795
Bin: 9
OL: ≥ 828
Median:
76.00
LoopC
Mean: 131.42
IQR: 141.00
Max: 15282
Bin: 9
OL: ≥ 828
Number
Loop length
0
20000
40000
60000
80000
100000
0
100
200
300
400
500
600
700
800

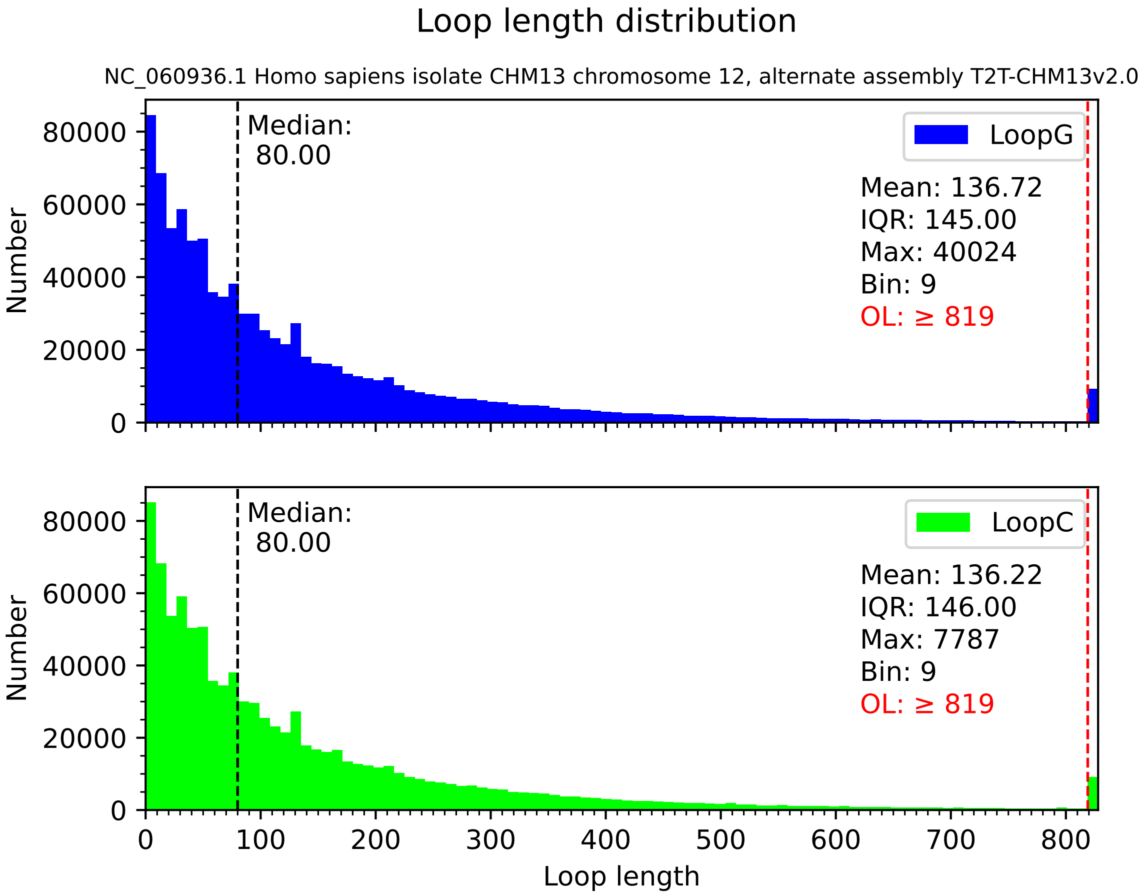

Loop length distribution
NC_060936.1 Homo sapiens isolate CHM13 chromosome 12, alternate assembly T2T-CHM13v2.0
Median:
80.00
LoopG
Mean: 136.72
IQR: 145.00
Max: 40024
Bin: 9
OL: ≥ 819
Median:
80.00
LoopC
Mean: 136.22
IQR: 146.00
Max: 7787
Bin: 9
OL: ≥ 819
Number
Loop length
0
20000
40000
60000
80000
0
100
200
300
400
500
600
700
800

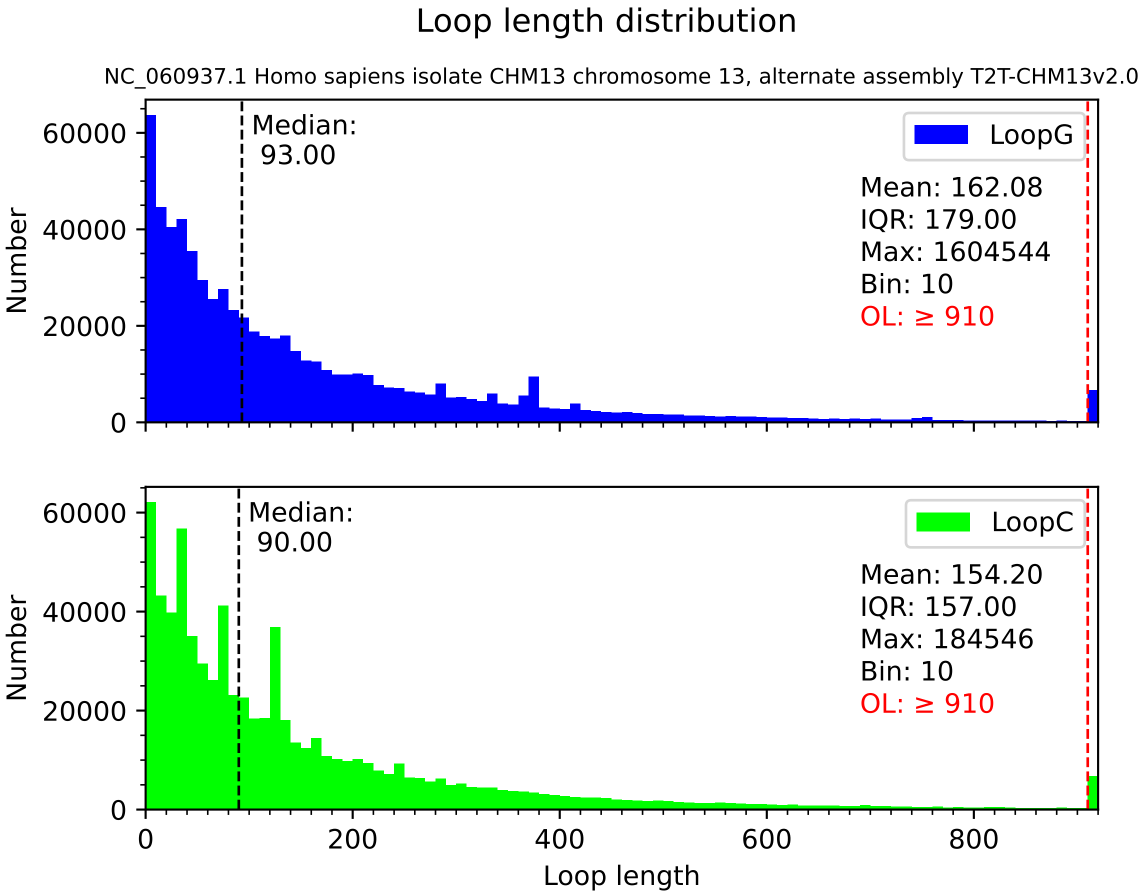

Loop length distribution
NC_060937.1 Homo sapiens isolate CHM13 chromosome 13, alternate assembly T2T-CHM13v2.0
Median:
93.00
LoopG
Mean: 162.08
IQR: 179.00
Max: 1604544
Bin: 10
OL: ≥ 910
Number
0
20000
40000
60000
Median:
90.00
LoopC
Mean: 154.20
IQR: 157.00
Max: 184546
Bin: 10
OL: ≥ 910
0
200
400
600
800
Loop length

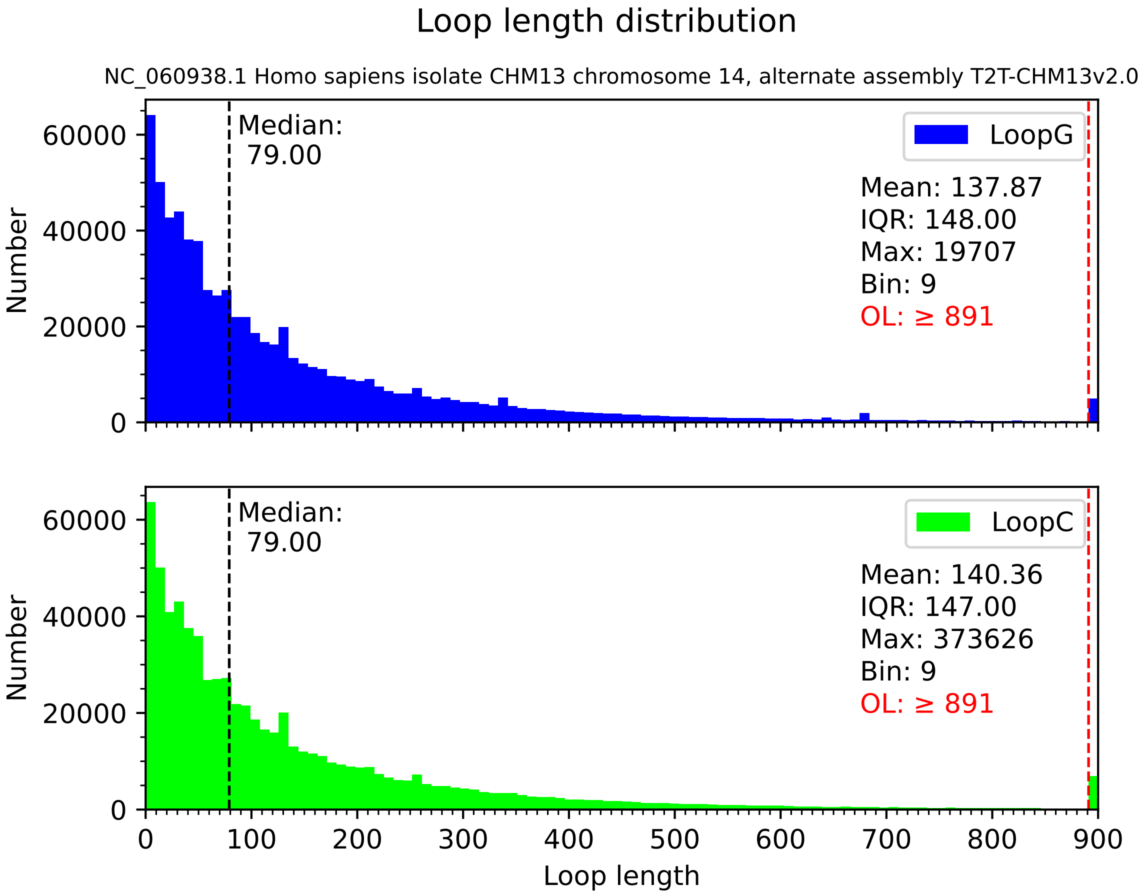

Loop length distribution
NC_060938.1 Homo sapiens isolate CHM13 chromosome 14, alternate assembly T2T-CHM13v2.0
Median: 79.00
LoopG
Mean: 137.87
IQR: 148.00
Max: 19707
Bin: 9
OL: ≥ 891
Number
0
20000
40000
60000
Median: 79.00
LoopC
Mean: 140.36
IQR: 147.00
Max: 373626
Bin: 9
OL: ≥ 891
Number
0
20000
40000
60000
0
100
200
300
400
500
600
700
800
900
Loop length

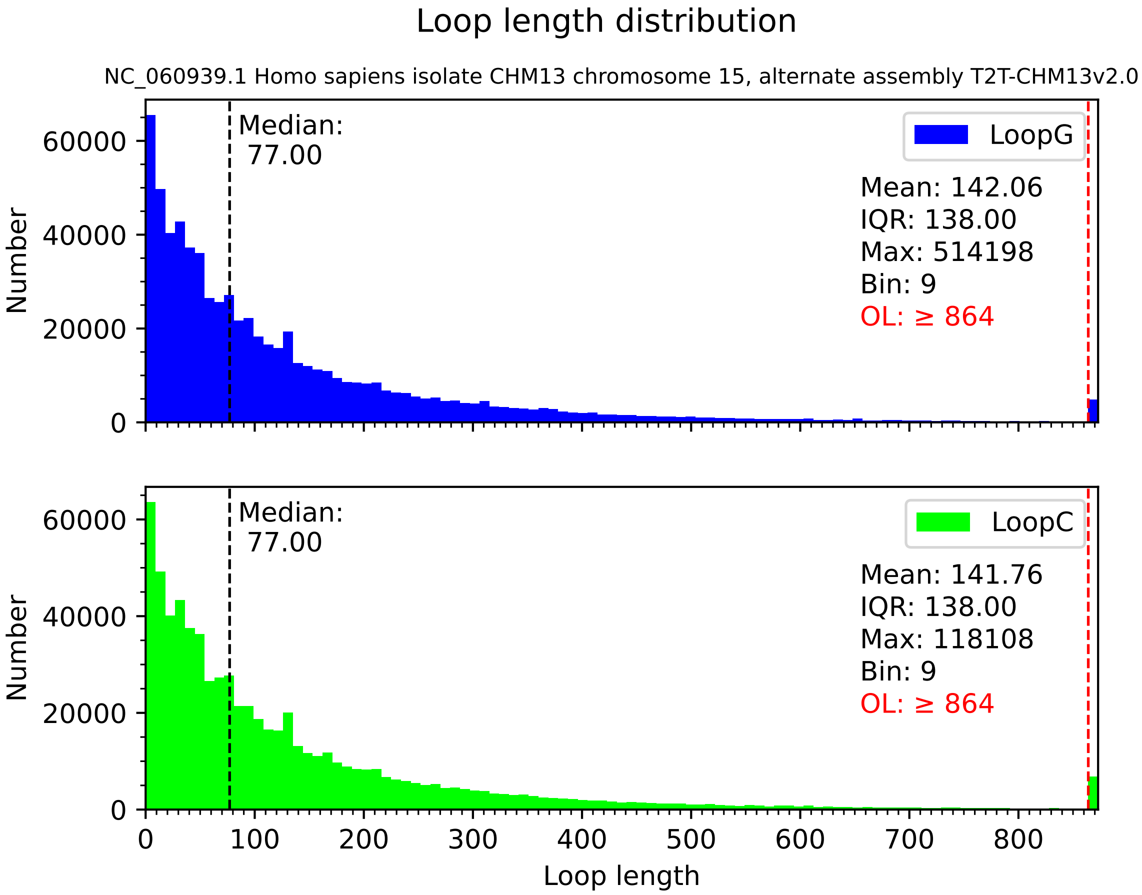

Loop length distribution
NC_060939.1 Homo sapiens isolate CHM13 chromosome 15, alternate assembly T2T-CHM13v2.0
Number
Median:
77.00
LoopG
Mean: 142.06
IQR: 138.00
Max: 514198
Bin: 9
OL: ≥ 864
0
20000
40000
60000
Number
Median:
77.00
LoopC
Mean: 141.76
IQR: 138.00
Max: 118108
Bin: 9
OL: ≥ 864
0
20000
40000
60000
0
100
200
300
400
500
600
700
800
Loop length

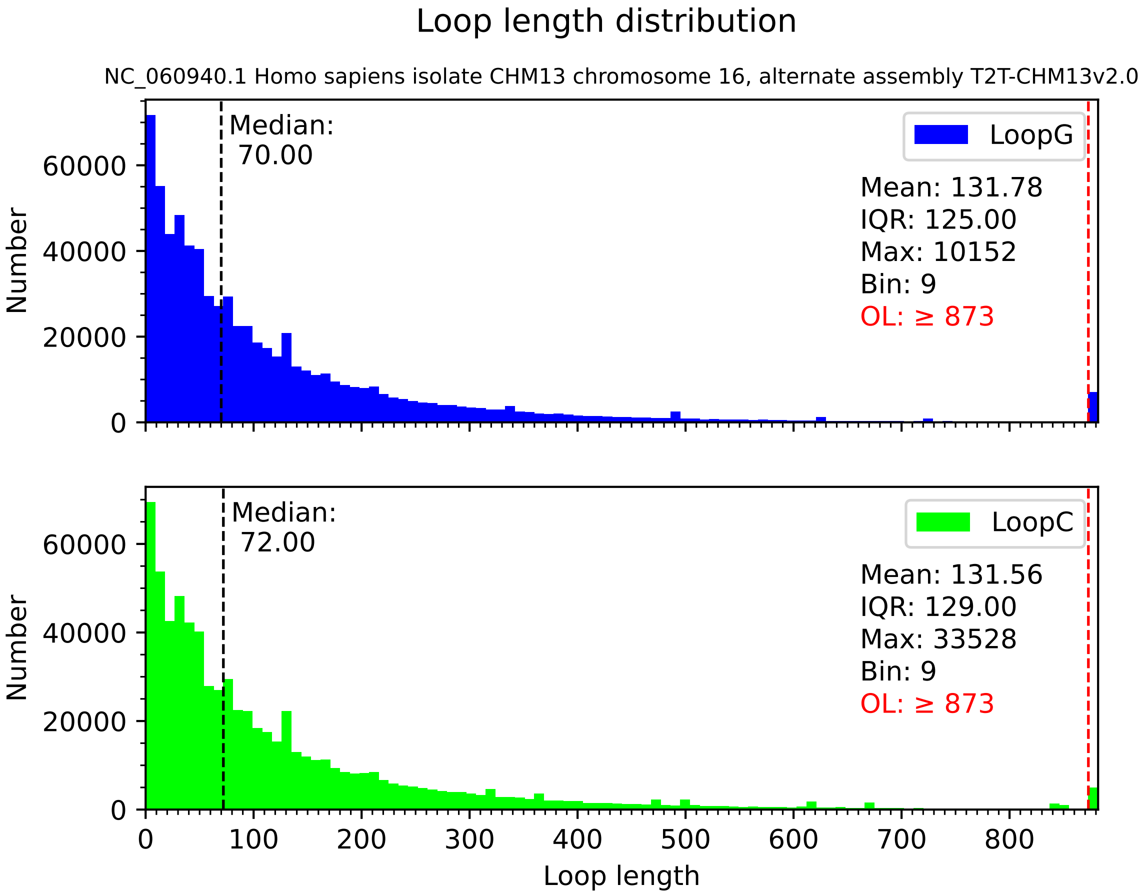

Loop length distribution
NC_060940.1 Homo sapiens isolate CHM13 chromosome 16, alternate assembly T2T-CHM13v2.0
Median:
70.00
LoopG
Mean: 131.78
IQR: 125.00
Max: 10152
Bin: 9
OL: ≥ 873
Number
0
20000
40000
60000
Median:
72.00
LoopC
Mean: 131.56
IQR: 129.00
Max: 33528
Bin: 9
OL: ≥ 873
Number
0
20000
40000
60000
0
100
200
300
400
500
600
700
800
Loop length

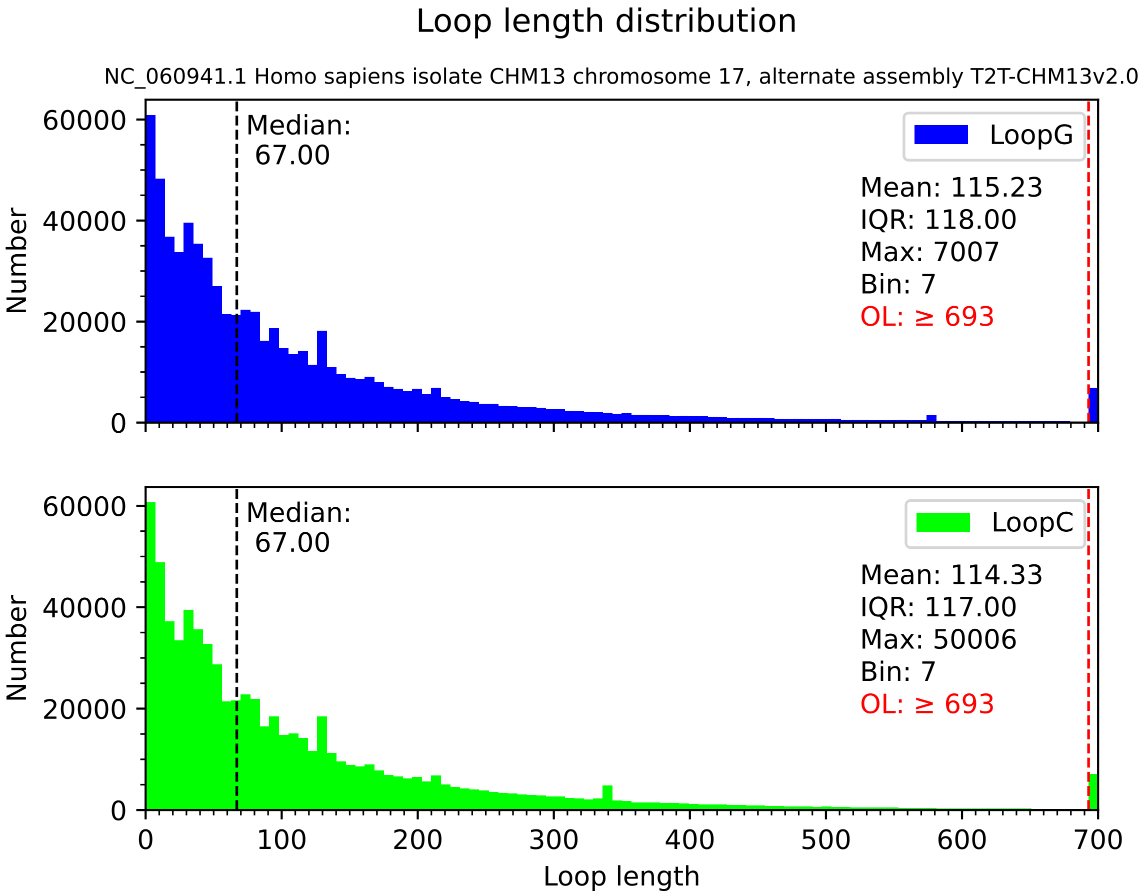

Loop length distribution
NC_060941.1 Homo sapiens isolate CHM13 chromosome 17, alternate assembly T2T-CHM13v2.0
Number
Median:
67.00
LoopG
Mean: 115.23
IQR: 118.00
Max: 7007
Bin: 7
OL: ≥ 693
Median:
67.00
LoopC
Mean: 114.33
IQR: 117.00
Max: 50006
Bin: 7
OL: ≥ 693
0
20000
40000
60000
0
100
200
300
400
500
600
700
Loop length

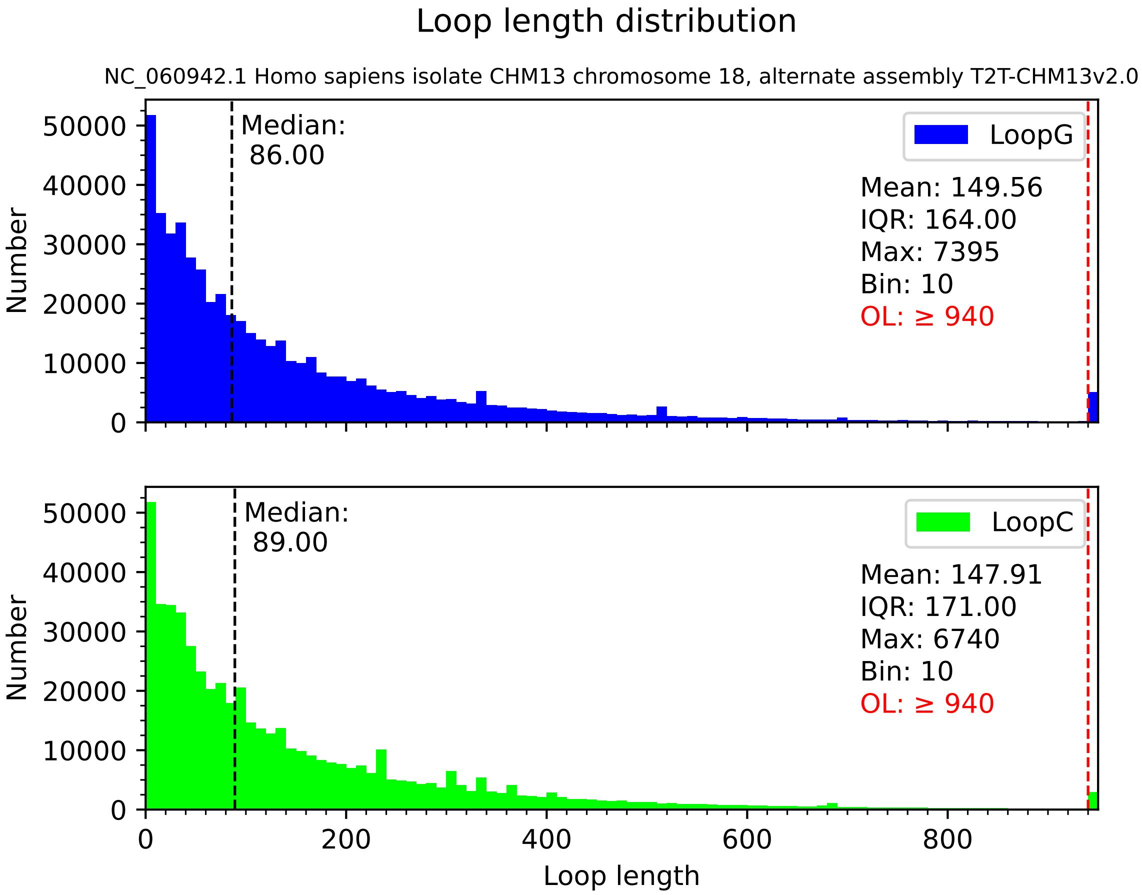

Loop length distribution
NC_060942.1 Homo sapiens isolate CHM13 chromosome 18, alternate assembly T2T-CHM13v2.0
Median:
86.00
LoopG
Mean: 149.56
IQR: 164.00
Max: 7395
Bin: 10
OL: ≥ 940
Median:
89.00
LoopC
Mean: 147.91
IQR: 171.00
Max: 6740
Bin: 10
OL: ≥ 940
Number
Loop length
0
10000
20000
30000
40000
50000
0
200
400
600
800

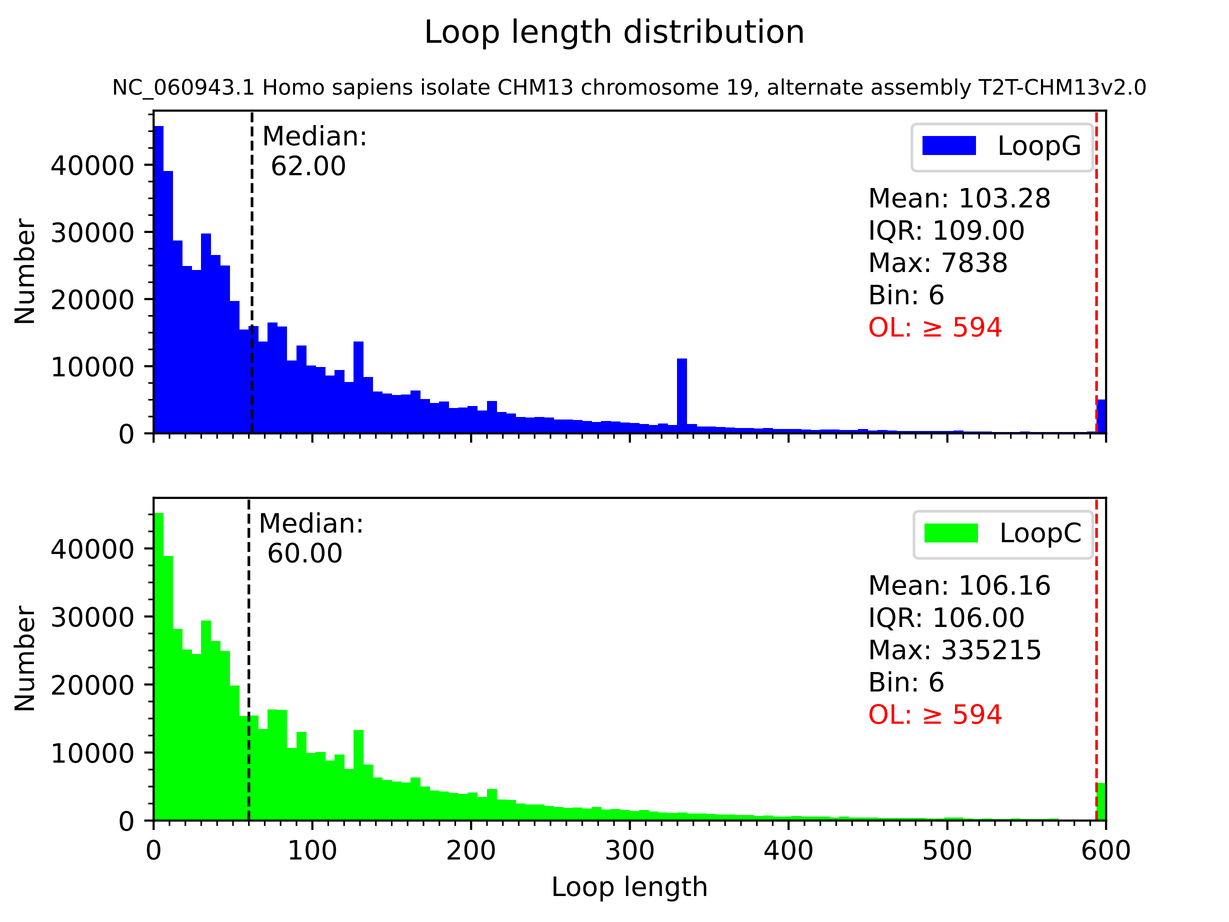

Loop length distribution
NC_060943.1 Homo sapiens isolate CHM13 chromosome 19, alternate assembly T2T-CHM13v2.0
Median:
62.00
LoopG
Mean: 103.28
IQR: 109.00
Max: 7838
Bin: 6
OL: ≥ 594
Median:
60.00
LoopC
Mean: 106.16
IQR: 106.00
Max: 335215
Bin: 6
OL: ≥ 594
Number
Loop length
0
10000
20000
30000
40000
100
200
300
400
500
600

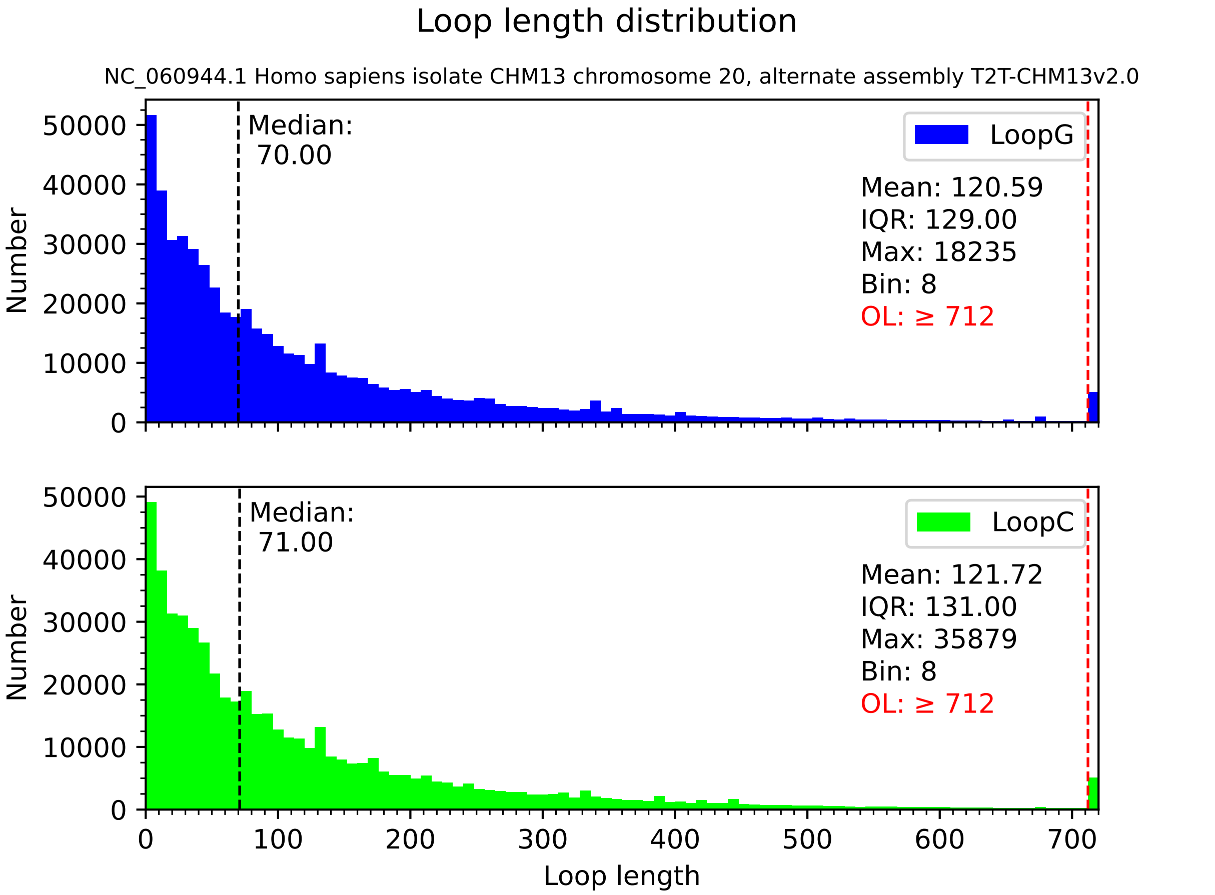

Loop length distribution
NC_060944.1 Homo sapiens isolate CHM13 chromosome 20, alternate assembly T2T-CHM13v2.0
Median:
70.00
LoopG
Mean: 120.59
IQR: 129.00
Max: 18235
Bin: 8
OL: ≥ 712
Median:
71.00
LoopC
Mean: 121.72
IQR: 131.00
Max: 35879
Bin: 8
OL: ≥ 712
Number
Loop length

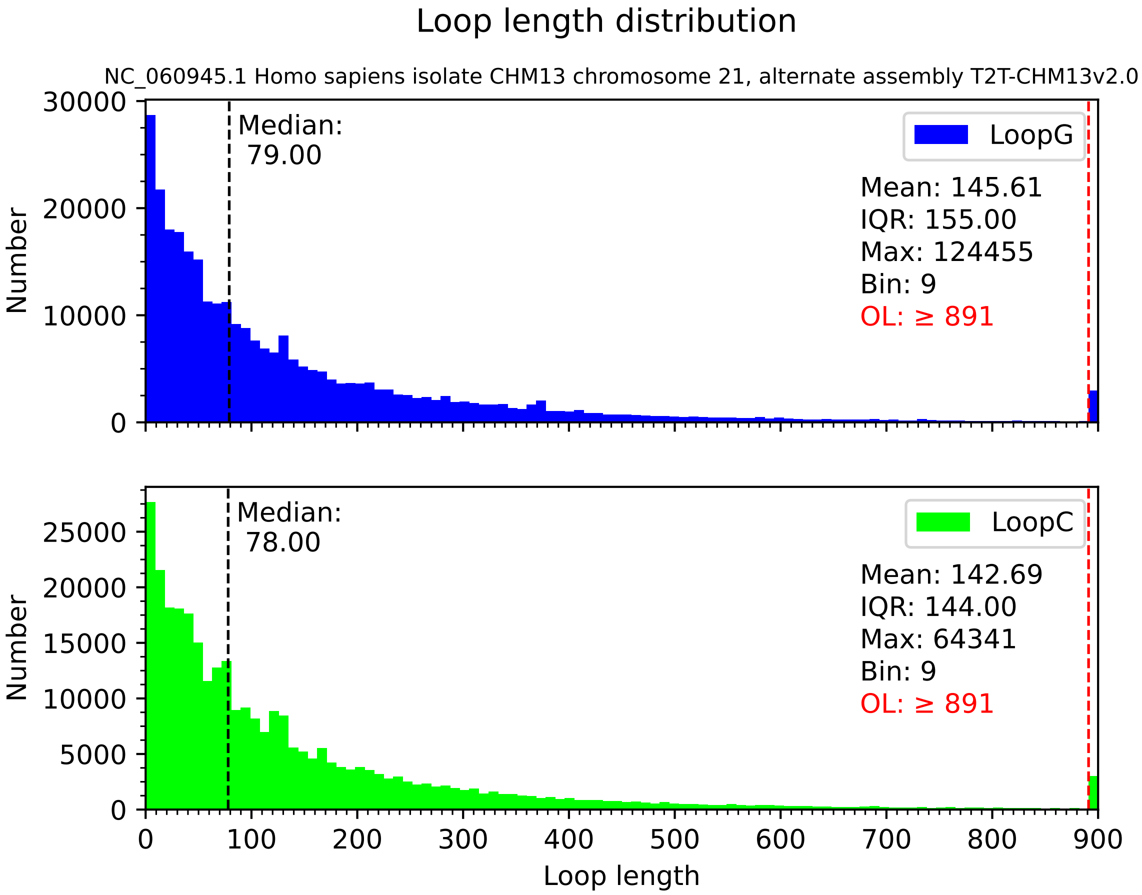

Loop length distribution
NC_060945.1 Homo sapiens isolate CHM13 chromosome 21, alternate assembly T2T-CHM13v2.0
LoopG
Median:
79.00
Mean: 145.61
IQR: 155.00
Max: 124455
Bin: 9
OL: ≥ 891
LoopC
Median:
78.00
Mean: 142.69
IQR: 144.00
Max: 64341
Bin: 9
OL: ≥ 891
Number
Loop length
0
10000
20000
30000
5000
15000
25000
100
200
300
400
500
600
700
800
900

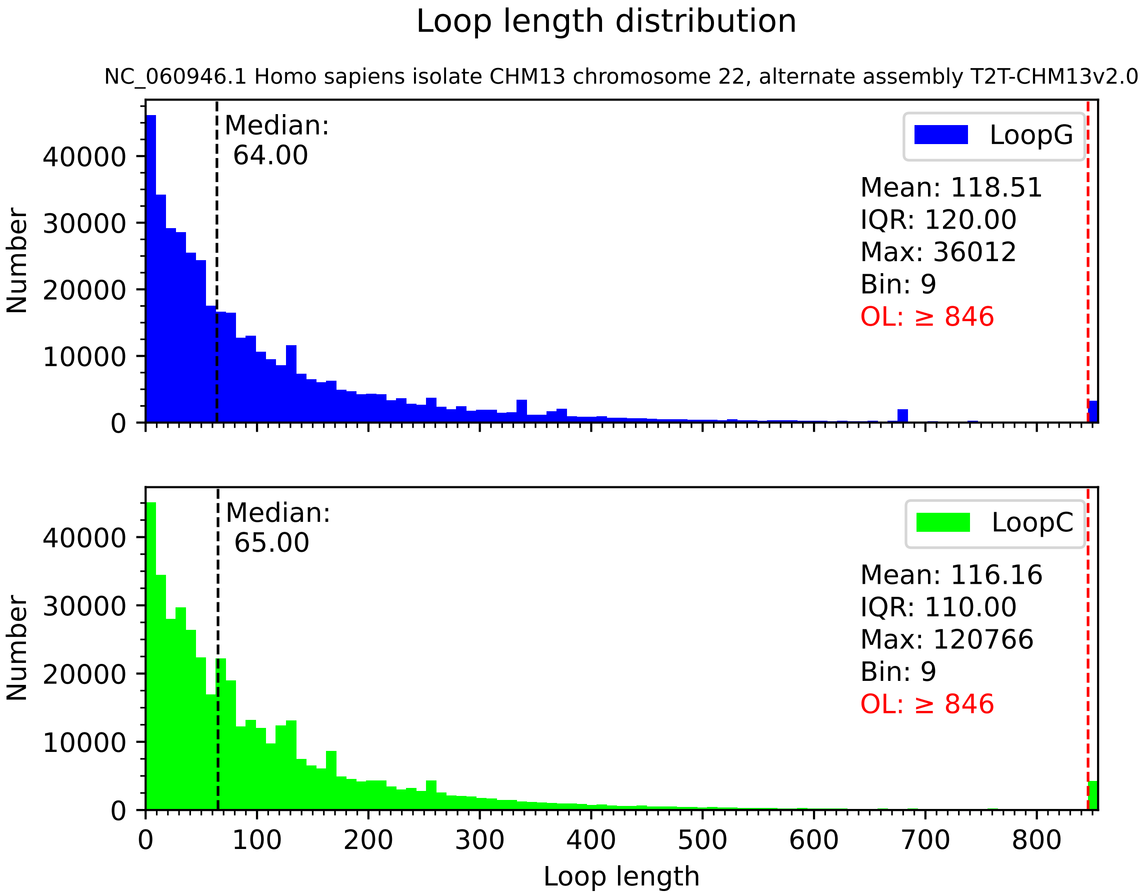

Loop length distribution
NC_060946.1 Homo sapiens isolate CHM13 chromosome 22, alternate assembly T2T-CHM13v2.0
Median:
64.00
LoopG
Mean: 118.51
IQR: 120.00
Max: 36012
Bin: 9
OL: ≥ 846
Median:
65.00
LoopC
Mean: 116.16
IQR: 110.00
Max: 120766
Bin: 9
OL: ≥ 846
Number
Number
0
10000
20000
30000
40000
0
100
200
300
400
500
600
700
800
Loop length

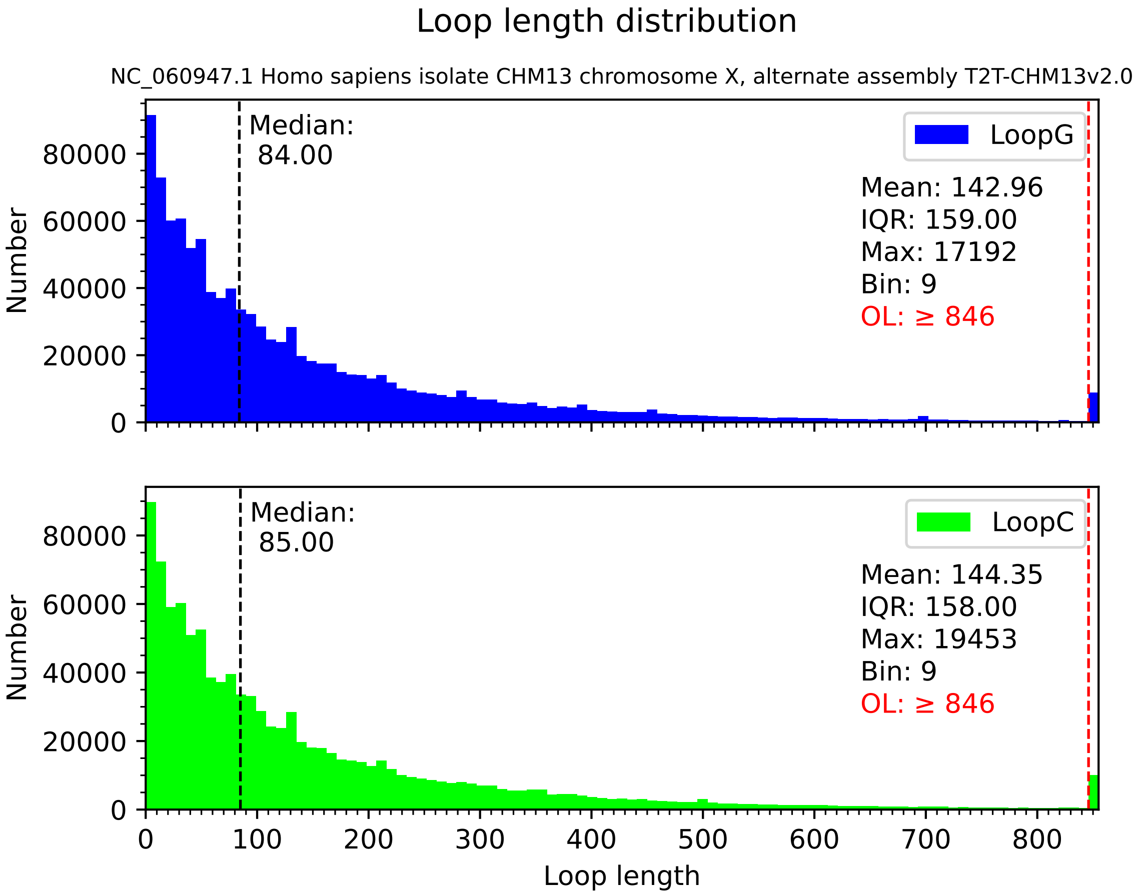

Loop length distribution
NC_060947.1 Homo sapiens isolate CHM13 chromosome X, alternate assembly T2T-CHM13v2.0
Median:
84.00
LoopG
Mean: 142.96
IQR: 159.00
Max: 17192
Bin: 9
OL: ≥ 846
Number
0
20000
40000
60000
80000
Median:
85.00
LoopC
Mean: 144.35
IQR: 158.00
Max: 19453
Bin: 9
OL: ≥ 846
Number
0
20000
40000
60000
80000
0
100
200
300
400
500
600
700
800
Loop length

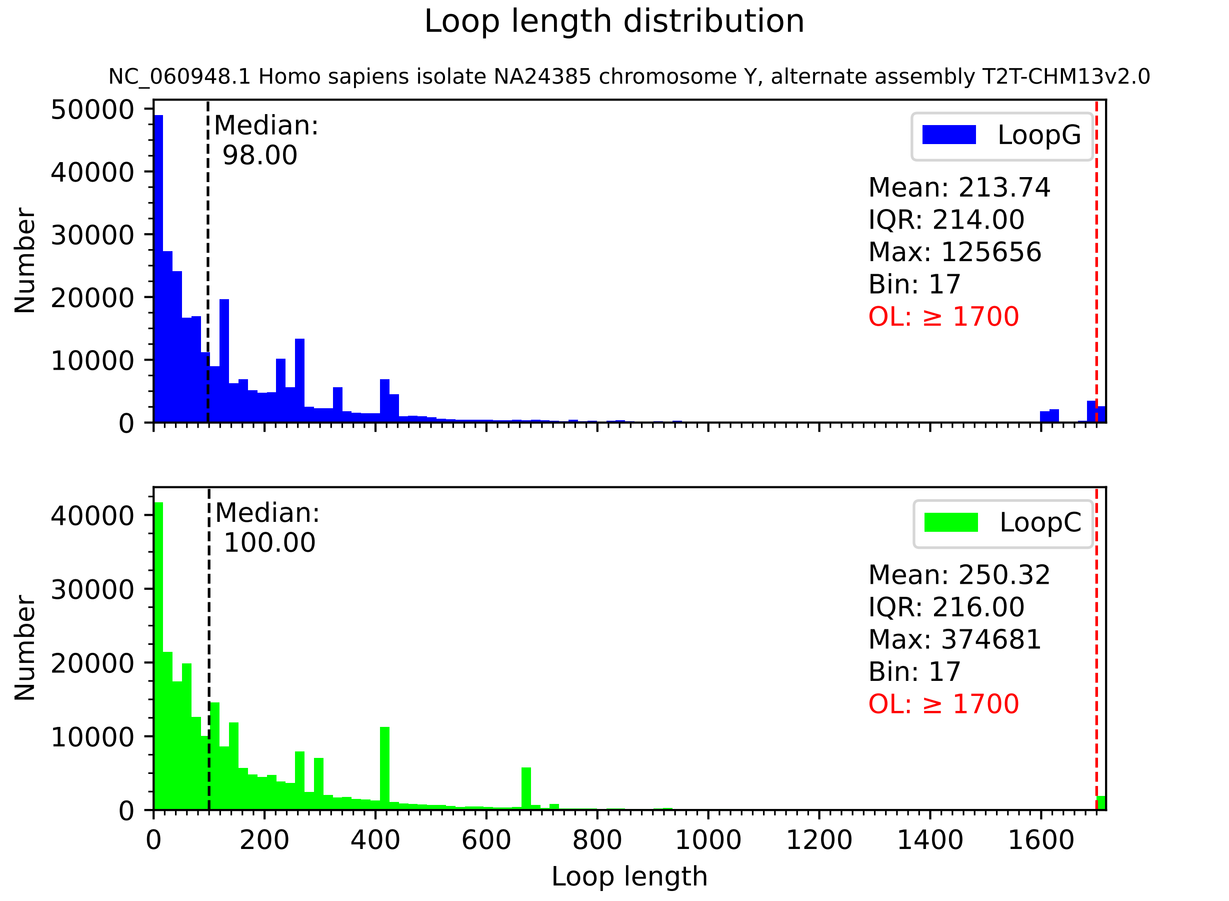
Loop length distribution
NC_060948.1 Homo sapiens isolate NA24385 chromosome Y, alternate assembly T2T-CHM13v2.0
Median:
98.00
LoopG
Mean: 213.74
IQR: 214.00
Max: 125656
Bin: 17
OL: ≥ 1700
Number
Median:
100.00
LoopC
Mean: 250.32
IQR: 216.00
Max: 374681
Bin: 17
OL: ≥ 1700
Number
Loop length
