## Supplementary file 6 for "FuzzyClusTeR: a web server for analysis of tandem and diffuse DNA repeat clusters with application to telomeric-like repeats"

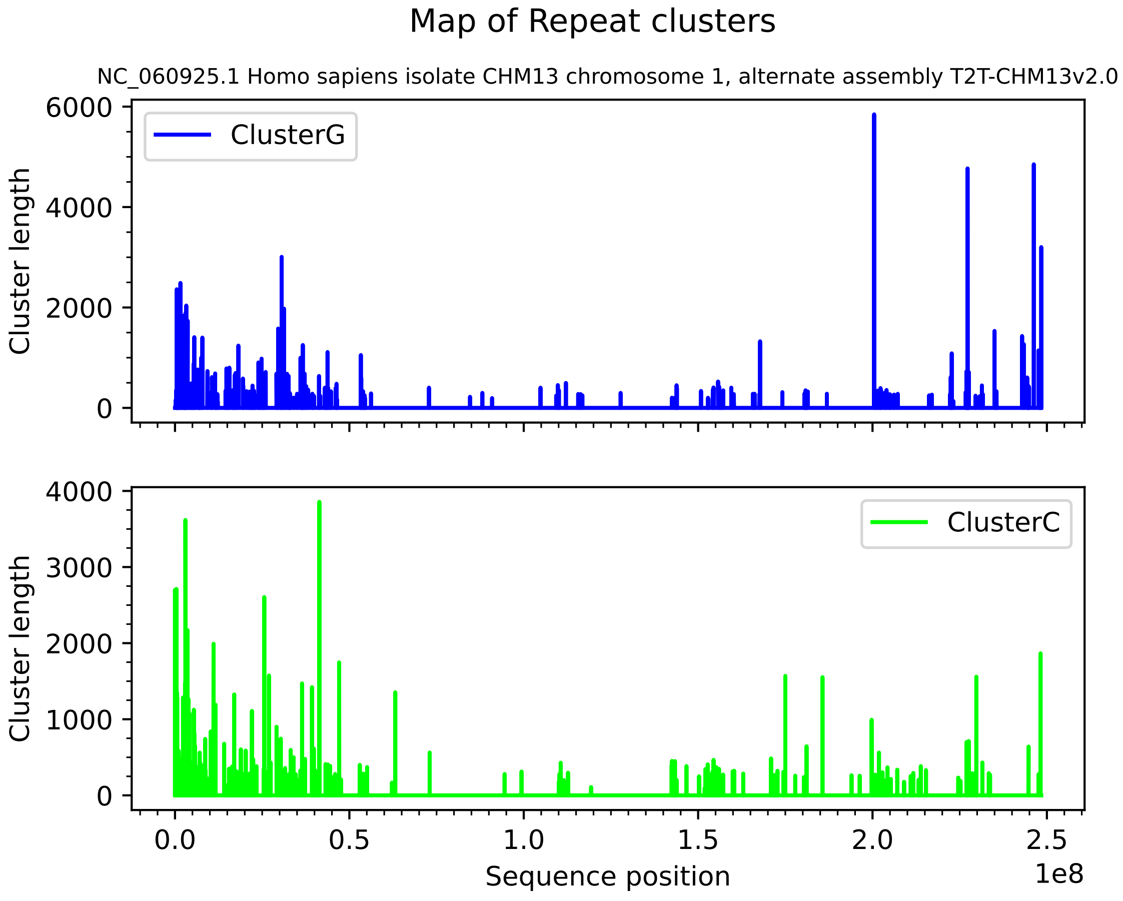
Map of Repeat clusters
NC_060925.1 Homo sapiens isolate CHM13 chromosome 1, alternate assembly T2T-CHM13v2.0
ClusterG
ClusterC
Cluster length
Cluster length
6000
4000
2000
0
4000
3000
2000
1000
0
0.0
0.5
1.0
1.5
2.0
2.5
1e8
Sequence position

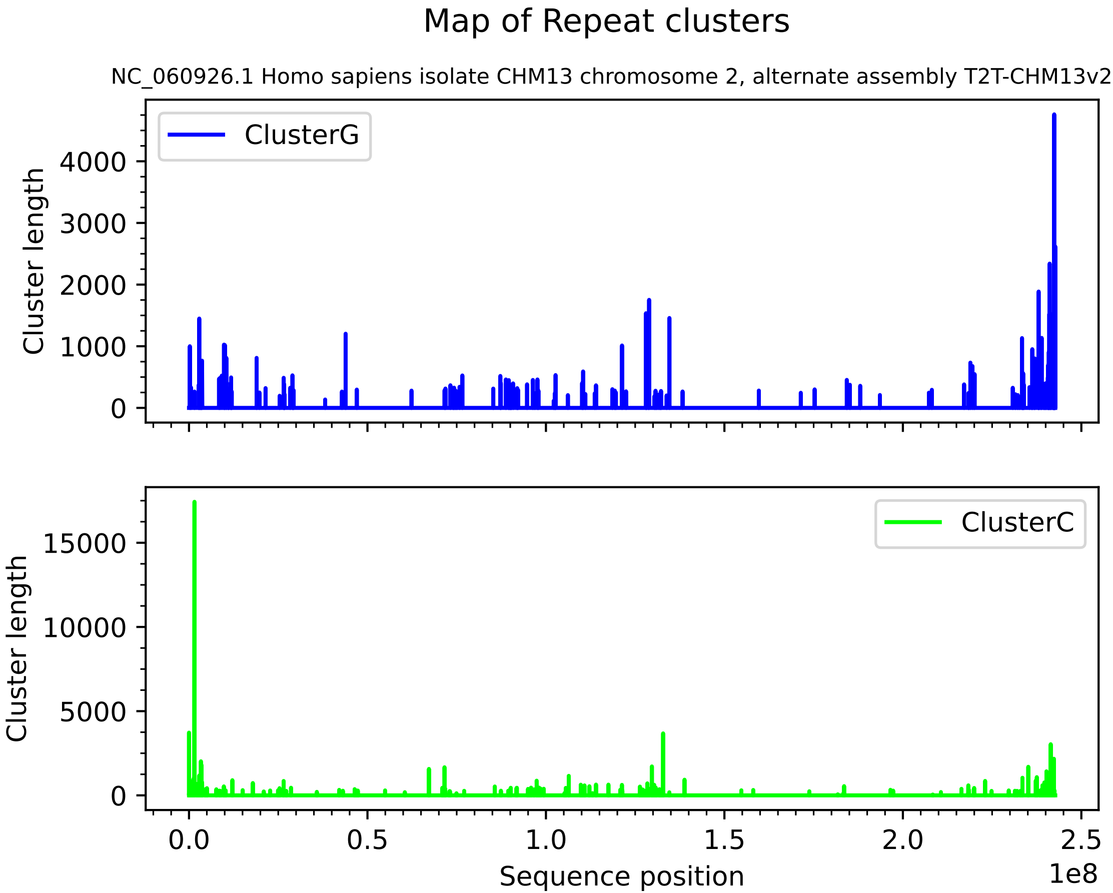
Map of Repeat clusters
NC_060926.1 Homo sapiens isolate CHM13 chromosome 2, alternate assembly T2T-CHM13v2.0
ClusterG
Cluster length
0
1000
2000
3000
4000
ClusterC
Cluster length
0
5000
10000
15000
0.0
0.5
1.0
1.5
2.0
2.5
1e8
Sequence position

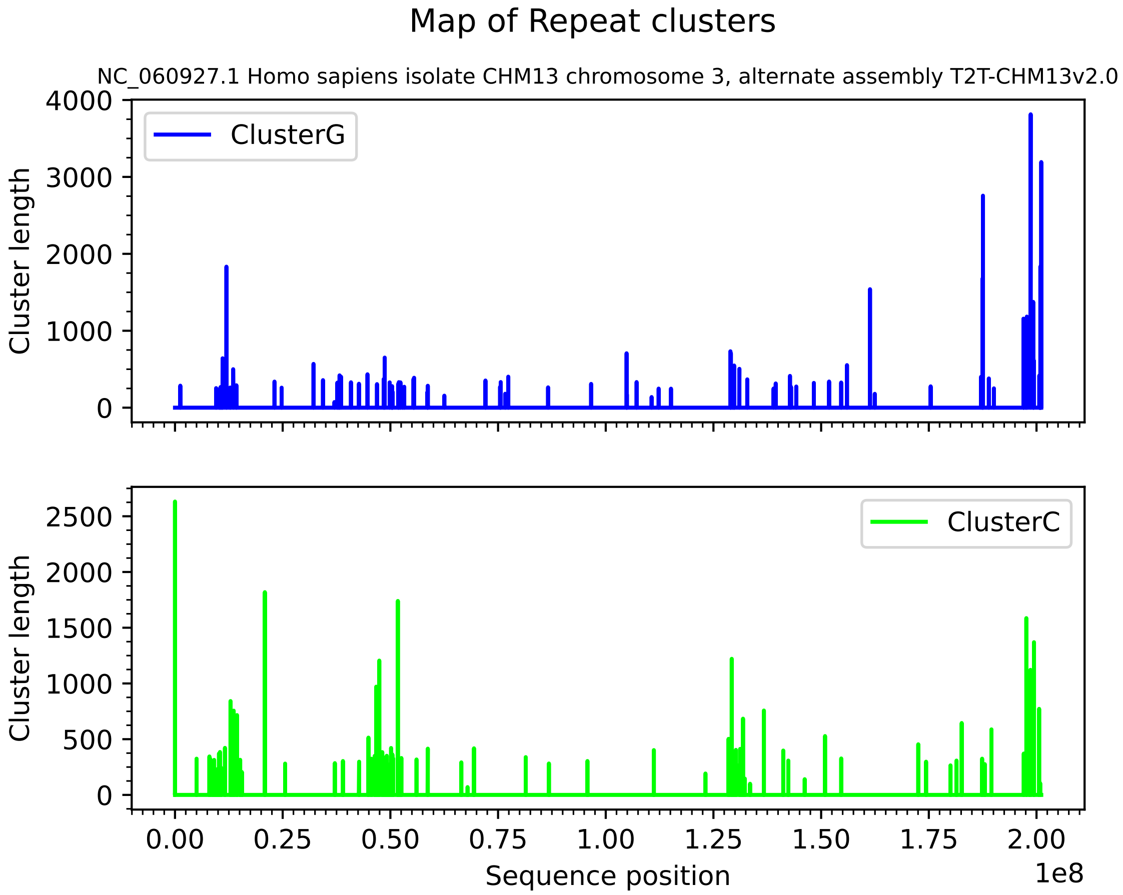

Map of Repeat clusters
NC_060927.1 Homo sapiens isolate CHM13 chromosome 3, alternate assembly T2T-CHM13v2.0
ClusterG
Cluster length
0
1000
2000
3000
4000
ClusterC
Cluster length
0
500
1000
1500
2000
2500
0.00
0.25
0.50
0.75
1.00
1.25
1.50
1.75
2.00
1e8
Sequence position

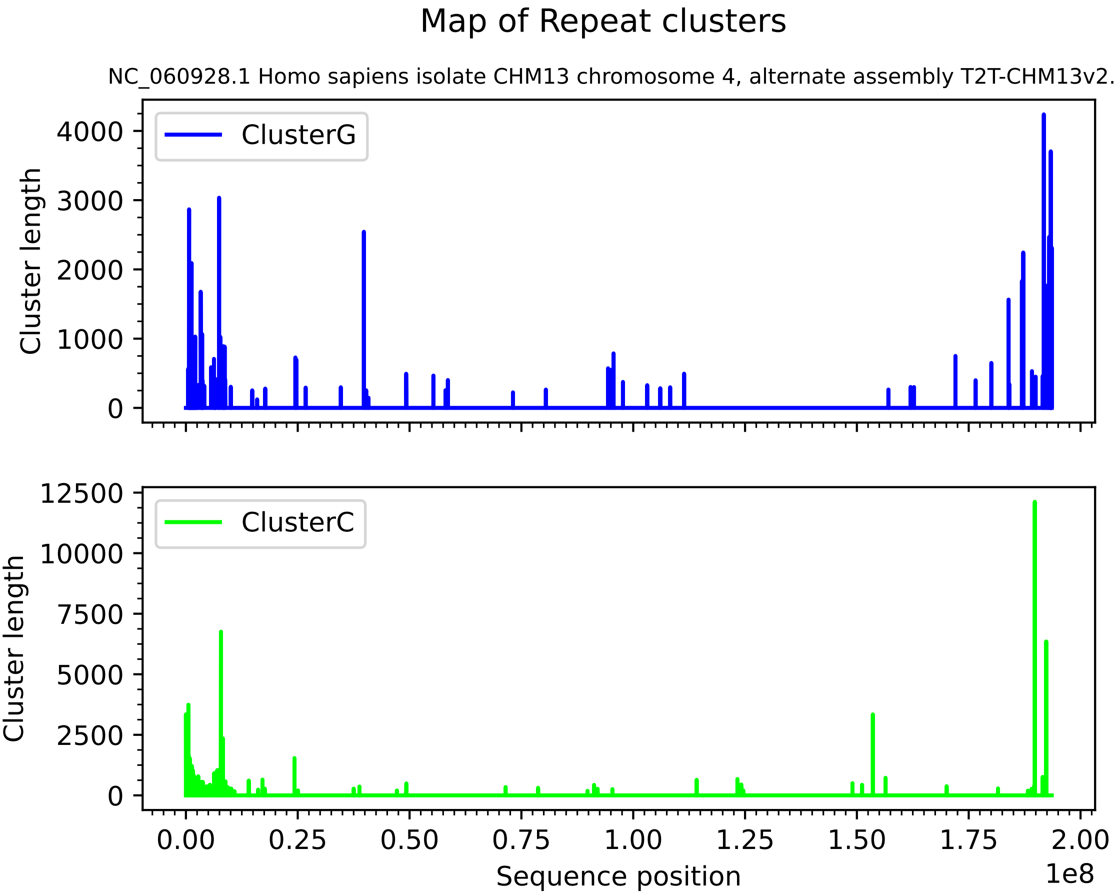
Map of Repeat clusters
NC_060928.1 Homo sapiens isolate CHM13 chromosome 4, alternate assembly T2T-CHM13v2.0
ClusterG
Cluster length
4000
3000
2000
1000
0
ClusterC
Cluster length
12500
10000
7500
5000
2500
0
0.00
0.25
0.50
0.75
1.00
1.25
1.50
1.75
2.00
1e8
Sequence position

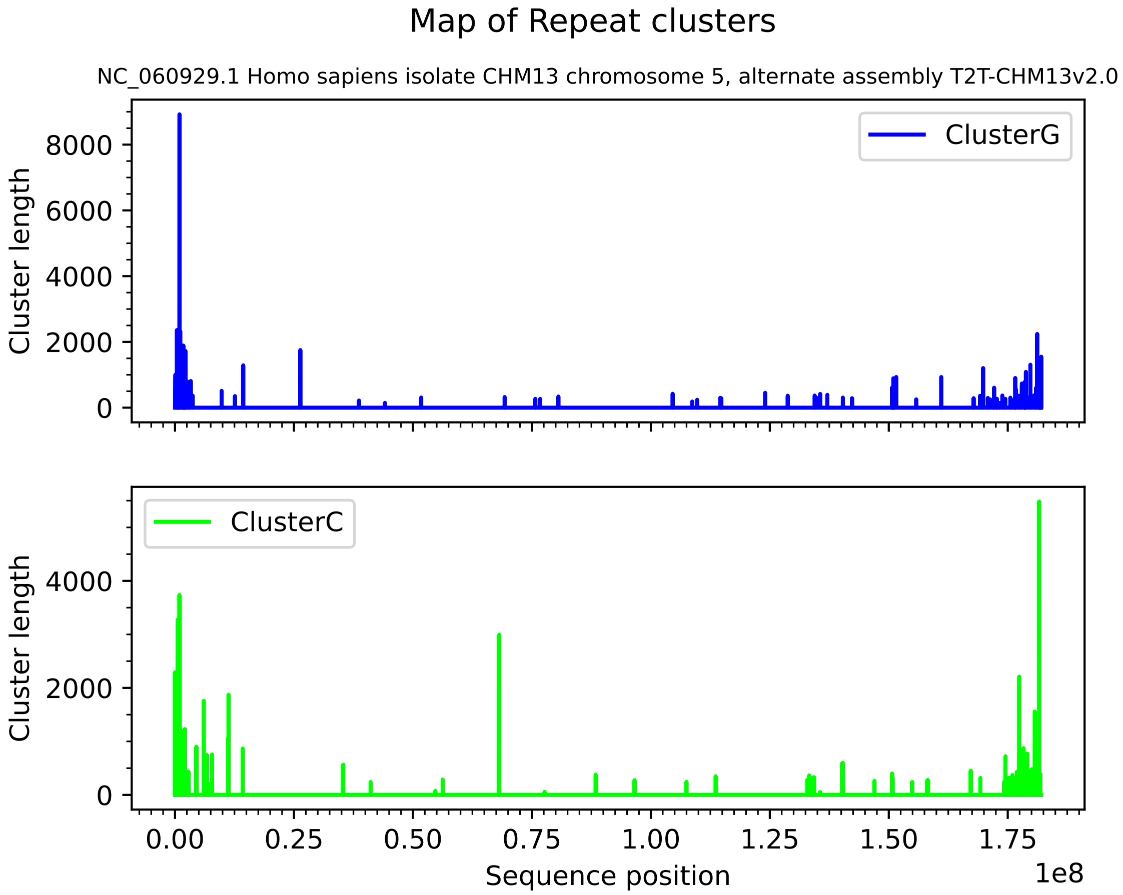

Map of Repeat clusters
NC_060929.1 Homo sapiens isolate CHM13 chromosome 5, alternate assembly T2T-CHM13v2.0
ClusterG
Cluster length
0
2000
4000
6000
8000
ClusterC
Cluster length
0
2000
4000
0.00
0.25
0.50
0.75
1.00
1.25
1.50
1.75
1e8
Sequence position

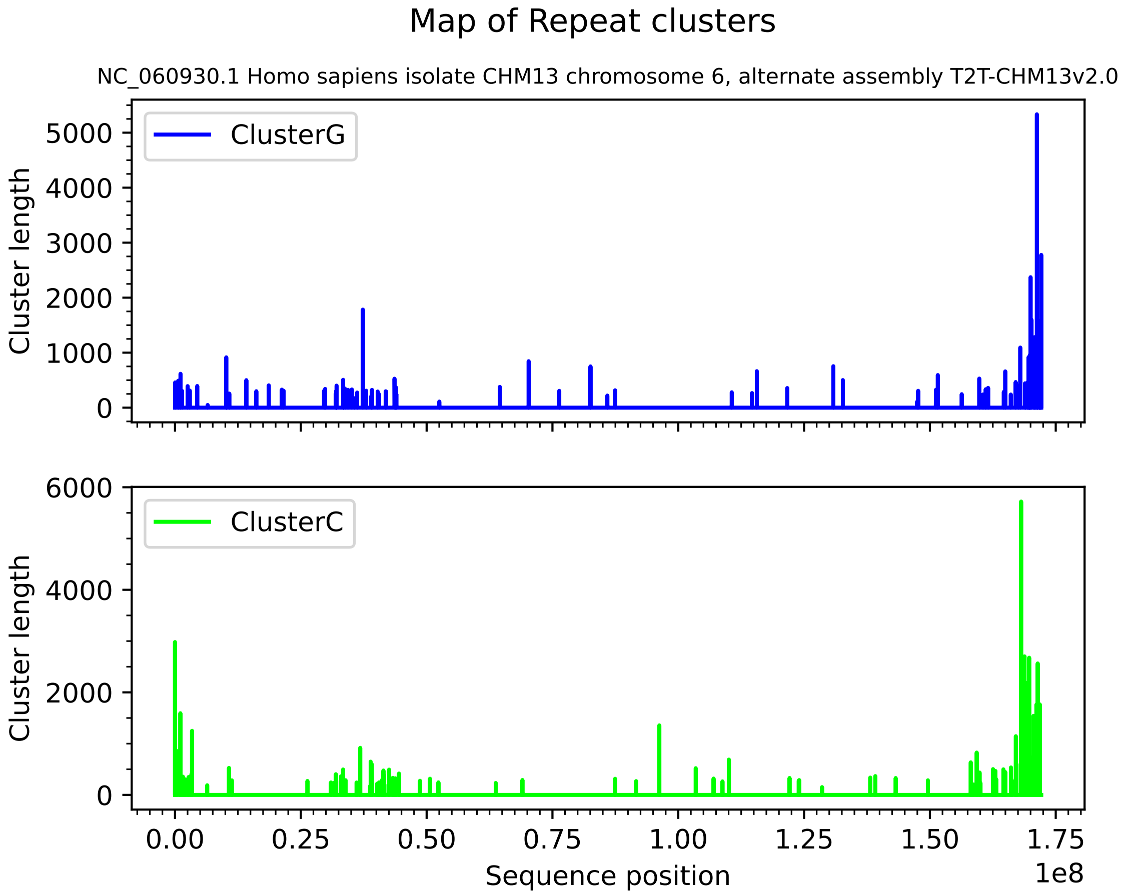
Map of Repeat clusters
NC_060930.1 Homo sapiens isolate CHM13 chromosome 6, alternate assembly T2T-CHM13v2.0
ClusterG
Cluster length
5000
4000
3000
2000
1000
0
ClusterC
Cluster length
6000
4000
2000
0
0.00
0.25
0.50
0.75
1.00
1.25
1.50
1.75
1e8
Sequence position

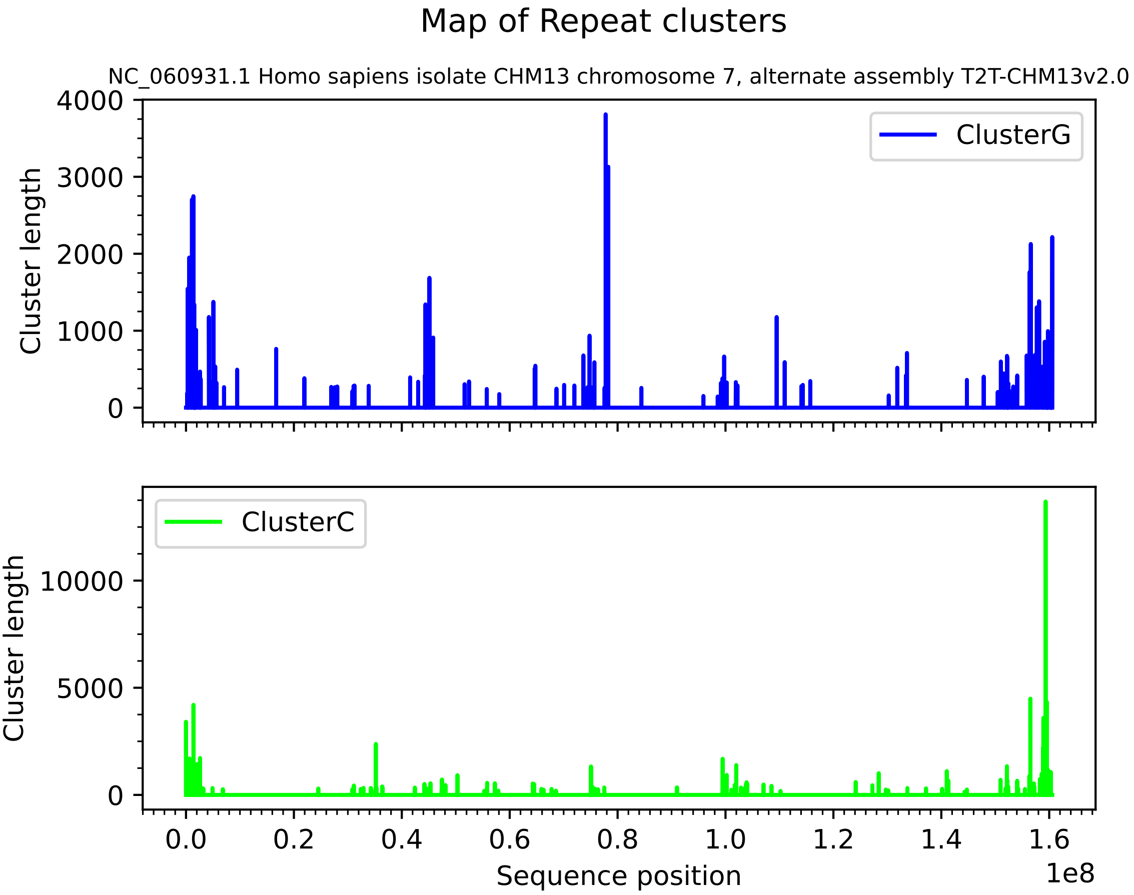

Map of Repeat clusters
NC_060931.1 Homo sapiens isolate CHM13 chromosome 7, alternate assembly T2T-CHM13v2.0
ClusterG
Cluster length
4000
3000
2000
1000
0
ClusterC
Cluster length
10000
5000
0
0.0
0.2
0.4
0.6
0.8
1.0
1.2
1.4
1.6
1e8
Sequence position

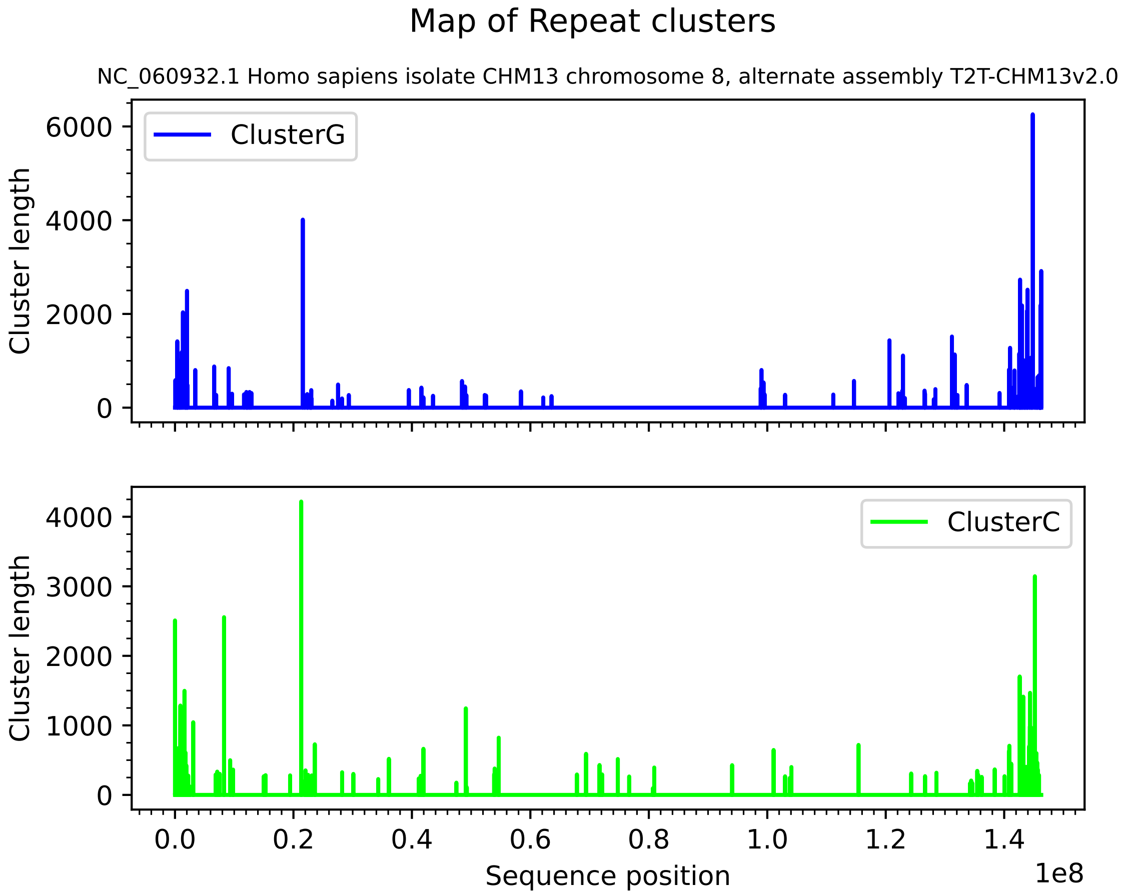

Map of Repeat clusters
NC_060932.1 Homo sapiens isolate CHM13 chromosome 8, alternate assembly T2T-CHM13v2.0
ClusterG
ClusterC
Cluster length
Cluster length
6000
4000
2000
0
4000
3000
2000
1000
0
0.0
0.2
0.4
0.6
0.8
1.0
1.2
1.4
1e8
Sequence position

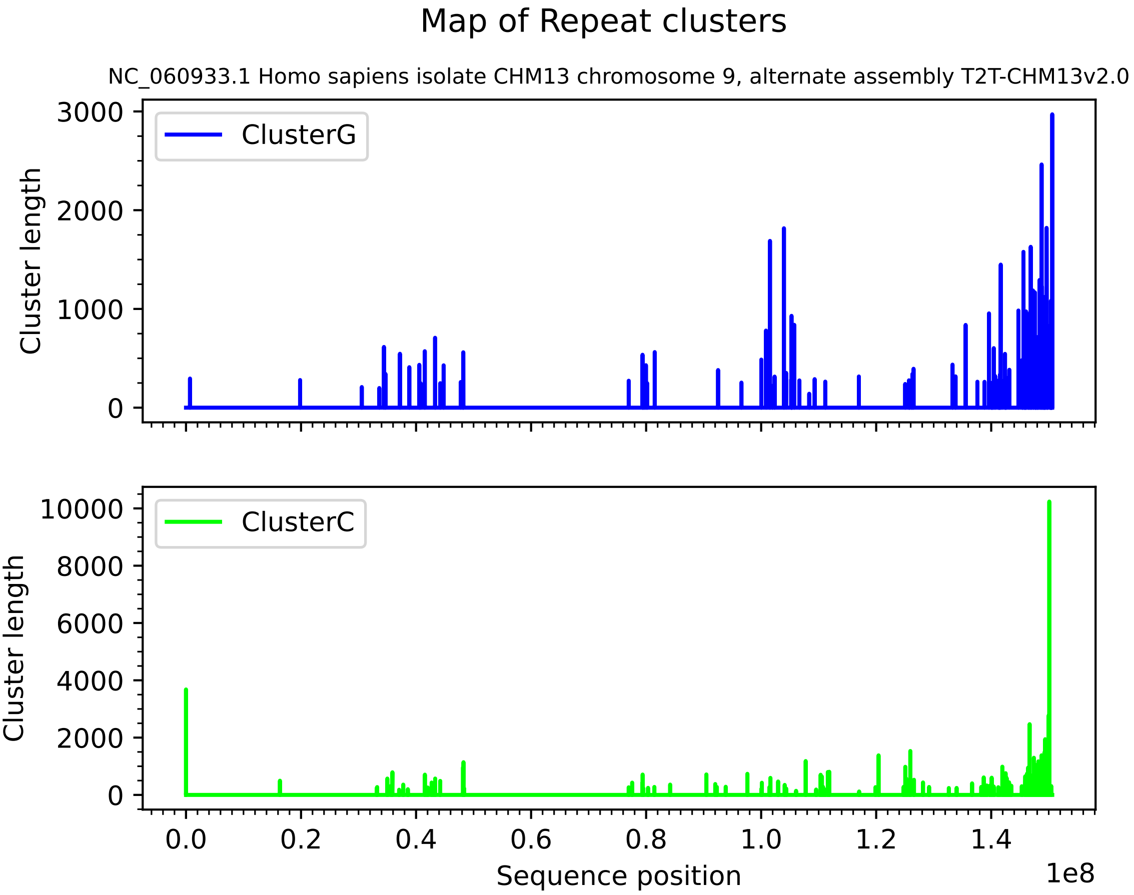

Map of Repeat clusters
NC_060933.1 Homo sapiens isolate CHM13 chromosome 9, alternate assembly T2T-CHM13v2.0
ClusterG
ClusterC
Cluster length
Cluster length
3000
2000
1000
0
10000
8000
6000
4000
2000
0
0.0
0.2
0.4
0.6
0.8
1.0
1.2
1.4
1e8
Sequence position

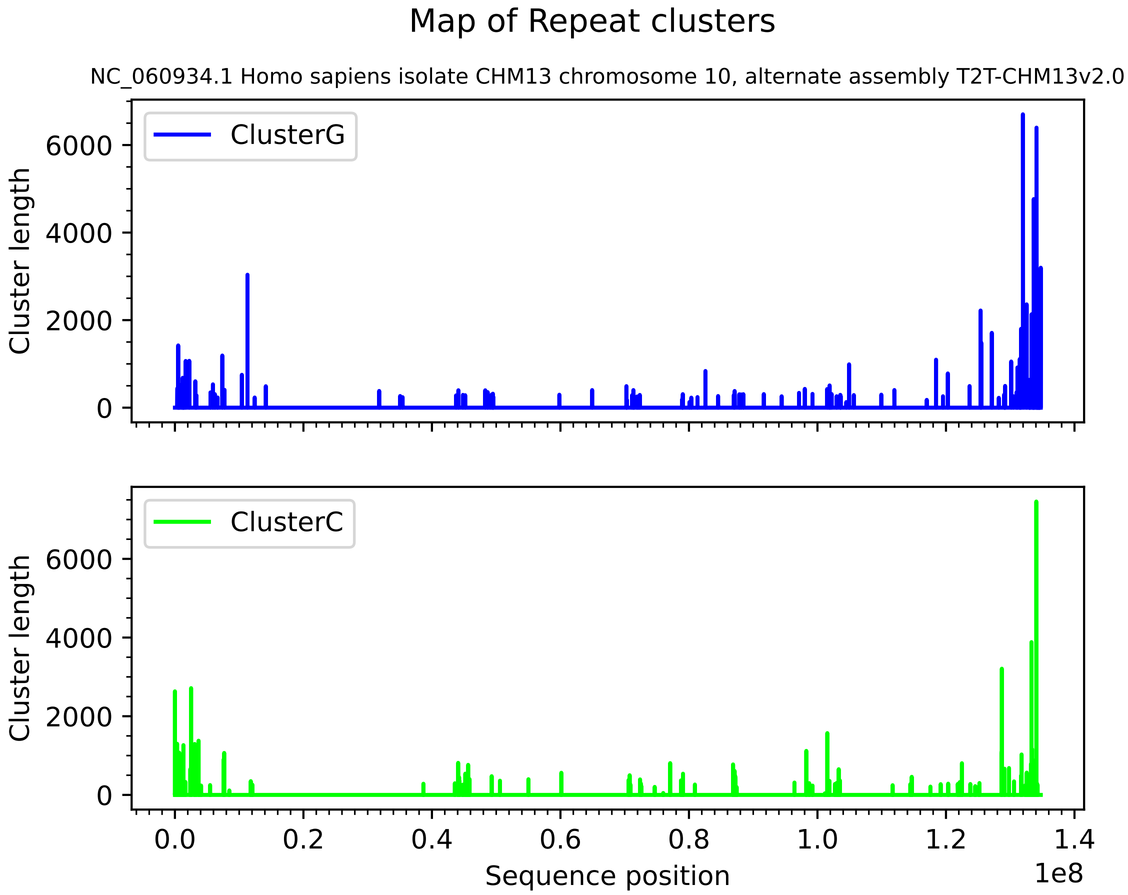

Map of Repeat clusters
NC_060934.1 Homo sapiens isolate CHM13 chromosome 10, alternate assembly T2T-CHM13v2.0
ClusterG
Cluster length
6000
4000
2000
0
ClusterC
Cluster length
6000
4000
2000
0
0.0
0.2
0.4
0.6
0.8
1.0
1.2
1.4
1e8
Sequence position

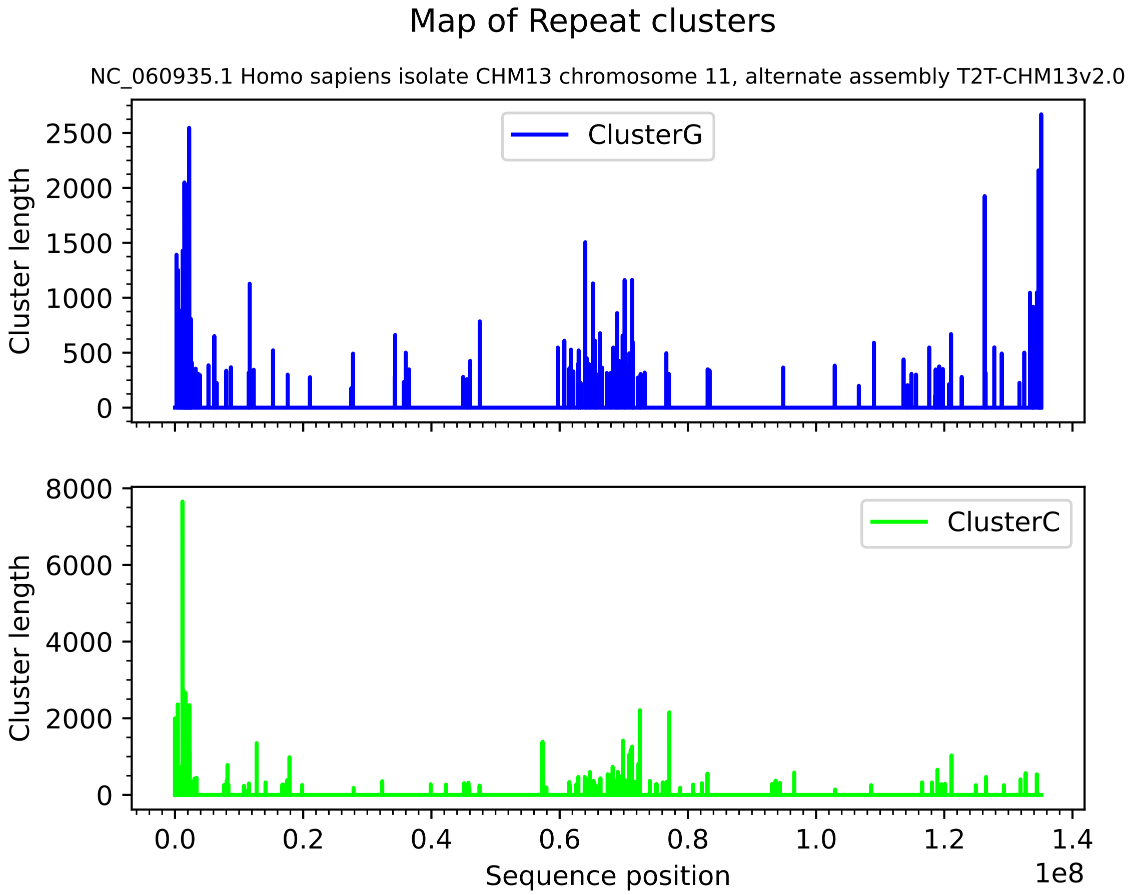
Map of Repeat clusters
NC_060935.1 Homo sapiens isolate CHM13 chromosome 11, alternate assembly T2T-CHM13v2.0
ClusterG
ClusterC
Cluster length
Cluster length
0
500
1000
1500
2000
2500
0
2000
4000
6000
8000
0.0
0.2
0.4
0.6
0.8
1.0
1.2
1.4
1e8
Sequence position

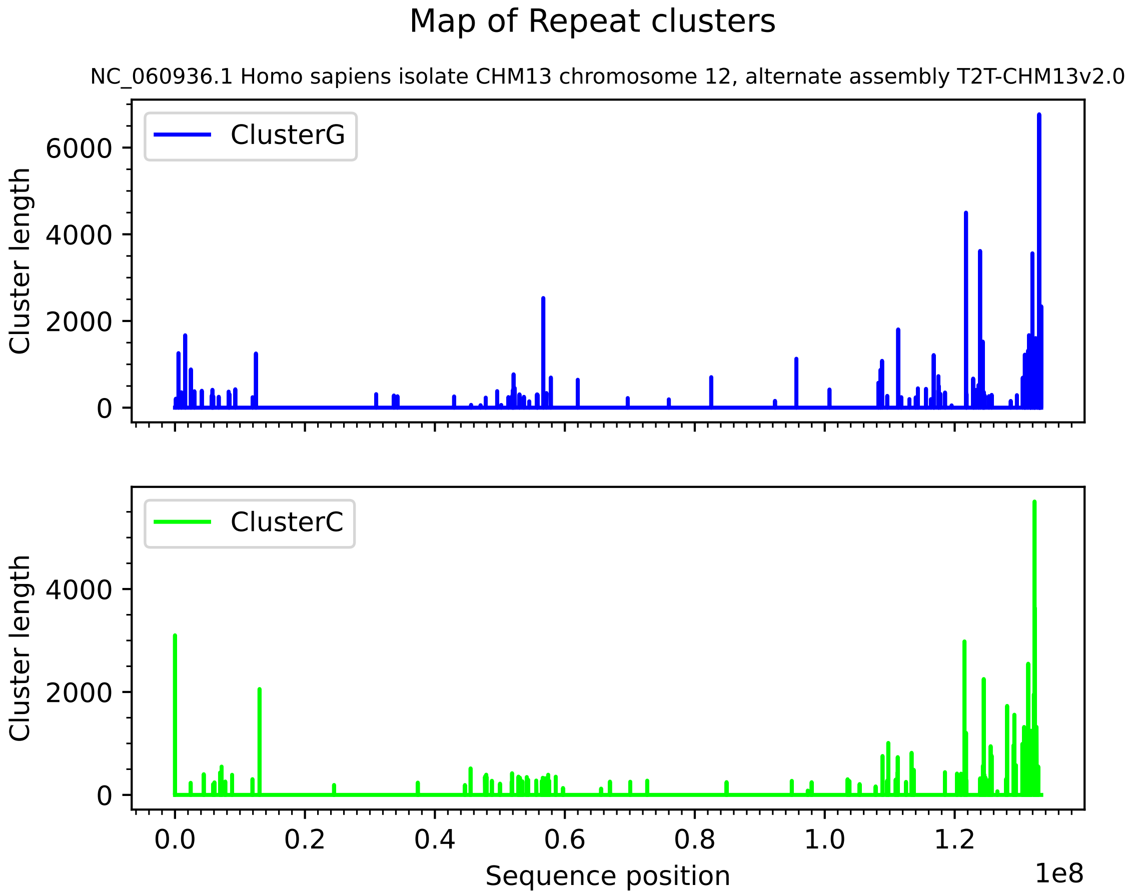
Map of Repeat clusters
NC_060936.1 Homo sapiens isolate CHM13 chromosome 12, alternate assembly T2T-CHM13v2.0
ClusterG
ClusterC
Cluster length
Cluster length
6000
4000
2000
0
4000
2000
0
0.0
0.2
0.4
0.6
0.8
1.0
1.2
1e8
Sequence position

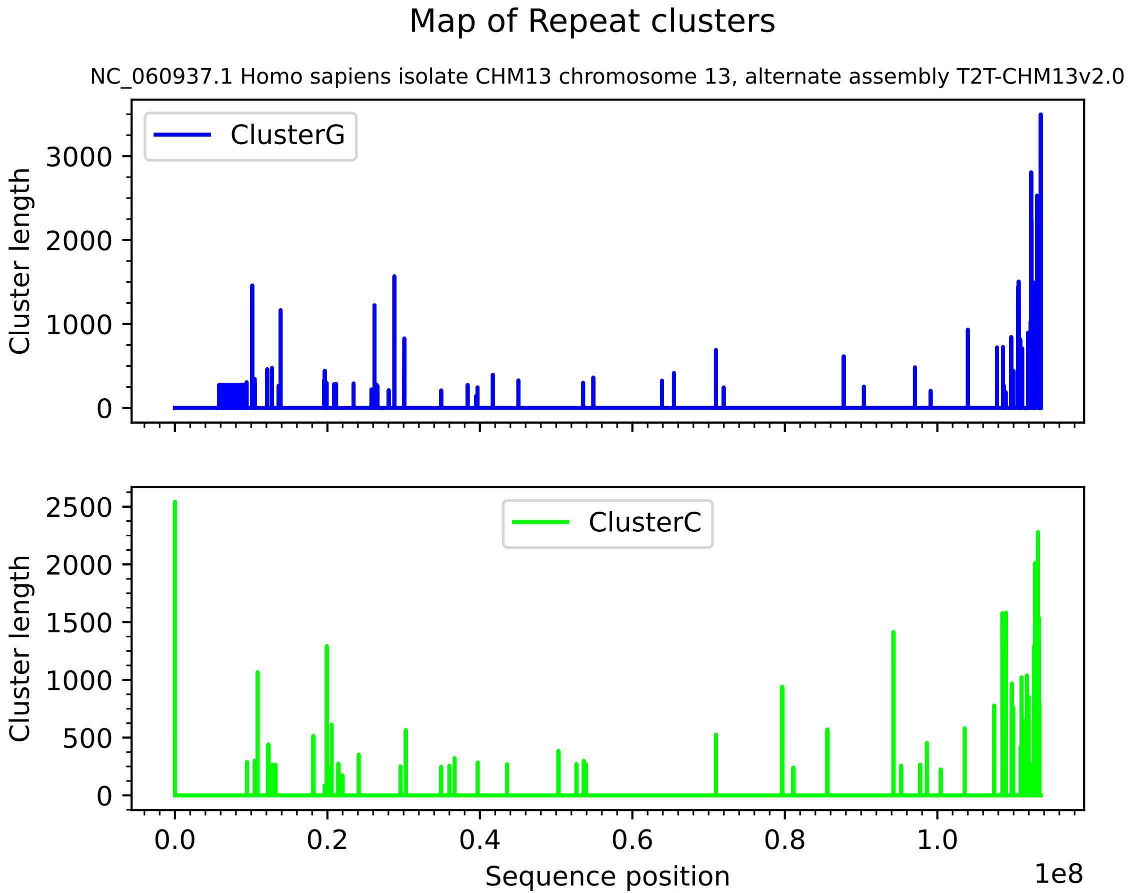

Map of Repeat clusters
NC_060937.1 Homo sapiens isolate CHM13 chromosome 13, alternate assembly T2T-CHM13v2.0
ClusterG
Cluster length
3000
2000
1000
0
ClusterC
Cluster length
2500
2000
1500
1000
500
0
0.0
0.2
0.4
0.6
0.8
1.0
1e8
Sequence position

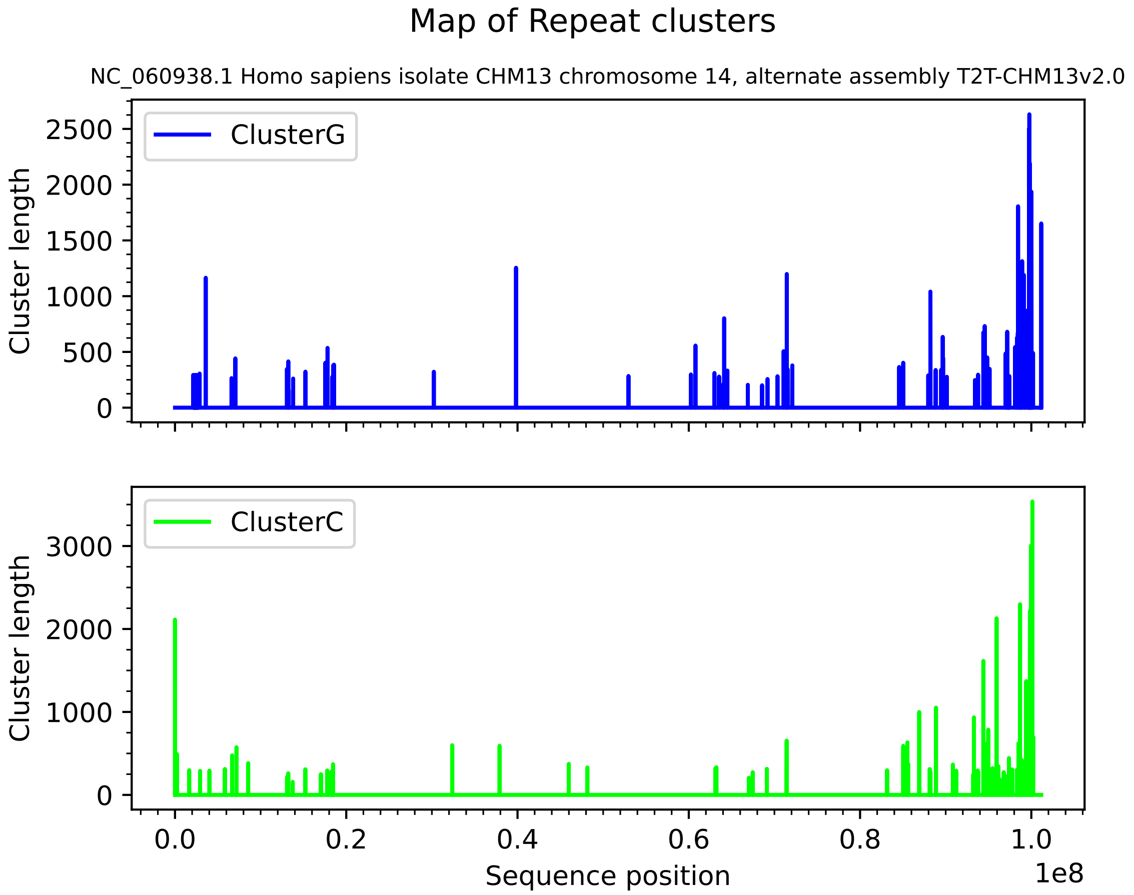
Map of Repeat clusters
NC_060938.1 Homo sapiens isolate CHM13 chromosome 14, alternate assembly T2T-CHM13v2.0
ClusterG
Cluster length
2500
2000
1500
1000
500
0
ClusterC
Cluster length
3000
2000
1000
0
0.0
0.2
0.4
0.6
0.8
1.0
1e8
Sequence position

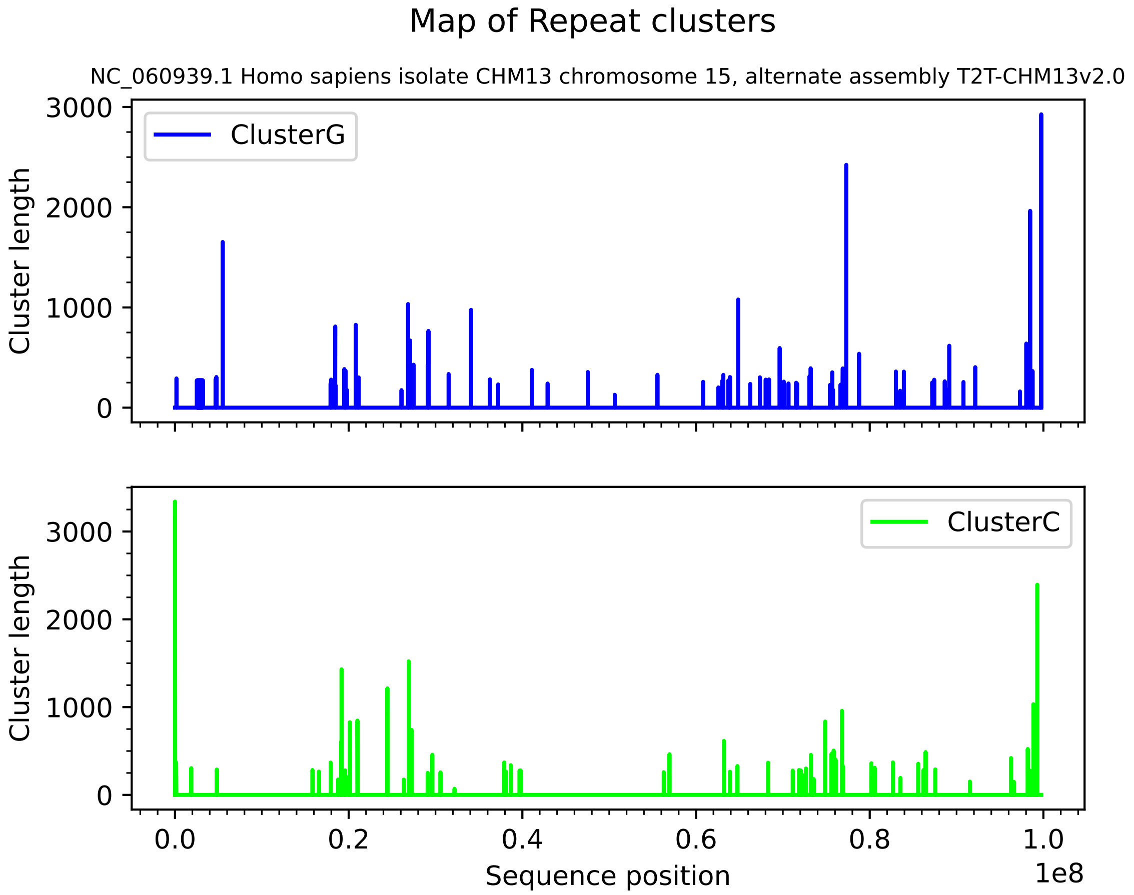

Map of Repeat clusters
NC_060939.1 Homo sapiens isolate CHM13 chromosome 15, alternate assembly T2T-CHM13v2.0
ClusterG
ClusterC
Cluster length
Cluster length
3000
2000
1000
0
3000
2000
1000
0
0.0
0.2
0.4
0.6
0.8
1.0
1e8
Sequence position

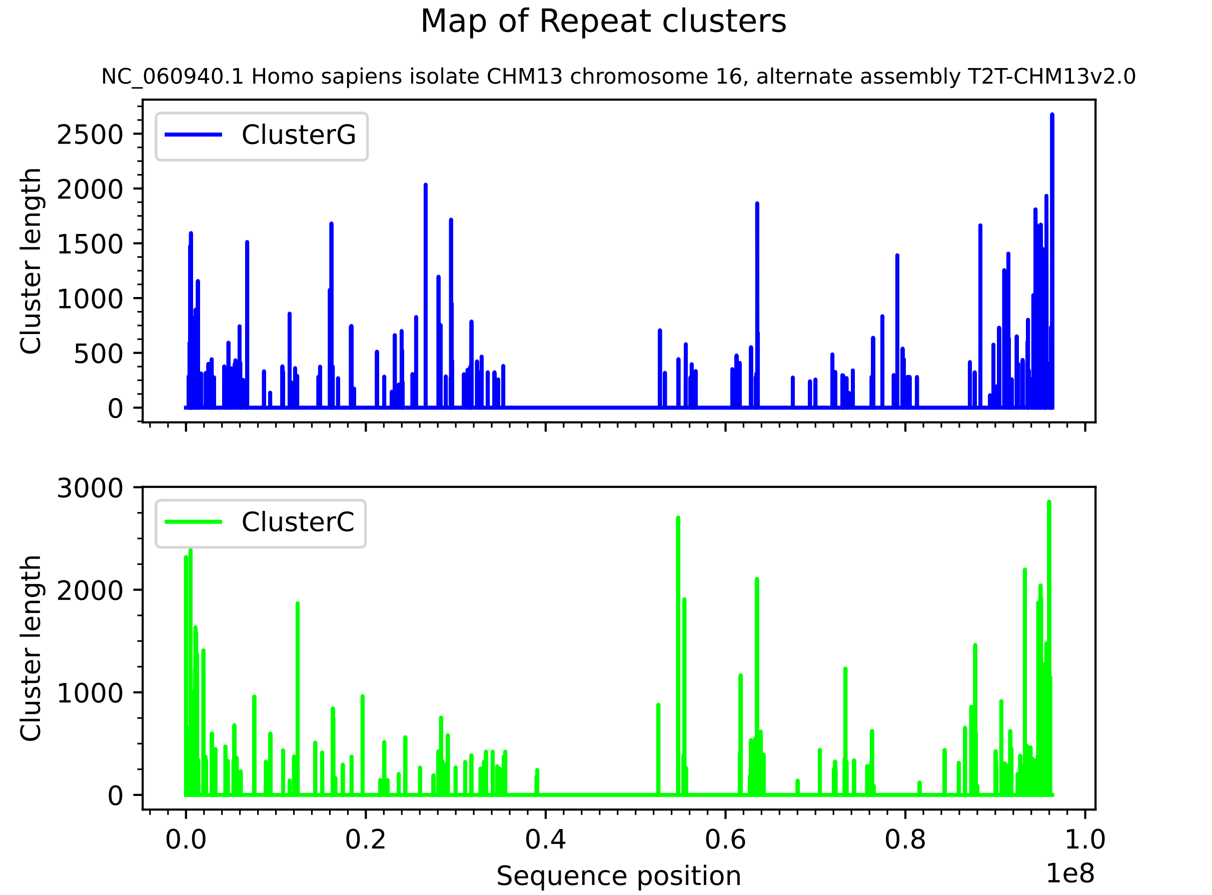

Map of Repeat clusters
NC_060940.1 Homo sapiens isolate CHM13 chromosome 16, alternate assembly T2T-CHM13v2.0
ClusterG
ClusterC
Cluster length
Cluster length
0
500
1000
1500
2000
2500
0
1000
2000
3000
0.0
0.2
0.4
0.6
0.8
1.0
1e8
Sequence position

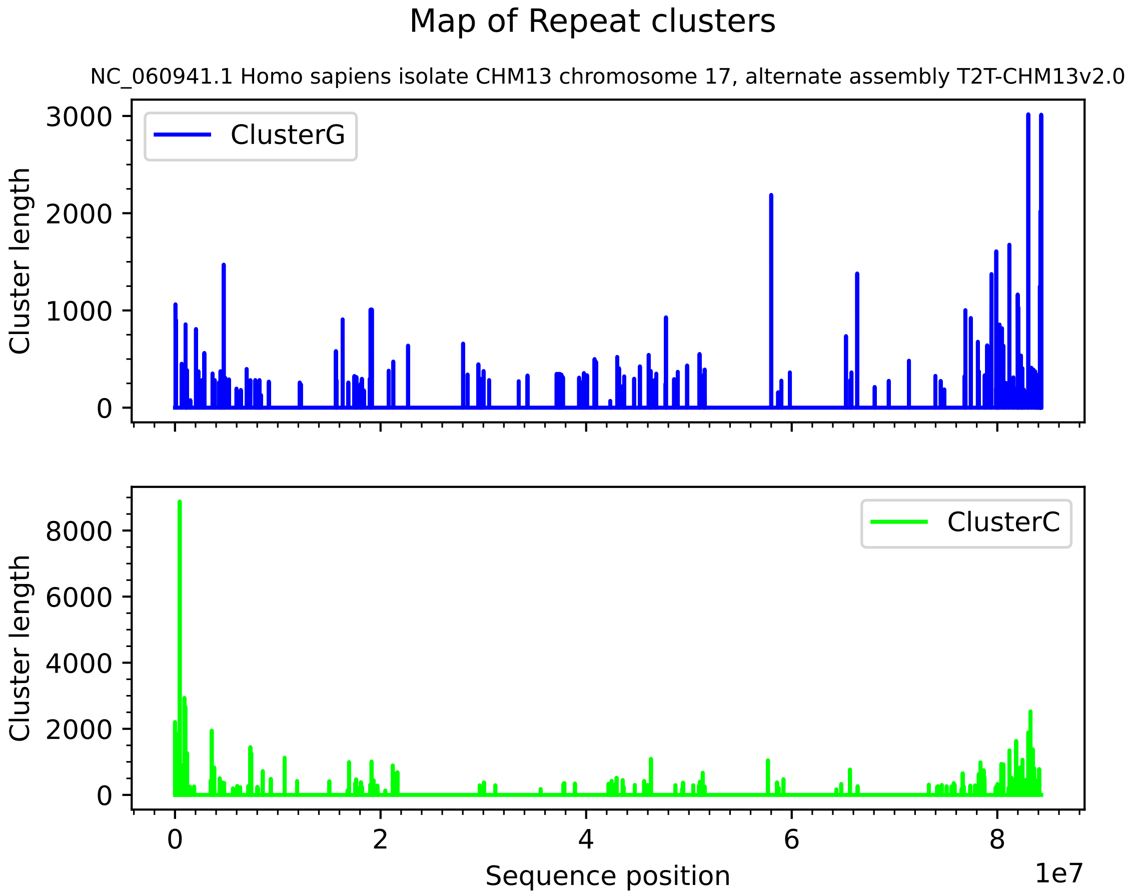

Map of Repeat clusters
NC_060941.1 Homo sapiens isolate CHM13 chromosome 17, alternate assembly T2T-CHM13v2.0
ClusterG
Cluster length
3000
2000
1000
0
ClusterC
Cluster length
8000
6000
4000
2000
0
0
2
4
6
8
1e7
Sequence position

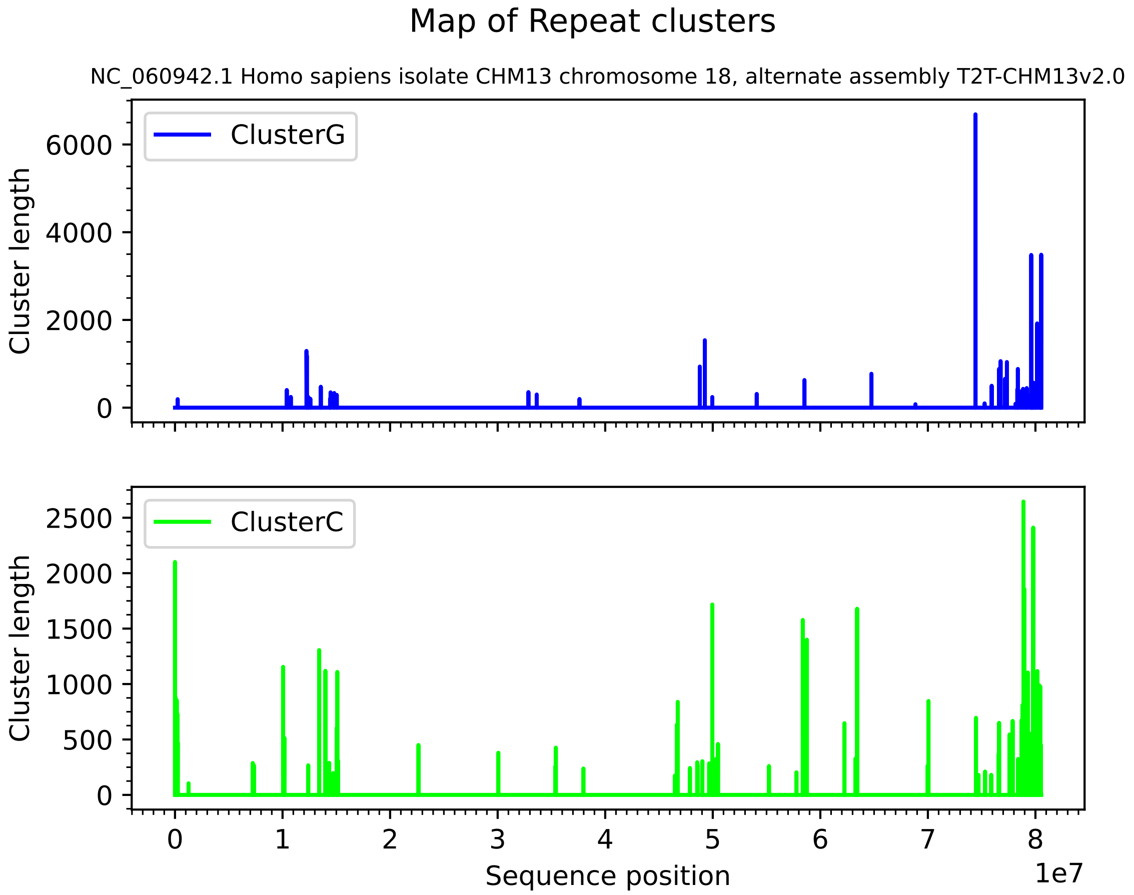

Map of Repeat clusters
NC_060942.1 Homo sapiens isolate CHM13 chromosome 18, alternate assembly T2T-CHM13v2.0
ClusterG
Cluster length
6000
4000
2000
0
ClusterC
Cluster length
2500
2000
1500
1000
500
0
0
1
2
3
4
5
6
7
8
Sequence position
1e7

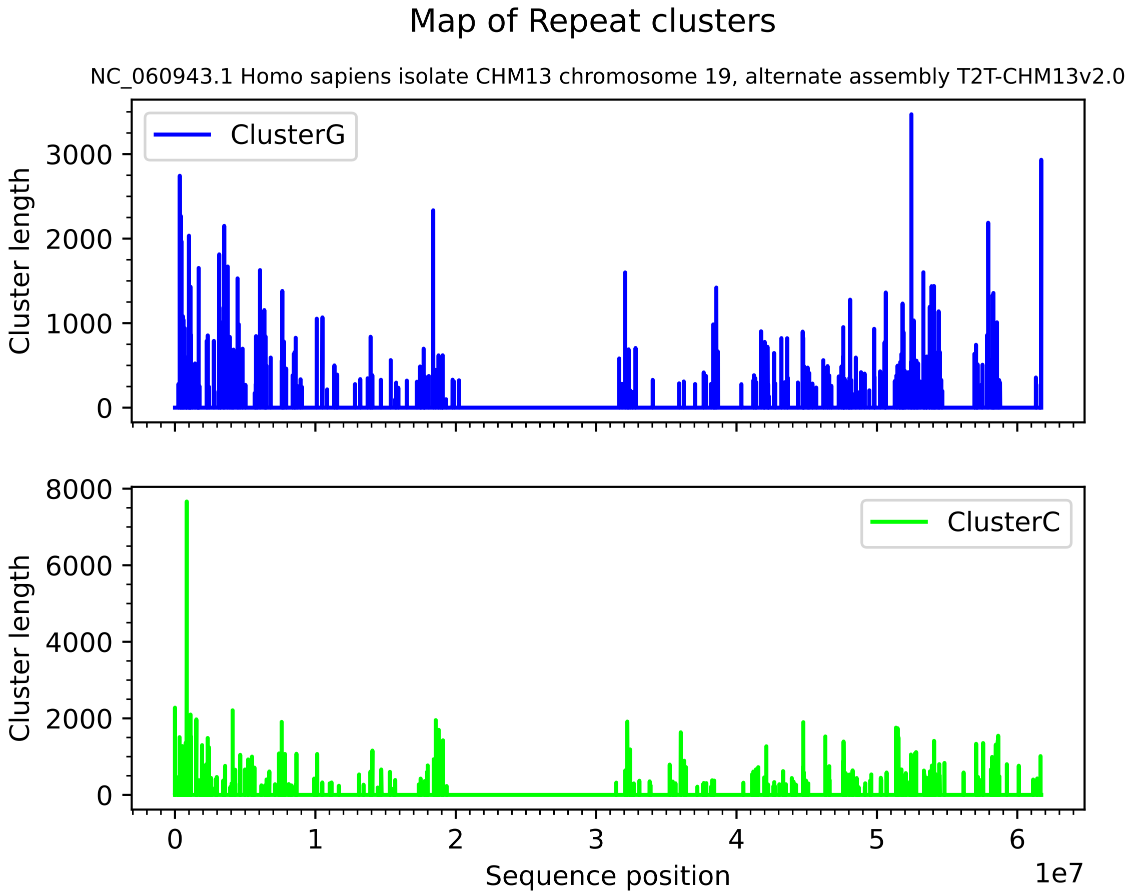

Map of Repeat clusters
NC_060943.1 Homo sapiens isolate CHM13 chromosome 19, alternate assembly T2T-CHM13v2.0
ClusterG
Cluster length
3000
2000
1000
0
ClusterC
8000
6000
4000
2000
0
0
1
2
3
4
5
6
1e7
Sequence position

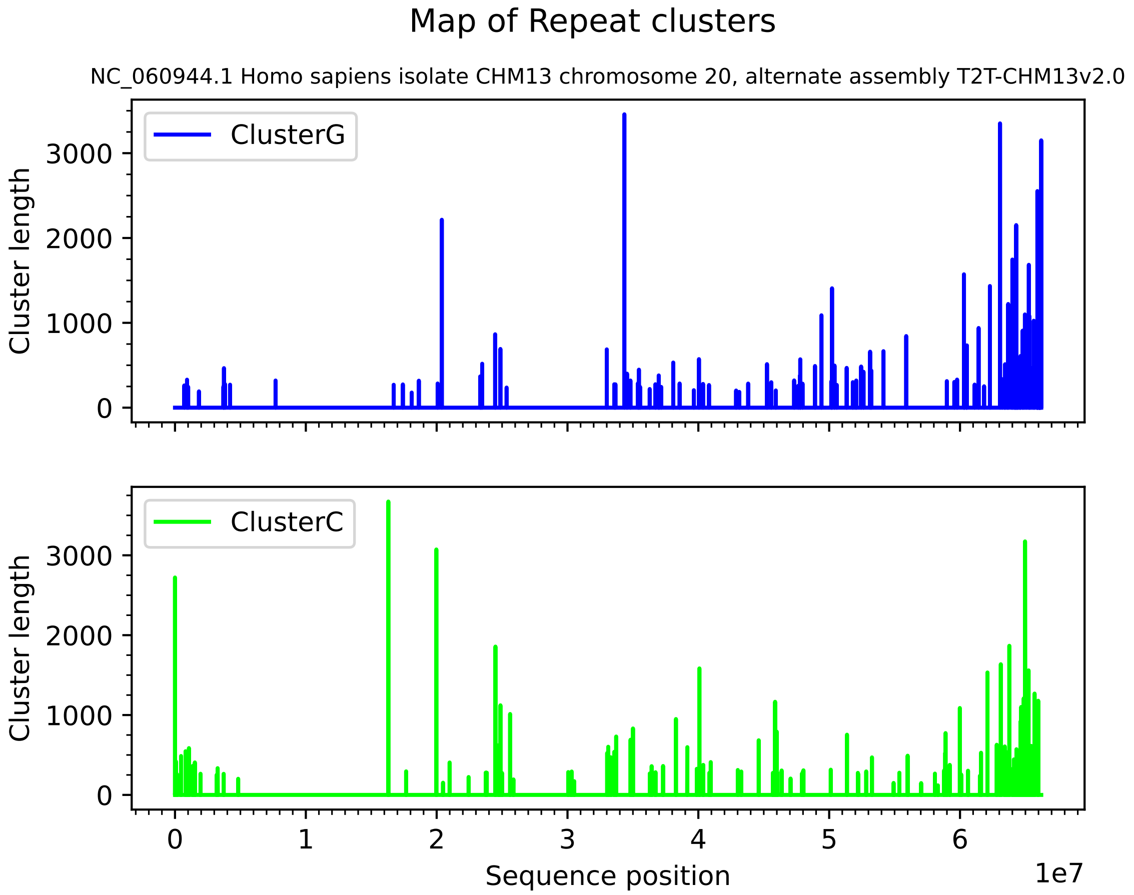
Map of Repeat clusters
NC_060944.1 Homo sapiens isolate CHM13 chromosome 20, alternate assembly T2T-CHM13v2.0
ClusterG
Cluster length
0
1000
2000
3000
ClusterC
Cluster length
0
1000
2000
3000
0
1
2
3
4
5
6
Sequence position
1e7

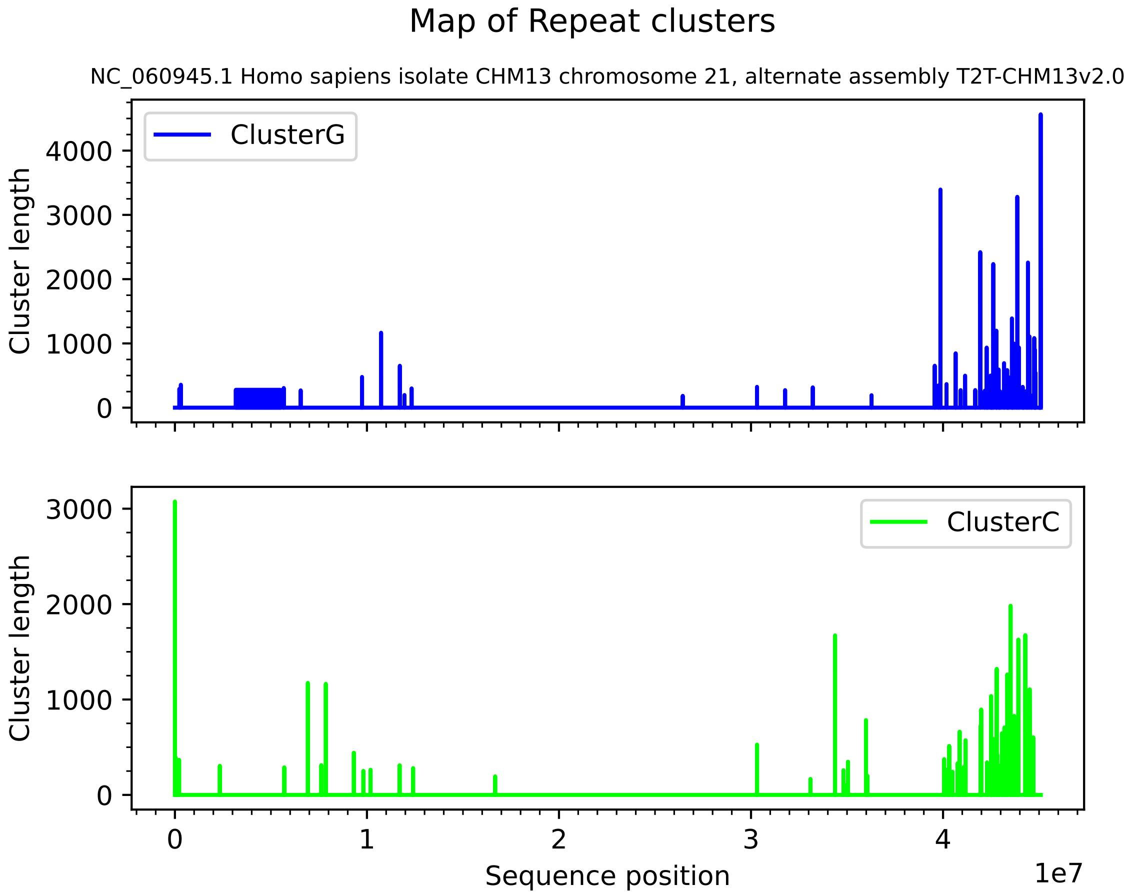

Map of Repeat clusters
NC_060945.1 Homo sapiens isolate CHM13 chromosome 21, alternate assembly T2T-CHM13v2.0
ClusterG
Cluster length
0
1000
2000
3000
4000
ClusterC
Cluster length
0
1000
2000
3000
0
1
2
3
4
Sequence position
1e7

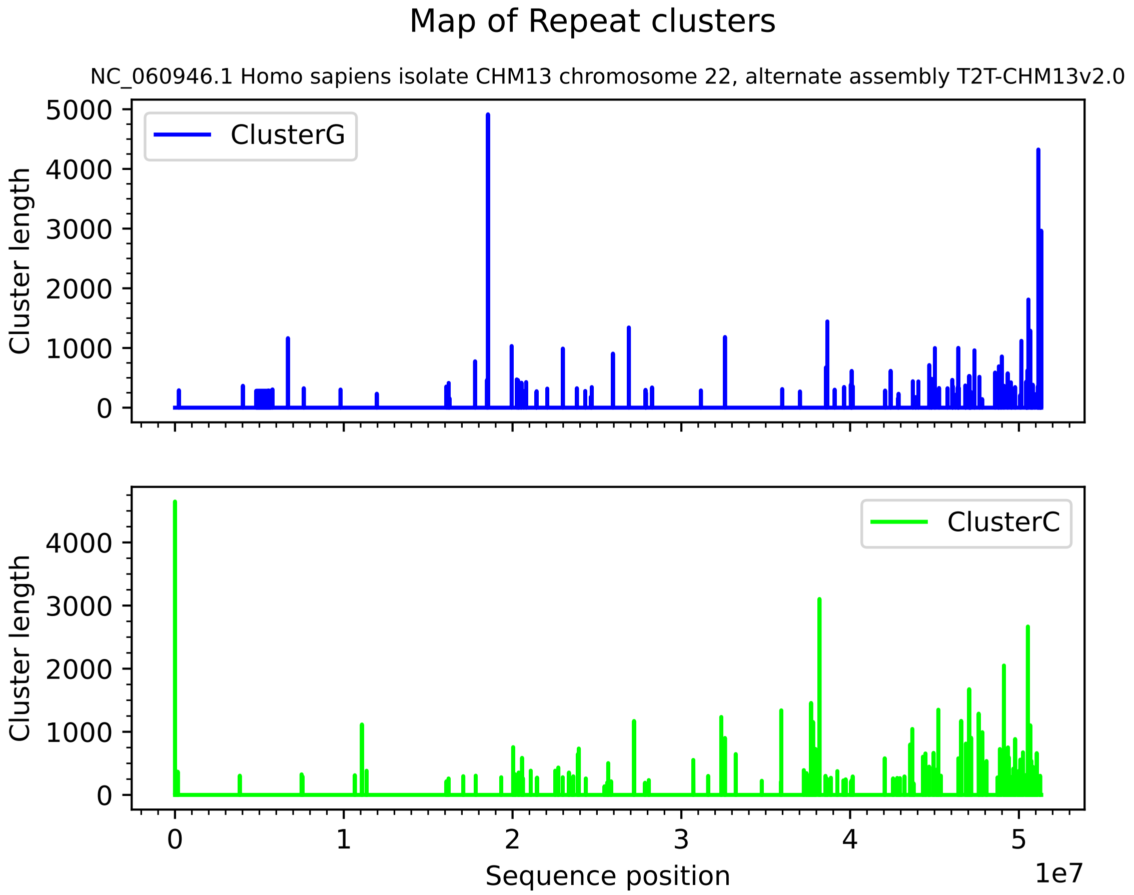

Map of Repeat clusters
NC_060946.1 Homo sapiens isolate CHM13 chromosome 22, alternate assembly T2T-CHM13v2.0
ClusterG
ClusterC
Cluster length
Cluster length
0
1000
2000
3000
4000
5000
0
1000
2000
3000
4000
0
1
2
3
4
5
Sequence position
1e7

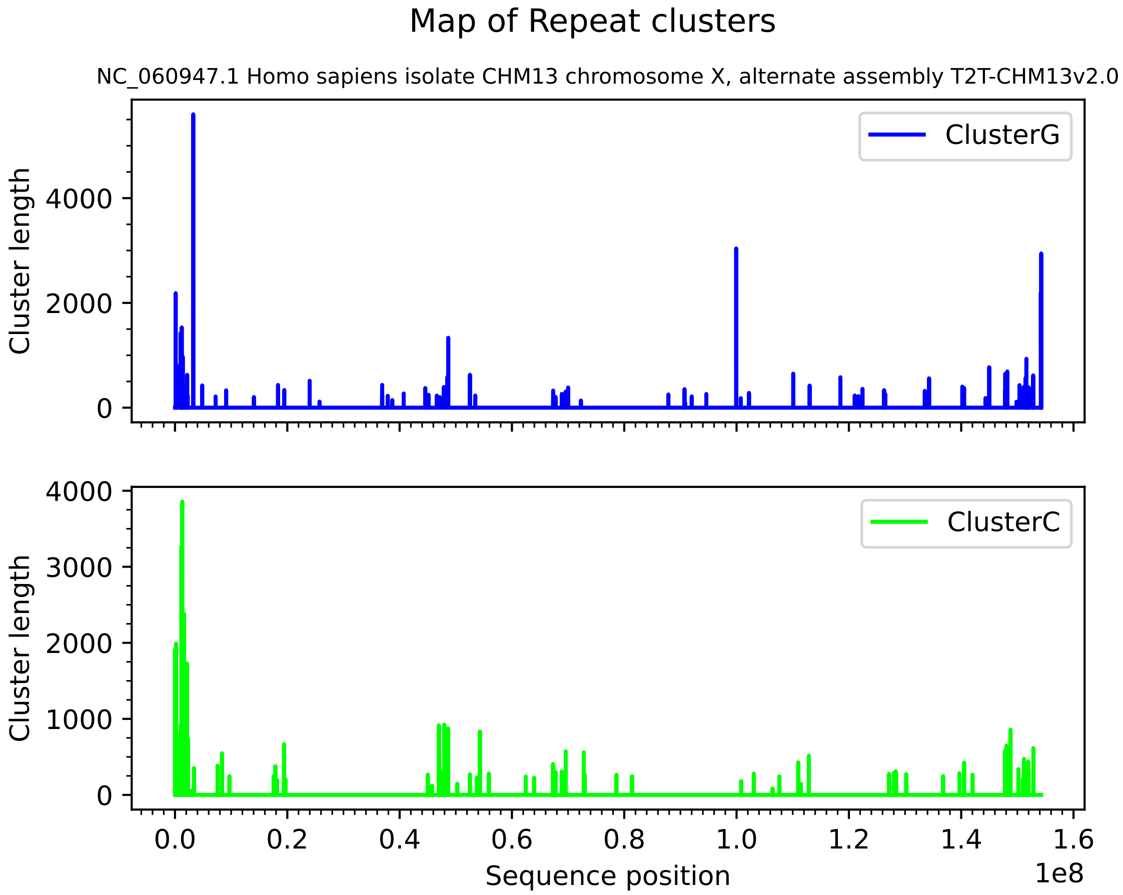

Map of Repeat clusters
NC_060947.1 Homo sapiens isolate CHM13 chromosome X, alternate assembly T2T-CHM13v2.0
ClusterG
ClusterC
Cluster length
Sequence position
1e8
0
1000
2000
3000
4000
0.0
0.2
0.4
0.6
0.8
1.0
1.2
1.4
1.6

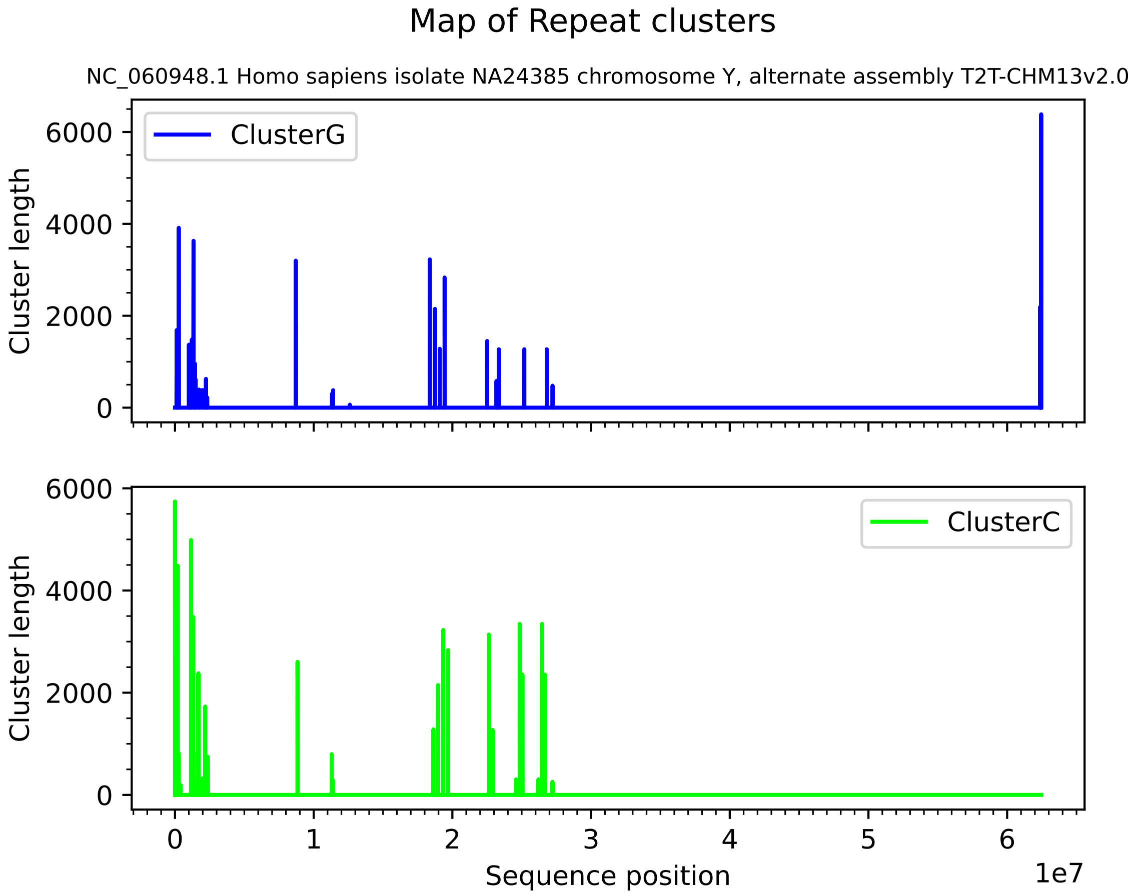
Map of Repeat clusters
NC_060948.1 Homo sapiens isolate NA24385 chromosome Y, alternate assembly T2T-CHM13v2.0
ClusterG
ClusterC
Cluster length
Cluster length
6000
4000
2000
0
6000
4000
2000
0
0
1
2
3
4
5
6
1e7
Sequence position
